# Genome-wide loss of chromatin accessibility coexists with selectively elevated RNA abundance in nutrient-starved U2OS cells

**DOI:** 10.64898/2026.10.30.763108

**Authors:** Xiyuan Qi, Sham Armouch Al Refaai, Sergei Muslinov, Konstantin I. Panov

## Abstract

**Background:** Chromatin accessibility is widely used as a proxy for regulatory potential and is often interpreted together with RNA-seq to infer gene-regulatory responses. However, the quantitative relationship between chromatin accessibility and transcript abundance may become difficult to interpret, particularly under acute cellular stress. Metabolic challenges such as nutrient deprivation are known to induce extensive chromatin remodelling and transcriptional reprogramming, raising the question of whether changes in transcript abundance are directly governed by chromatin accessibility in these contexts. Clarifying this relationship is critical for interpreting epigenomic data under stress conditions, and requires careful distinction between Tn5 accessibility, relative steady-state RNA abundance and transcriptional output.

**Results:** We integrated ATAC-seq and RNA-seq analyses in human U2OS osteosarcoma cells following 18 h of severe nutrient deprivation. We found that starvation was associated with a genome-wide reduction in accessibility and almost all significantly altered accessible regions showed reduced signal relative to actively growing cells. This decrease affected promoter- and enhancer-associated regions alike, including housekeeping-gene promoters and CTCF-motif sites.

In contrast, RNA-seq revealed a more selective change in relative steady-state RNA abundance: only a minority of expressed genes were differentially regulated, and approximately 64% of differentially expressed genes showed increased relative RNA abundance.

Changes in accessibility and transcript abundance were modestly correlated at promoters (Pearson r = 0.25) and weakly at enhancers (r = 0.10), and together explained only ∼6% of the variance in transcript abundance changes. The relationship was graded: genes with the smallest promoter losses were about four times as likely to show increased RNA abundance as genes with the largest losses, independently of baseline promoter accessibility. Within-condition analyses similarly showed only modest relationships between promoter accessibility and RNA abundance in both actively growing and starved cells.

Pathway-level analysis of mRNA abundance revealed the expected repression of biosynthetic and growth-related programmes, including ribosome biogenesis, translation and cellular growth, together with a less anticipated relative activation of stress-adaptive programmes, including extracellular matrix organisation, lipid and cholesterol metabolism, small GTPase signalling and developmental processes. For genes within these activated programmes, increased steady-state mRNA abundance was observed despite reduced ATAC-seq accessibility at associated regulatory regions.

Exon-intron analyses of RNA-seq data indicated that these changes predominantly reflect altered RNA production. Metabolic labelling of nascent RNA at selected *loci* confirmed increased transcription of genes involved in adaptive programmes despite reduced promoter and enhancer accessibility, alongside with reduced production of biosynthetic transcripts and rRNA.

**Conclusions:** These findings demonstrate that chromatin accessibility and steady-state RNA abundance are only modestly coupled during severe nutrient deprivation, against a baseline of similarly modest promoter accessibility-abundance coupling. This underscores the limited ability of ATAC-seq accessibility to predict transcript abundance under this stress condition and supports a model in which chromatin accessibility acts as a permissive rather than deterministic layer of gene regulation. The selective induction of adaptive programmes, confirmed here at the level of RNA production, shows that transcriptional activation can occur despite genome-wide loss of ATAC-seq accessibility. Because ATAC-seq reports Tn5 accessibility rather than physical chromatin compaction, we interpret compaction as a plausible mechanistic model compatible with the current observations, which will require direct orthogonal validation.

## Introduction

In eukaryotic cells, the packaging of genomic DNA into chromatin imposes a fundamental constraint on gene regulation: transcription factors, RNA polymerase, and associated co-regulators must access their target sequences within a nucleosomal landscape that is largely restrictive by default. Regions of open or accessible chromatin, where nucleosome occupancy is locally reduced, are therefore widely regarded as sites of active or poised gene regulation [1, 2]. This principle underpins modern models of transcriptional control and has driven the development of genome-wide assays that map chromatin accessibility as a proxy for regulatory potential.

Genome-wide profiling of chromatin accessibility has been greatly accelerated by ATAC-seq (Assay for Transposase-Accessible Chromatin using sequencing), which is now routinely used across diverse biological contexts (e.g. in developmental biology, immunology, cancer research, and disease modelling) [3].

A frequent and often implicit assumption in the interpretation of ATAC-seq data is that changes in chromatin accessibility reflect, and can therefore predict, changes in transcriptional activity. This coupling has been well documented in stable cell-state transitions, including differentiation and development, where chromatin remodelling accompanies durable transcriptional changes [4, 5]. Large-scale projects such as ENCODE and the Roadmap Epigenomics Project have further reinforced this view by revealing concordant accessibility and expression patterns across cell types and tissues [5, 6]. As a result, chromatin accessibility is increasingly used as a surrogate for transcriptional activity in both bulk and single-cell studies [7, 8].

However, accumulating evidence indicates that the relationship between chromatin accessibility and gene expression is more complex and context-dependent than this framework implies. Kiani et al. systematically analysed accessibility and expression changes across diverse single-factor perturbations in human cells and found that changes in chromatin accessibility were frequently discordant with transcriptional responses[9]. Similar weak or absent coupling has been reported under oxidative stress, heat stress and other perturbations, where changes in accessibility and RNA abundance often do not scale proportionally at matched genomic *loci* [10–13]. These observations suggest that chromatin accessibility provides important regulatory context but is not necessarily a simple quantitative predictor of gene-expression responses.

Notably, many previous studies examined perturbations that produced mixed gains and losses of accessibility or involved complex multi-step differentiation processes. It therefore remains unclear how accessibility-expression coupling behaves under conditions dominated by a broad directional reduction in ATAC-seq accessibility, where standard assumptions about chromatin accessibility, normalization and transcriptional interpretation may be particularly difficult to apply.

Cellular stress responses provide a particularly useful setting for examining this relationship. In *Saccharomyces cerevisiae*, entry into quiescence is accompanied by condensin-mediated chromatin compaction and genome-wide transcriptional repression[14, 15]. In *Caenorhabditis elegans*, starvation triggers a global chromatin compaction pathway that represses germline gene expression [16].

In mammalian cells, arginine deprivation and transient ischemia similarly induce widespread changes in chromatin accessibility with complex and context-dependent gene-expression outcomes [17, 18]. Conversely, stress can also produce localized increases in chromatin accessibility rather than uniform closure, as illustrated by the dramatic nucleosome loss at heat-shock *loci* in *Drosophila* and by more modest, locus-specific accessibility gains reported in heat-shocked human cells [19, 20], indicating that the chromatin response to stress is regionally heterogeneous rather than uniformly repressive. These studies indicate that stress-associated changes in chromatin structure and accessibility are widespread, but the quantitative relationship between accessibility and RNA abundance remains incompletely understood.

This gap in understanding has important practical and conceptual consequences. ATAC-seq data are increasingly used as a surrogate for regulatory or transcriptional state [7, 8, 21], yet this inference may be less reliable under stress conditions associated with broad changes in accessibility. Conceptually, it remains unclear whether stress-associated gene-expression programmes scale with local accessibility changes, or whether selected programmes can be maintained or elevated despite widespread reduction in ATAC-seq signal.

Here, we address this question by integrating ATAC-seq and RNA-seq analyses in human U2OS osteosarcoma cells subjected to severe nutrient deprivation and validate key findings by direct measurement of nascent RNA at selected *loci*. This model is associated with a genome-wide loss of ATAC-seq accessibility, providing a stringent context in which to examine how accessibility changes relate to steady-state RNA abundance. We find that accessibility-RNA abundance correlations are weak, that about 6% of genes of matched regulatory element-gene pairs show concordant directional changes, and that promoter and enhancer accessibility together explain only a small fraction of RNA-abundance variance. Pathway-level analysis further reveals coordinated repression of biosynthetic and growth-related programmes, including ribosome biogenesis, translation and cell-cycle progression, alongside selective elevation of stress-adaptive programmes, the latter showing increased steady-state mRNA abundance despite reduced ATAC-seq accessibility at associated regulatory regions.

Together, these findings highlight the limited ability of ATAC-seq accessibility alone to predict steady-state RNA-abundance responses during severe nutrient deprivation and support the need for integrative interpretation of chromatin accessibility and transcriptome data under stress conditions.

## Materials and Methods

### Cell culture and starvation

Human U2OS osteosarcoma cells (ATCC HTB-96) were maintained in McCoy 5A Medium supplemented with 10 % fetal bovine serum at 37 °C and 5 % CO_2_. Actively growing cells (AG) were collected at 70% confluency. For starvation, 70% confluent cultures were washed once with pre-warmed PBS and incubated for 18 h in serum-free PBS diluted (1:1) Low Glucose DMEM media (starved state, SS) as in our previous work [22]. Three biological replicates were collected from SS and AG conditions for ATAC-seq and two for RNA-seq analysis. Percent of viable cells was determined using Live/Dead Viability/Cytotoxicity Kit (Thermo Fisher Scientific) and cell cycle distribution was analysed by FACS of propidium iodide treated cells. Cell proliferative capacity was measured using the PrestoBlue reagent (Thermo Fisher Scientific). Equal numbers of starved and untreated cells were replated, and the metabolic signal from viable cells was recorded at 0, 24, 53, 74, and 96 hours post-replating.

### RNA Extraction, Library Preparation, and Sequencing

Total RNA was extracted (RNeasy Plus Mini Kit, Qiagen) and genomic DNA was eliminated with Ambion DNA-free DNA removal kit (Thermo Fisher Scientific). RNA was analyzed for integrity using Fragment analyser (Agilent). Libraries were prepared using KAPA RNA HyperPrep Kit with RiboErase (Roche), and barcoded libraries were pooled in equimolar ratios and sequenced using a NextSeq2000 (Illumina) at the Genomics CTU, Queen’s University Belfast (QUB). All sequencing data is available at Gene Expression Omnibus (accession number GSE327351).

Cellular poly(A)+ RNA content was quantified in AG and SS cells using two independent approaches as described in **Section 3** of Supplementary Data.

### RNA-seq data processing

Initial RNA-seq data processing, QC and postprocessing were performed by Genomics CTU (QUB) using GCTU RNA-seq pipeline v2.6.1 yielding QC Report and annotated raw reads. Briefly, STAR2.7.3a was used to align reads to the genome assembly GRCh38, SAMtools 1.17 and Picard 3.0.0 were used for post-alignment processing of BAM files such as sorting, indexing, merging, duplicate marking/removal, generating alignment statistics and RNA-specific quality metrics alongside with Qualimap 2.3.0, MultiQC 1.14 and FastQC 0.11.8. HTSeq 0.11.3 was used for read counts, and BioMart for transcripts annotation.

### RNA-seq data analysis

Differential expression analysis was performed using DESeq2 1.46, Wald test [23] yielding unfiltered, not shrunken list of DEGs. Statistical power and dispersion estimates were characterized to assess the detection sensitivity of the n=2 design. Using this list, gene-set enrichment analysis was performed using fgsea 1.32.4 [24] with curated MSigDB collections (C3:TFT, C5, Hallmark, C2:CP) [25, 26] to obtain pathway-level insights.

Genes were ranked by moderated Wald test statistics from DESeq2, which provide stable variance estimates and are recommended for gene set enrichment analysis, particularly in low-replicate design [27]. For SS vs AG comparison, fgsea was run with the following parameters: minimum gene-set size = 15, maximum gene-set size = 500, and 10,000 permutations. Enrichment was considered statistically significant at a false discovery rate (FDR) q-value ≤ 0.05.

To reduce redundancy among enriched pathways, we consolidated results into high-level biological categories. Redundancy was reduced by clustering pathways that passed an FDR threshold of ≤ 0.01 based on similarity in leading-edge gene overlap, quantified using a symmetric Cohen’s kappa coefficient matrix computed from pairwise overlaps within a defined background gene universe [28]. A weighted graph was then constructed by retaining edges with kappa values above an adaptive threshold (90th percentile of non-diagonal elements). Louvain community detection [29] was applied to identify pathway clusters. For each cluster, representative pathways (top three ranked by adjusted p-value and normalized enrichment score [NES]) were selected. Category names were derived via meta-enrichment of aggregated leading-edge genes using GO-BP, Reactome pathways, and MSigDB Hallmark sets, followed by manual curation. Category-level summaries were calculated using a weighted NES (wNES) and a combined Stouffer’s Z-score.

wNES were calculated for each biological category by computing a significance-weighted average of individual pathway NES values, where weights were defined as -log_10_(padj). This approach adapts the NES normalization concept [25] with meta-analytic weighting principles to enable category-level enrichment summarization.

To complement this effect size measure with formal statistical inference, Stouffer’s Z-score method was applied to combine pathway-level p-values within each category, providing a validated framework for assessing overall category significance. The Z-scores were capped at 10 for individual pathways to prevent extreme values from dominating visualizations while preserving biological interpretability. Combined Stouffer Z was calculated as: Stouffer_Z = ∑ (Z_i_) / √n .

This framework reduced functional redundancy while preserving biological interpretability, analogous to meta-analysis strategies widely used in transcriptomics [24, 25].

### Transcription factor and chromatin-state enrichment analysis

Candidate transcriptional regulators and chromatin-state associations were identified by over-representation analysis using GSEApy [30] and Enrichr [31, 32]. Four Enrichr libraries were queried: ChEA_2022, ENCODE_TF_ChIP-seq_2015, TF_Perturbations_Followed_by_Expression, and ENCODE_Histone_Modifications_2015 [6, 33]. Analyses were performed separately for up- and downregulated DEGs in SS versus AG cells (padj ≤ 0.05, |LFC| ≥ 1), and for leading-edge genes from the four upregulated adaptive programmes: ECM Remodelling, Lipid/Cholesterol Metabolism, Small GTPase Signalling, and Developmental/Differentiation (**Table S7**). The full set of expressed genes detected by RNA-seq was used as the statistical background. Enrichment was assessed using Fisher’s exact test with Benjamini-Hochberg correction, and terms with FDR ≤ 0.05 were retained. Where multiple terms represented the same regulator, results were summarised at regulator level by retaining the smallest adjusted p-value and calculating the mean odds ratio across contributing terms. These analyses were used to nominate candidate regulatory associations and do not directly measure transcription factor occupancy, histone-modification abundance, or regulator activity in starved U2OS cells.

### Exon-intron concordance and EISA analysis

Exonic and gene-body genomic intervals for protein-coding genes were defined from the GENCODE v33 basic annotation used to generate STAR indexes. Exonic intervals were derived as the strand-aware union of all exon records per gene, merged into non-overlapping intervals using GenomicRanges::reduce(); gene-body intervals corresponded to the full genomic span of each gene. Exonic and gene-body read counts were generated independently using featureCounts (Rsubread v2.20.0) [34], restricted to protein-coding genes. Intronic counts were estimated per gene as the arithmetic difference between gene-body and exon counts; genes yielding negative derived intronic counts in any sample were excluded. Genes were retained for analysis if intronic length ≥ 500 bp, mean raw exon count ≥ 10, mean raw intronic count ≥ 5, and intronic signal (count ≥ 1) detected in at least two samples.

Exonic and intronic count matrices were analysed independently using DESeq2 v1.46.0 [23] (design ∼ condition, SS vs AG), with apeglm-shrunken LFCs used for all concordance statistics. Genes were classified as UP, DOWN, or UNCHANGED independently in each compartment using thresholds of |LFC| ≥ 0.5 and padj ≤ 0.05. Directional concordance among genes with significant directional calls in both compartments was tested by permutation against a 10,000-iteration label-shuffled null.

The formal EISA interaction model [35] was implemented with eisaR v1.18.0 (runEISA, method = ‘Gaidatzis2015’), with AG as the reference level. Exonic and intronic count matrices were first filtered for intron length (≥ 500 bp), mean raw exonic count (≥ 10), mean raw intronic count (≥ 5) and intronic reads detected in at least two samples. The Gaidatzis2015 setting reproduces the published method: exonic and intronic counts are normalised independently (edgeR::calcNormFactors, sizeFactor = ‘individual’), genes are retained if log2(x + 8) > 5 for both exonic and intronic counts (10,881 genes), and the condition-by-region interaction (Dex - Din) is tested with a likelihood-ratio test (edgeR glmFit and glmLRT) in a model without sample-specific terms (modelSamples = FALSE).

Genes were classified as post-transcriptional stabilisation (Dex - Din > +0.5, FDR < 0.05), destabilisation (Dex - Din < -0.5, FDR < 0.05), significant with small effect size (|Dex - Din| < 0.5, FDR < 0.05) or concordant/background (FDR ≥ 0.05). Genes removed by the eisaR expression filter were not classified. A more stringent threshold (|Dex - Din| ≥ 1.0, FDR < 0.05) was additionally applied to define high-confidence post-transcriptional candidates. Sign convention follows [35]: because the intronic response approximates the change in transcriptional output, a positive interaction term indicates that mature transcript abundance changed more than production and is interpreted as post-transcriptional stabilisation, whereas a negative term indicates destabilisation or accelerated decay.

Two robustness analyses were performed. First, genes were stratified into quartiles by intron count, and the proportion classified as high-confidence destabilisation candidates (Dex – Din <= -1.0, FDR < 0.05) was compared across quartiles by chi-squared test and by logistic regression with baseline exonic expression as a covariate. The destabilisation class was chosen for this test because intron retention inflates intronic counts, raises Din and therefore drives the interaction term negative; it is this class, and not the stabilisation class, that is susceptible to a retention artefact. Second, concordance statistics (Pearson r, same-sign fraction, directional-only concordance) were recomputed after excluding the top intron-count quartile and, separately, all genes with ≥ 20 introns.

### Metabolic labelling and capture of nascent RNA

Nascent RNA was labelled with 5-ethynyluridine (5-EU), biotinylated by copper-catalysed azide-alkyne cycloaddition and captured on streptavidin beads, adapting Palozola et al. [36, 37]. Equal numbers (6 x10^6^ cells) of U2OS cells under actively growing (AG) conditions or following 18 h of starvation (SS) were processed in parallel in three independent experiments. 100× 5-EU stock in PBS was added directly to existing culture medium (10μl of per 1ml of medium) yielding final 1 mM 5-EU concentration. Cells were incubated for 30 min at 37 °C in 5% CO_2_, then were washed with cold PBS and lysed using PureLink RNA Mini Kit (Thermo Fisher Scientific Scientific) lysis buffer. One no-5-EU control per condition was processed per experiment.

Each lysate received 3.5 × 10^10^ copies of 5-EU-labelled spike-in RNA (Sp) produced as described in **Section 5** of Supplementary Data. RNA was purified with the PureLink RNA Mini Kit, treated with the DNA-free Kit (Thermo Fisher Scientific Scientific) and quantified using Quant-iT RNA Assay Kit, Broad Range (Thermo Fisher Scientific Scientific). RNA was split in to total (T) and nascent (NS) branches (∼1:10). The first was reverse transcribed directly and the second used for nascent capture. Biotinylation proceeded for 30 min at room temperature in 50 mM MOPS (pH 7.0), 1 mM azide-PEG3-biotin, 2 mM CuSO_4_, 10 mM sodium ascorbate and 12 mM THPTA. After EDTA addition (15mM final concentration) and repurification (PureLink RNA Mini Kit), RNA was denatured at 68°C for 5 min and incubated with 40 µL Dynabeads MyOne Streptavidin T1 (Thermo Fisher Scientific Scientific) for 30 min. Washed bead-bound RNA was split in 5 tubes and reverse transcribed using Transcriptor First Strand cDNA Synthesis Kit (Roche), random hexamers in 20 µL reactions. Reaction conditions were: 65 °C, 10 min; 25 °C, 10 min; 55 °C, 30 min; 85 °C, 5 min. On completion reactions were pooled. Total RNA underwent identical reverse transcription in two 20 µL reactions, which were pooled. Final cDNA volumes were 40 µL (total) and 100 µL (nascent).

### Quantitative PCR

LightCycler 480 qPCR (Roche) used 10 µL reactions containing 5 µL LightCycler 480 SYBR Green I Master, 500 nM each primer and 2 µL diluted cDNA: 95 °C for 5 min; 40 cycles of 95 °C, 60 °C and 72 °C for 10 s each; then melting curve analysis. Primers were designed using NCBI Primer-BLAST and each primer pair produced a single melting peak (Supplementary **Table S16**).

Total and nascent cDNA dilutions were 1:5/1:25/1:125 and 1:5/1:10/1:25, respectively, with one well per gene per dilution. Linearised plasmid standards (separate for 28S and Sp) comprised six concentrations (3.2 × 10^5^-2.0 × 10^9^ copies/reaction; two or three wells each). No-template (NTC), no-reverse-transcription (no-RT) and no-5-EU controls were included.

### qPCR data analysis

Acceptance and exclusion criteria were specified before analysis. Reporting followed MIQE 2.0 [38]. Cq values (reported as Cp) were determined by second-derivative maximum in LightCycler 480 software v1.5.0. Measurements required Cq ≤ 35 and at least five cycles before same-plate NTCs and, for total RNA, no-RT controls adjusted by adding log_2_(DF_sample_/DF_noRT_), where DF is dilution factor. Values were not altered or imputed. Standard-curve filtering removed end levels deviating by > 1 cycle from the remaining-level fit and wells with leave-one-out residuals > 0.5 cycle. Curves required ≥ 3 levels spanning ≥ 2 orders of magnitude, R^2^ ≥ 0.98 and slopes -3.6 to -3.1. A curve failing the slope criterion was refitted with the pooled slope of the other plates for that assay, and its intercept was re-estimated from its own standards for that fixed slope. Gene assays with an amplification efficiency outside 85-115% were excluded. Sample wells outside calibration or differing by > 0.5 cycle after dilution correction were excluded. Copies/reaction were multiplied by DF × V/2 (V, final cDNA volume), divided by branch specific RNA input (µg) and averaged on the log scale. Fraction-specific Sp recoveries (R_T_, R_NS_) were relative to the expected Sp copies/µg (3.5 × 10^10^ divided by the RNA yield of the sample); capture efficiency was R_NS_/R_T_.

Gene efficiencies were estimated across all six total-RNA dilution series, regressing Cq on - log₁₀ DF with common slope b and sample-specific intercepts: E = 10^-1/b^ - 1. Assays outside 85-115% were excluded. Undiluted Cq was C_q0_ = C_q_ + b log_10_ DF. Nascent 1:5 points exceeding the mean undiluted Cq of more dilute points by > 0.5 cycle were excluded for inhibition. Remaining points within 0.5 cycle of their mean were averaged after removing at most one discordant point. Relative quantities per µg were calculated as log_2_ Q = -Cq_o_ log_2_ (1 + E) + log_2_ (V/2m), where Cq₀ is the mean undiluted Cq, V the final cDNA volume (µL) and m the branch-specific RNA input (µg); the same efficiency applied to both fractions.

Total quantities were normalised to 28S copies/µg measured in the same cDNA. Nascent quantities were divided by R_NS_; log₂ nascent-to-total ratios were log_2_ ((Q_NS_/R_NS_)/(Q_T_/R_T_)), using fraction-specific target quantities Q_T_ and Q_NS_. Sensitivity analyses used unnormalised total RNA, Sp-normalised total RNA (relative to Sp copies in the same cDNA), uncorrected nascent RNA and uncorrected nascent-to-total ratios.

Combined dilution estimates were analysed as three paired biological replicates using limma [39] on log₂ values (design ∼ experiment + condition; empirical Bayes moderation with trend = FALSE and robust = FALSE; Benjamini-Hochberg correction within each analysis; q < 0.05). 95% confidence intervals were obtained using topTable. Concordant genes changed significantly in the same direction in both fractions without significant ratio changes; divergent genes had significant ratio changes. 28S and Sp endpoints were compared using two-sided paired t-tests on log₂ values. No a priori power calculation was performed. qPCR LFCs were compared with gene-level and intronic DESeq2 LFCs and with EISA Dex - Din (sign-reversed to match the nascent-to-total ratio) by Spearman correlation, with 95% confidence intervals from 10,000 bootstrap resamples of genes. Analyses used R 4.3.3 with limma 3.58.1 and Python 3.13 (NumPy, SciPy and pandas).

### ATAC-seq library preparation and sequencing

Libraries were produced essentially as described by Corces et al. and Grandi et al. using 50,000 nuclei per single library and the same batch of Tn5 for all libraries[40, 41].

Pre-sequencing library quality was assessed by the trace variation (1-6000 bp) with the Agilent 5200 Fragment Analyzer. Libraries showing acceptable pre-sequencing quality were pooled and processed for paired-end sequencing using a NextSeq2000 (Illumina) at the Genomics CTU, QUB.

### ATAC-seq data processing

Initial ATAC-seq data processing, QC and postprocessing were performed by Genomics CTU (QUB) using nf-core/atacseq analysis pipeline developed by Nextflow [42], but processing was terminated before peak calling stage. Briefly, reads were trimmed (cutadapt 3.4), aligned (bwa 0.7.17-r1188), filtered to remove mitochondrial reads, PCR-duplicates and low-quality reads yielding sorted and indexed BAM/BAI files and QC Report. SAMtools 1.17 and Picard 3.0.0 were used for post-alignment processing of BAM files.

To assess library quality we performed analysis of the mapped paired-end fragment-size distributions and the fractions of nucleosome-free (NFR; ≤100 bp), mononucleosome (147-247 bp), and dinucleosome (315-473 bp) fragments. The data were derived from the Picard CollectInsertSizeMetrics output of the nf-core/atacseq pipeline (v2.1.2), computed on post-trimmed, post-aligned, post-deduplicated, and mitochondrially-cleaned BAM files. Per-base-pair histograms were exported from MultiQC v1.13, and the proportion of fragments falling within each size window was computed by integrating the histogram over the window and expressing the result as a percentage of total mapped paired-end fragments per sample. Size windows correspond to genomic insert sizes after adapter trimming and follow the conventions of Buenrostro et al. [43] and Corces et al. [40]: the NFR window (≤100 bp) captures fragments protected only by a single (or no) nucleosome contact; the mononucleosome window (147-247 bp) brackets a single 147 bp nucleosomal footprint plus 0-100 bp of flanking linker; and the dinucleosome window (315-473 bp) brackets two nucleosomal footprints (2 × 147 = 294 bp) plus 21-179 bp of inter-nucleosomal linker. Pre-sequencing library quality was independently assessed by capillary electrophoresis on an Agilent 5200 Fragment Analyzer using the DNF-474 High Sensitivity NGS Fragment kit (1-6000 bp). Note that the Fragment Analyzer measures full library molecules including ∼130 bp of sequencing adapter (Tn5 mosaic ends + i5/i7 indices + P5/P7 sequences) and is therefore not directly comparable in absolute scale to the mapped genomic insert sizes.

Subsequent peak calling and reproducibility analysis followed the ENCODE ATAC-seq pipeline [44]. Read 5′ ends were shifted by +4 bp (plus strand) and −5 bp (minus strand) to mark Tn5 insertion sites, and peaks were called with MACS2 [45] (-p 0.01 --nomodel --shift - 75 --extsize 150 --keep-dup all --call-summits) on individual biological replicates, pooled replicates, and self- and pooled pseudoreplicates generated by randomly partitioning reads; the 500,000 peaks with the lowest p-values were retained. Reproducible peaks were identified with IDR [46] (--rank p.value --use-best-multisummit-IDR, with the pooled peaks as the reference peak list), retaining peaks with a global IDR ≤ 0.05 after removal of ENCODE blacklist regions (hg38, v2) [47] and non-standard chromosomes. For each condition the optimal peak set was the larger of the best true-replicate set (Nt) and the pooled-pseudoreplicate set (Np), and rescue and self-consistency ratios were calculated as in ENCODE (Table S2). IDR analysis was performed separately for AG and SS conditions.

A genome-wide consensus peak set was generated from the condition-specific IDR optimal peaks: peaks shorter than 50 bp or longer than 2,000 bp and peaks overlapping the ENCODE hg38 blacklist (v2) were removed, peaks from the two conditions within 50 bp of each other were merged using GenomicRanges [48], and every peak present in at least one condition was retained.

Fragments were quantified across the consensus peak set using featureCounts [34], counting each properly paired fragment once. Library size was defined as the total number of mapped fragments per library after removal of mitochondrial, duplicate and low-quality reads.

### ATAC-seq data analysis

Differential accessibility between starved (SS) and actively growing (AG) cells was analysed with DESeq2 [23] (Wald test). Peaks with a mean of at least 20 fragments in either condition were tested. Counts were normalised to library size (total mapped fragments per library), the default method in DiffBind [49] to preserve changes common to all peaks which would otherwise be removed [50].

Differentially accessible peaks (DAPs) were defined as padj ≤ 0.05 and |LFC| ≥ 1, the results at a relaxed threshold of |LFC| ≥ 0.58 (1.5-fold) are given in Supplementary **Table S20.** Calls were based on unshrunken LFCs and LFCs shrunken with apeglm [51] were used for ranking and for integration with RNA-seq. As a secondary analysis, counts were normalised to fragments in consensus peaks, which expresses each peak’s share of the total peak-associated signal (Supplementary **Table S3**).

To test whether the change in accessibility was global, two reference classes of sites were analysed separately: promoter peaks of housekeeping genes [52] and distal CTCF-motif sites, defined as non-promoter peaks with a match to the CTCF position weight matrix (JASPAR MA0139.1 [53], score ≥ 51% of the maximum, corresponding to p ≤ 10^−5^) within 100 bp of the peak centre on either strand. The contribution of non-specific background was estimated for each condition as the number of fragments outside consensus peaks per base pair of non-peak genome, scaled to the width of each peak plus 150 bp; a background-corrected global LFC was calculated from condition means after subtraction of this expectation. Peak-centred signal profiles and summit, flank and background signal were computed from coverage scaled to counts per million mapped fragments (Supplementary **Figure S7**, **Table S12**; Supplementary Data, Section 1.6).

Promoter-associated peaks were defined as consensus peaks overlapping a promoter window of any annotated transcript, extending 2000 bp upstream and 500 bp downstream of the transcription start site (TSS), using TxDb.Hsapiens.UCSC.hg38.knownGene 3.20.0. Asymmetric promoter windows are employed in ATAC-seq analyses to capture the directional polarity of metazoan promoter architecture, with core regulatory elements and transcription factor binding sites predominantly enriched upstream of the TSS while minimising overlap with downstream gene bodies exhibiting distinct elongation-associated chromatin patterns [7, 54]. Each promoter-associated peak was assigned to the gene whose transcript TSS was closest to the peak centre.

Non-promoter peaks were queried against the GeneHancer database v5.25 [55] to identify peaks associated with predicted enhancers. The GeneHancer “double elite” subset was used, defined as high-confidence regulatory elements where both the enhancer itself is supported by evidence from two or more independent sources (elite enhancer) and the enhancer-gene link is supported by two or more independent association methods (elite association). Peaks whose centre lay within such an enhancer were classified as enhancer-associated, and all remaining peaks as other. Enhancer-gene associations were restricted to the top three ranked target genes per peak, creating a many-to-many relationship (multiple peaks ↔ multiple genes). Despite this stringent filtering, computational enhancer-gene predictions remain imperfect and may include false-positive and false-negative associations.

ChIPseeker [56] was used for genomic feature annotation of all consensus peaks.

### Integration of RNA-seq and ATAC-seq data

RNA-seq and ATAC-seq results were matched at the gene level, following the approach of Kiani et al. [9]. Genes with a DESeq2 baseMean of at least 10 were included. For each gene, the promoter value was the median LFC of its promoter-associated peaks, and the enhancer value the GeneHancer-score-weighted mean LFC of its linked enhancer peaks. LFCs shrunken with apeglm were used for all continuous analyses in both datasets. For categorical analyses, a gene’s promoter (or enhancer) was classified as reduced if at least one of its peaks showed a significant loss and none a significant gain, as increased in the converse case, and as unchanged if none of its peaks changed significantly (padj ≤ 0.05 and |LFC| ≥ 1, as for RNA-seq; |LFC| ≥ 0.58 in Supplementary **Table S20**).

Associations between accessibility and RNA changes were quantified with Pearson and Spearman correlations across all matched genes, without filtering on significance; 95% confidence intervals were obtained from 10,000 bootstrap resamples of genes, and promoter and enhancer correlations were compared by paired bootstrap in genes with both values. Directional agreement was assessed with gene-level contingency tables and Fisher’s exact test (odds ratios with 95% confidence intervals). The relative contributions of promoter and enhancer changes were estimated by multivariable linear regression of RNA LFC on promoter and enhancer LFCs, with and without baseline promoter accessibility, defined as log2(median normalised fragments + 1) of the gene’s promoter peaks in AG cells.

To test whether RNA changes depend on the size of the accessibility change, genes were divided into quintiles of promoter LFC. Trends across quintiles were tested with Spearman correlation (RNA LFC) and the Cochran-Armitage test (proportion of genes with increased or decreased RNA), and adjusted for baseline promoter accessibility by linear and logistic regression with quintile and baseline accessibility as covariates. Within-condition relationships between promoter accessibility and RNA abundance were assessed by correlating log2(normalised mean + 1) values in AG and in SS cells. For the leading-edge genes of enriched programmes (Supplementary **Table S7**), promoter LFCs were compared with those of all other genes by the Wilcoxon rank-sum test and by linear regression adjusted for baseline promoter accessibility, with Benjamini-Hochberg correction.

Correlations restricted to genes with significant changes in both assays, and a list of high-confidence concordant genes (padj ≤ 0.01 and |LFC| ≥ 2 in both assays; Table S6), are reported as secondary analyses only, because selection on significance inflates the apparent coupling.

## Results

### Experimental design, and ATAC-seq quality assessment

To assess the impact of severe nutrient and energy deprivation (hereafter referred to as starvation) on chromatin accessibility, we performed ATAC-seq analysis on starved (SS) and actively growing (AG) U2OS cells. Starvation resulted in a moderate accumulation of cells in the G1 phase of the cell cycle (Supplementary **Figure S1A-C**), consistent with a moderate starvation-associated shift in cell-cycle distribution. The evaluation of the restoration of proliferative capacity after nutrient replenishment showed that our starvation protocol did not irreversibly impair proliferative competence of U2OS cells (Supplementary **Figure S1D**). Cell viability was measured immediately prior to library preparation and exceeded 95%.

Peaks were called using MACS2, and reproducible peaks were identified using the ENCODE IDR framework (threshold ≤ 0.05), yielding 137,830 optimal peaks in AG and 52,842 in SS cells (Supplementary **Table S2**). Merging the two sets gave a genome-wide consensus set of 88,589 peaks with a median width of 511 bp (**Figure 1B**).

**Figure 1.**
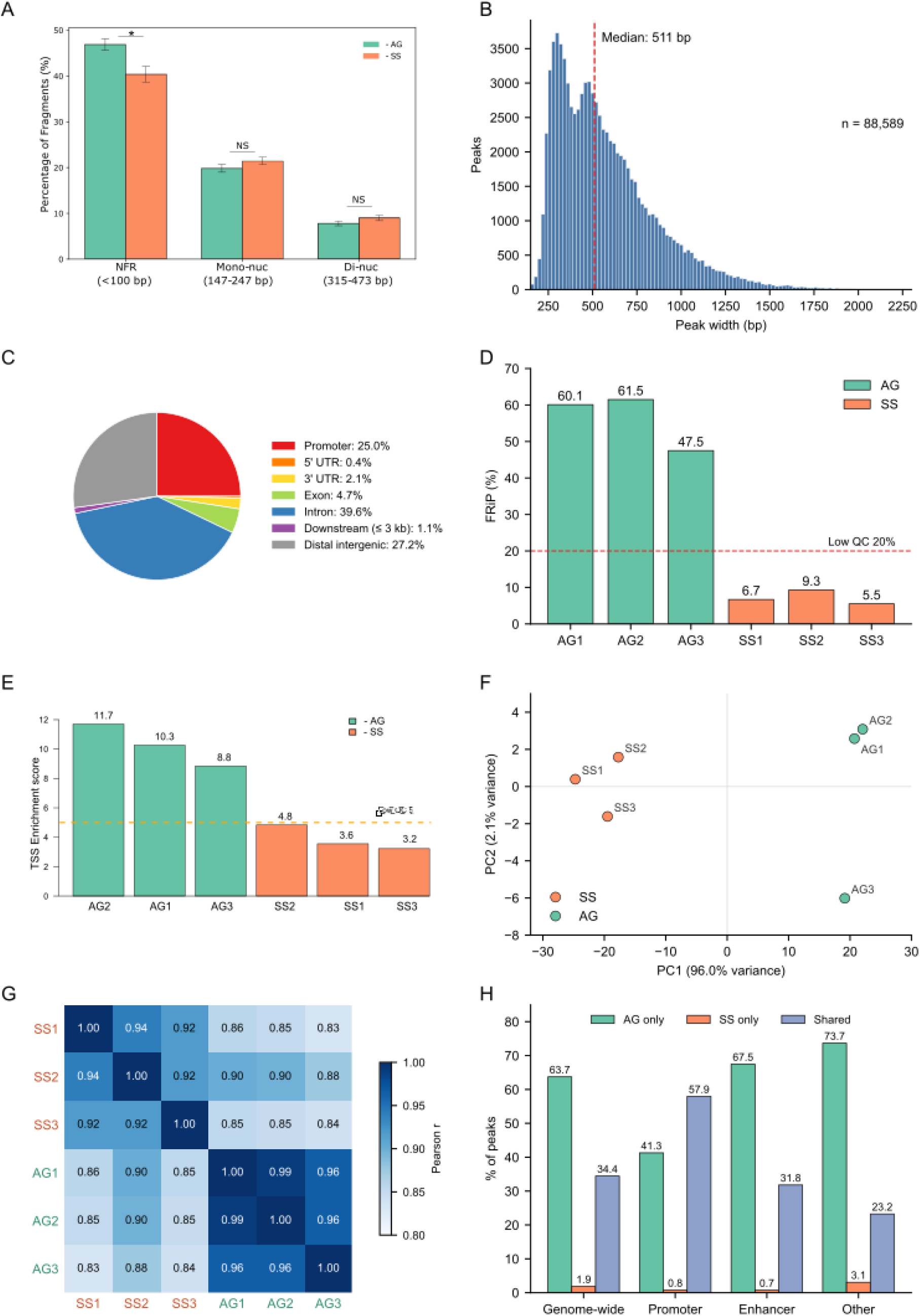
ATAC-seq quality assessment and global chromatin accessibility landscape in starved versus actively growing U2OS cells. (A) Quantification of fragments in nucleosome-free regions (NFR, ≤100 bp), mononucleosome (147-247 bp) and dinucleosome (315-473 bp) regions for actively growing (AG; green bars) and starved (SS; orange bars) samples (n = 3). Statistical significance assessed by two-tailed unpaired t-test; *p ≤ 0.05, NS = not significant. (B) Distribution of peak widths across the genome-wide consensus peak set (n = 88,589 peaks from the IDR optimal peak sets of AG and SS). Median peak width = 511 bp (red dashed line). (C) Genomic annotation of consensus peaks using ChIPseeker: promoter (25.0%), intron (39.6%), distal intergenic (27.2%), exon (4.7%), 3′ UTR (2.1%), downstream (≤3 kb; 1.1%) and 5′ UTR (0.4%). (D) Fraction of reads in peaks (FRiP) for individual libraries from AG (AG1-AG3; green bars) and SS (SS1-SS3; orange bars), calculated for the IDR optimal peak set of each sample’s own condition. The red dashed line indicates the ENCODE-recommended QC threshold of 20%. (E) Transcription start site (TSS) enrichment scores for individual libraries from AG (green bars) and SS (orange bars). The orange dashed line indicates the ENCODE-recommended QC threshold of 5.0. (F) Principal component analysis (PCA) of ATAC-seq peak accessibility (variance-stabilised counts, 500 most variable consensus peaks). PC1 (96.0% of variance) separates AG (green) from SS (orange) samples; PC2 (2.1%) captures within-condition variability. (G) Pearson correlation matrix of variance-stabilised counts across all 88,589 consensus peaks. Colour scale from 0.80 (white) to 1.00 (dark blue). (H) Peak provenance: percentage of consensus peaks detected only in AG (green), only in SS (orange) or in both conditions (shared; blue) in the IDR optimal peak sets, genome-wide and for promoter, enhancer and other peaks.

Genomic annotation revealed peaks distributed across intronic regions (39.6%), distal intergenic regions (27.2%) and promoter regions (25.0%), with the remainder mapping to exons, untranslated regions and downstream regions (**Figure 1C**).

ATAC-seq quality was evaluated following ENCODE guidelines using two standard ATAC-seq quality control metrics: fraction of reads in peaks (FRiP) and transcription start site (TSS) enrichment score.

Starvation samples showed markedly reduced FRiP scores (7.2 ± 1.9% vs 56.4 ± 7.7%, p = 0.006) and TSS enrichment scores (3.9 ± 0.8 vs 10.3 ± 1.4, p = 0.005) compared with actively growing samples (**Figure 1D, E**; Supplementary **Table S1A**). These reduced enrichment metrics are consistent with a broad reduction in ATAC-seq signal at canonical accessible regions in SS cells. This pattern is consistent with previous reports of starvation- or stress-associated chromatin compaction [16–18], where compaction was established by orthogonal methods, and represents the accessibility profile that would be expected if a similar process were occurring in starved U2OS cells. However, reduced Tn5 accessibility alone does not constitute evidence of compaction in the present study.

Mapped insert-size analysis showed that both AG and SS libraries retained detectable ATAC-seq fragment architecture, including a shared short-fragment mode, nucleosome-associated fragment classes and Tn5-associated rotational periodicity (**Figure 1A**; Supplementary **Figure S2B-D**; **Table S1B**). Pre-sequencing Fragment Analyzer traces were broader and less resolved in SS than in AG (Supplementary **Figure S2A**), whereas mapped insert-size distributions remained interpretable in both conditions (Supplementary **Figure S2B-D**).

Quantitative analysis of mapped fragment architecture revealed a pattern of size-class shifts that is inconsistent with over-tagmentation (**Table S1B**). NFR-sized fragments (≤100 bp) were significantly less abundant in SS than AG libraries (40.7 ± 2.0% vs 47.0 ± 1.4%, p = 0.015), while mono- and dinucleosomal size-class fractions were preserved and not significantly reduced (mono: 21.3 ± 1.0% vs 19.7 ± 1.1%, p = 0.122; di: 8.9 ± 0.7% vs 7.8 ± 0.7%, p = 0.104). Consequently, the Mono/NFR and Di/NFR ratios were significantly higher in SS than AG (Mono/NFR: 0.526 ± 0.051 vs 0.420 ± 0.036, p = 0.048; Di/NFR: 0.221 ± 0.025 vs 0.166 ± 0.019, p = 0.043), and median insert size was significantly larger in SS (122.7 ± 5.0 bp vs 108.0 ± 4.4 bp, p = 0.020). The ∼10 bp Tn5 rotational periodicity was detectable in both conditions, though with significantly lower magnitude in SS (FFT periodicity score: 1.12 ± 0.19 vs 3.51 ± 0.42, p = 0.004; FFT magnitude: 0.0146 ± 0.0025 vs 0.0650 ± 0.0041, p < 0.001), with no significant difference in dominant period between conditions (SS: 9.5 bp vs AG: 10.9 bp, p = 0.212). Over-tagmentation predicts increased generation of short sub-nucleosomal fragments, which would manifest as a higher NFR fraction, lower Mono/NFR ratio, shorter median insert size, and degraded rotational periodicity. The SS libraries show the opposite pattern on the first three metrics and retained periodicity on the fourth, collectively arguing against over-tagmentation as an explanation for the reduced FRiP and TSS enrichment scores observed in SS cells.

To further test whether the reduced FRiP and TSS enrichment scores in SS reflect genuine loss of peak-centred accessibility rather than a spatial redistribution of signal away from peaks, we quantified ATAC-seq signal, scaled to counts per million mapped fragments, at peak summits (±250 bp), local flanking regions (1-2 kb) and matched genome-wide random-background regions (Supplementary F**igure S10**; **Table S12**). Summit signal was reduced 5.5-fold in SS relative to AG (p = 0.006), consistent in direction and magnitude with the metaplot analysis in Supplementary **Figure S7**, and the proportion of summit-proximal signal concentrated within the summit window itself fell from 89% in AG to 49% in SS. This reduction was accompanied by smaller increases in flank (1.6-fold, p = 0.014) and random-background (2.1-fold, p = 0.004) signal. Because each library contributes a similar total number of fragments, fragments no longer captured in peaks are necessarily distributed over the rest of the genome: with the fraction of fragments in each condition’s own peaks falling from 56% to 7%, a background increase of about 2.1-fold is expected, as observed. Together with the exclusion of the molecular hallmarks of over-tagmentation (short-fragment enrichment, degraded Tn5 periodicity) by the fragment-architecture analysis above, this indicates that the dominant change in SS is a loss of peak-centred accessibility, and that the rise in non-specific signal is its counterpart rather than a separate, non-specific over-digestion process.

Together, these observations support the use of the dataset for differential chromatin-accessibility analysis. We therefore refer to the primary chromatin readout as ATAC-seq/Tn5 accessibility, while recognising that ATAC-seq alone does not directly distinguish physical chromatin compaction from other mechanisms that alter Tn5 integration.

### Starvation induces a distinct chromatin accessibility state

Principal component analysis (PCA) separates the two conditions along the first principal component (PC1: 96.0% of the variance among the 500 most variable peaks) ), with replicates clustering by condition (**Figure 1F**). Pearson correlations were higher within conditions (SS: r = 0.92-0.94; AG: r = 0.96-0.99) than between conditions (r = 0.83-0.90; **Figure 1G**), indicating starvation induces a fundamentally distinct chromatin accessibility state.

Analysis of peak provenance revealed marked asymmetry: of the 88,589 consensus peaks, 63.7% were detected only in AG cells and only 1.9% only in SS cells, while 34.4% were detected in both conditions (**Figure 1H** and **Table 1**). Thus, 94.9% of the peaks detected in starved cells were also detected in growing cells. Starvation-specific peaks were rare at promoters (0.8%) and enhancers (0.7%), suggesting that the starvation response operates predominantly through pre-existing regulatory elements. Promoters were, however, more often retained as detectable peaks in SS cells (57.9% detected in both conditions) than enhancers (31.8%) or other regions (23.2%).

**Table 1.**
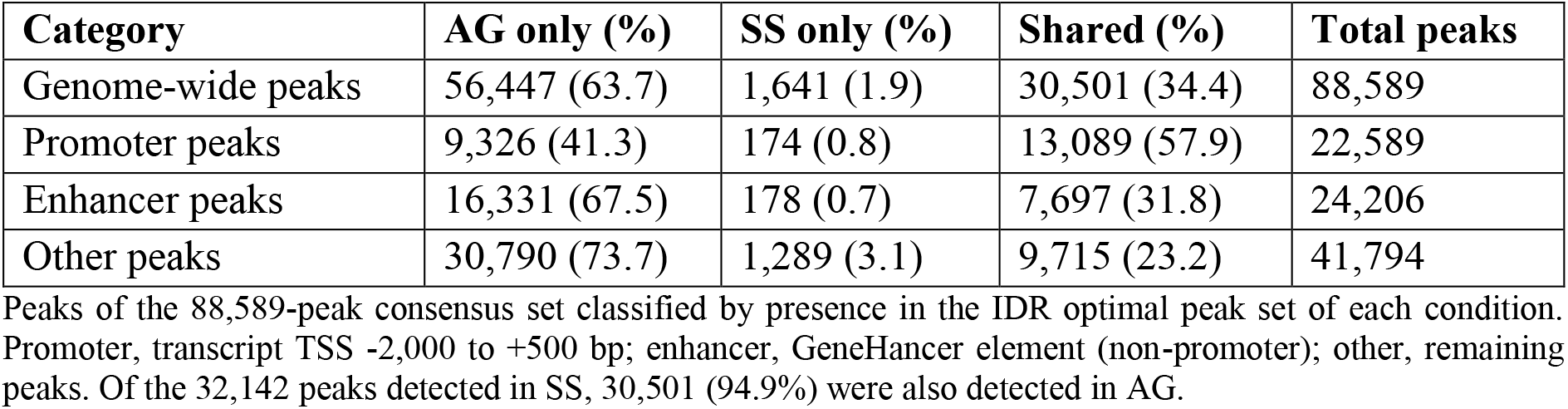
Consensus peak provenance in starved (SS) and growing (AG) U2OS cells.

### Normalization strategy for differential accessibility analysis

The genome-wide reduction in ATAC-seq accessibility makes the choice of normalisation critical [50]. SS and AG libraries contained similar numbers of mapped fragments (22.8 and 22.1 million on average), but the fraction of fragments in peaks was about five-fold lower in SS. Consequently, normalisation methods based on reads in peaks, which equalise peak-associated signal between conditions, produced size factors about five-fold lower for SS than for AG (SS/AG ratio 0.19), whereas normalisation to library size gave nearly equal factors (ratio 1.04; **Table 2**, **Figure 2A**).

**Figure 2.**
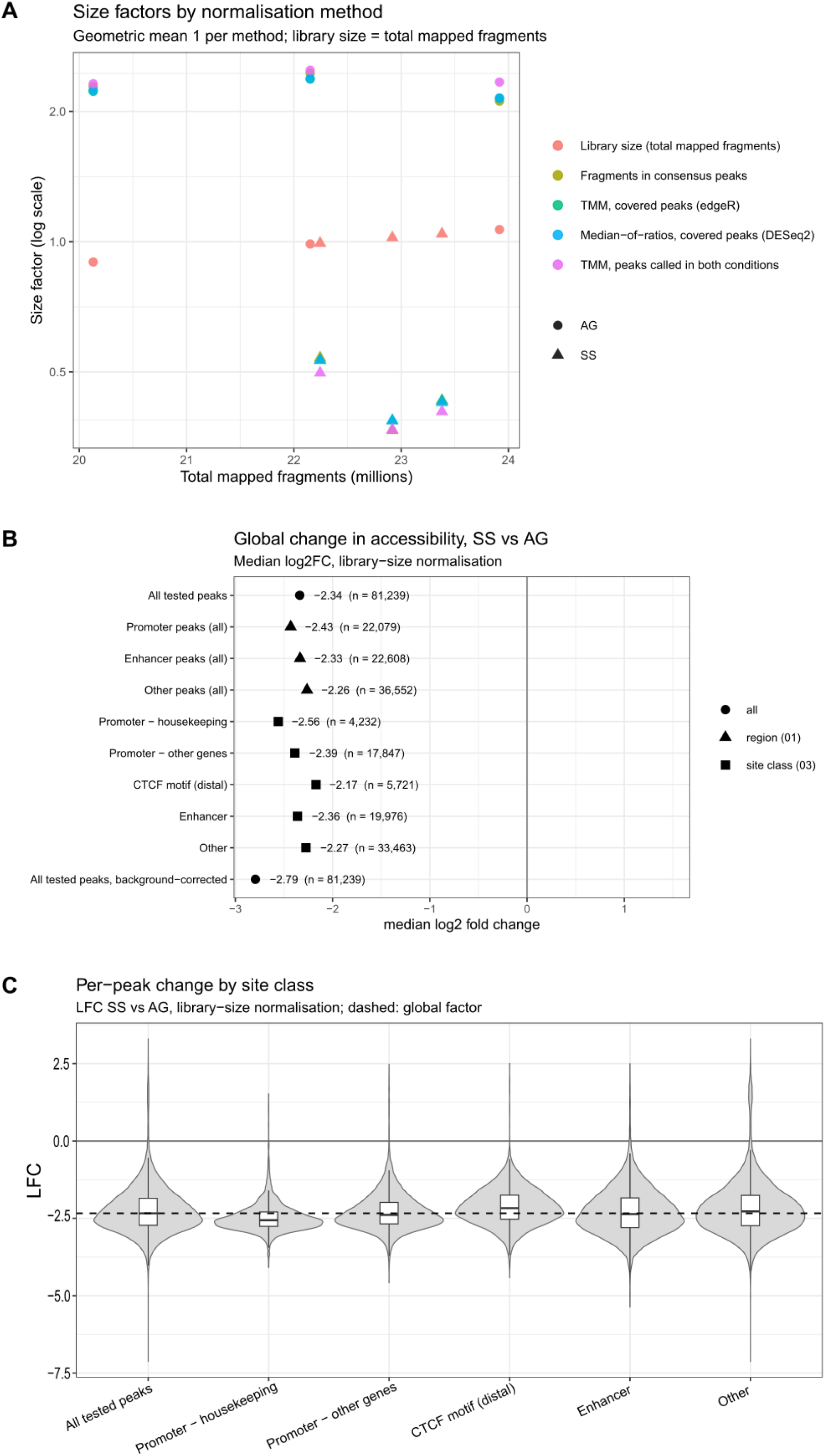
Normalisation of ATAC-seq data under a global loss of accessibility. (A) Size factors for each sample (AG, circles, and SS, triangles) obtained by library-size normalisation (total mapped fragments, primary) and by normalisation to fragments in consensus peaks, plotted against total mapped fragments. Size factors are scaled to a geometric mean of 1 per method (log scale). Library-size factors are similar in AG and SS (SS/AG ratio 1.04), whereas consensus-peak factors are about five-fold lower in SS (ratio 0.19), reflecting the lower fraction of fragments in consensus peaks in SS (10.6% vs 56.5%, **Table 2**). (B) Global change in accessibility: median LFC (SS/AG, DESeq2 with library-size normalisation) of tested peaks by region (promoter, enhancer, other) and site class (housekeeping-gene promoters, promoters of other genes, distal CTCF-motif sites, enhancers, other), and for all tested peaks after correction for the expected non-specific background. Numbers of peaks are shown. (C) Distribution of per-peak LFCs by site class (violin plots with box plots showing median and interquartile range). The dashed line marks the genome-wide median (-2.34).

**Table 2.** Normalisation methods for differential accessibility analysis.

| **Method** | **Role** | **Assumption** | **SF SS/AG** | **r (total fragments)** | **Median LFC, all peaks** | **Median LFC, HK promoters** | **Median LFC, distal CTCF sites** |
|---|---|---|---|---|---|---|---|
| Library size (total mapped fragments) | Primary | Equal input (nuclei, Tn5); retains global changes | 1.038 | 1.00 | -2.29 | -2.54 | -2.11 |
| Fragments in consensus peaks | Relative representation | Total peak signal equal between conditions, removes global changes | 0.193 | -0.38 | 0.08 | -0.15 | 0.25 |

SF, size factor (geometric mean of SS / AG size factors). r, Pearson correlation of per-sample size factors with total mapped fragments. Median log2FC, median log2 ratio of size-factor-normalised condition means (pseudocount 0.5) over peaks with a mean of at least 20 fragments in AG or SS. HK, housekeeping-gene promoters (n = 4,275); distal CTCF sites, non-promoter peaks with a CTCF motif within 100 bp of the summit (n = 6,668). Model-based estimates for library-size normalisation (DESeq2, tested peaks): all -2.34, HK promoters -2.56, distal CTCF -2.17 (Table 3, Figure 2B). Covered peaks, peaks with pooled mean ≥ 20 fragments (n = 68,841).

The two approaches therefore answer different questions. Library-size normalisation expresses the signal at each peak per mapped fragment and retains changes common to all peaks. Normalisation to fragments in consensus peaks equalises the total peak-associated signal between conditions and therefore expresses each peak’s share of the remaining peak signal. On this scale, a positive LFC indicates that a peak retained more of its signal than the peak pool as a whole, which can occur while its signal per mapped fragment decreases. Because libraries were prepared from equal numbers of nuclei using the same batch and amount of Tn5, amplified with the same number of PCR cycles, and showed no evidence of technical over- or under-tagmentation based on fragment-architecture analysis, we used library-size normalisation for the primary analysis and normalisation to fragments in consensus peaks as a secondary measure of relative accessibility preservation.

Under library-size normalisation, the median LFC across all tested peaks was -2.34 (a 5.1-fold reduction, or -2.79 after correction for non-specific background), and the reduction extended to promoters of housekeeping genes (-2.56) and to distal CTCF-motif sites (-2.17), classes of sites often regarded as constitutively accessible (**Figure 2B, C**). Aggregate summit-centred profiles showed the same reduction for all peaks, housekeeping-gene promoters and distal CTCF-motif sites (Supplementary **Figure S7**).

Per-peak LFCs on the two scales differ almost entirely by a constant. They were correlated at r = 0.999 and shifted by 2.42 log2 units (Supplementary **Table S3**), which matches the log2 ratio of the fraction of fragments in consensus peaks in AG and SS (56.5% and 10.6%). Rankings of peaks, correlations between accessibility and RNA changes, and comparisons of accessibility change between groups of genes are therefore essentially unaffected by the choice of scale. Fixed-threshold calls are affected. With library-size normalisation, 698 peaks gained and 77,075 lost signal, whereas with normalisation to fragments in consensus peaks 8,316 peaks gained and 1,457 lost signal relative to the peak pool (Supplementary **Table S3**). Gained and lost peaks, gene-level accessibility calls and the classification of genes by the direction of their changes are therefore reported on the per-fragment scale.

### Starvation induces global changes in chromatin accessibility

Differential accessibility analysis using library-size normalisation identified 77,773 of 81,239 tested peaks (95.7%) as significantly changed in SS relative to AG cells (padj ≤ 0.05, |LFC| ≥ 1), and 99.1% of these showed decreased accessibility (**Figure 3A-C**, **Table 3**). Only 698 peaks gained accessibility, most of them (606) outside annotated promoters and enhancers.

**Figure 3.**
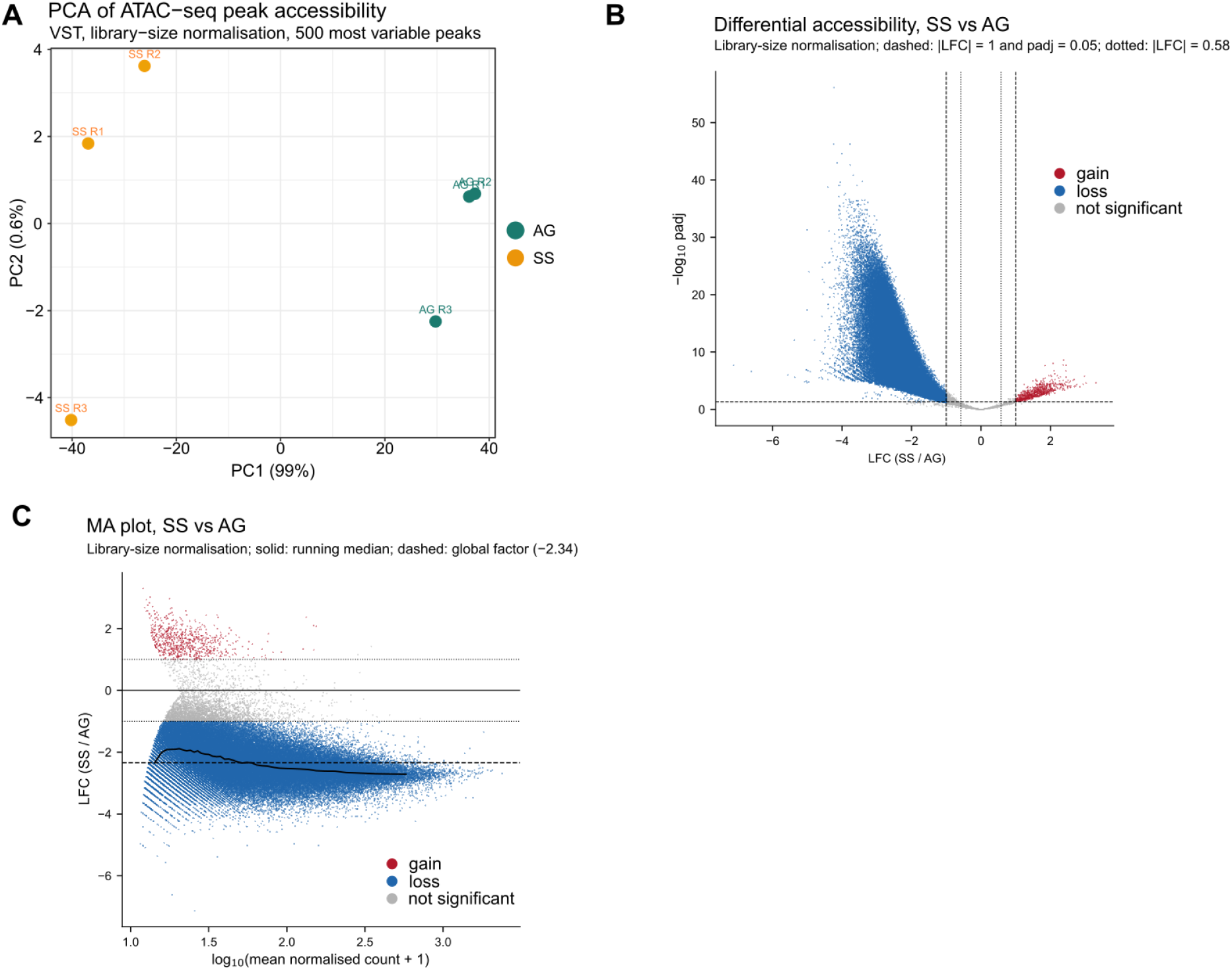
Differential chromatin accessibility analysis reveals widespread loss of accessibility upon starvation. (A) PCA of variance-stabilised, library-size-normalised counts (500 most variable peaks). AG (green) and SS (orange) samples separate along PC1 (99% of variance); PC2 (0.6%) reflects replicate variability. Each point represents an individual sample. (Variance shares differ from Figure 1F, which uses a blind transformation with median-of-ratios size factors for quality control.) (B) Volcano plot of differential accessibility between SS and AG (DESeq2, library-size normalisation; 81,239 tested peaks). The x-axis shows the LFC used for significance calls and the y-axis -log10 padj. Peaks are coloured as gained (red; n = 698), lost (blue; n = 77,075) or not significant (grey). Dashed lines: |LFC| = 1 and padj = 0.05; dotted lines: |LFC| = 0.58 (relaxed threshold, Table S20). (C) MA plot of LFC against log10(mean normalised count + 1) for all tested peaks, coloured as in (B). Solid line, running median; dashed line, genome-wide median LFC (-2.34); dotted lines, |LFC| = 1.

**Table 3.**
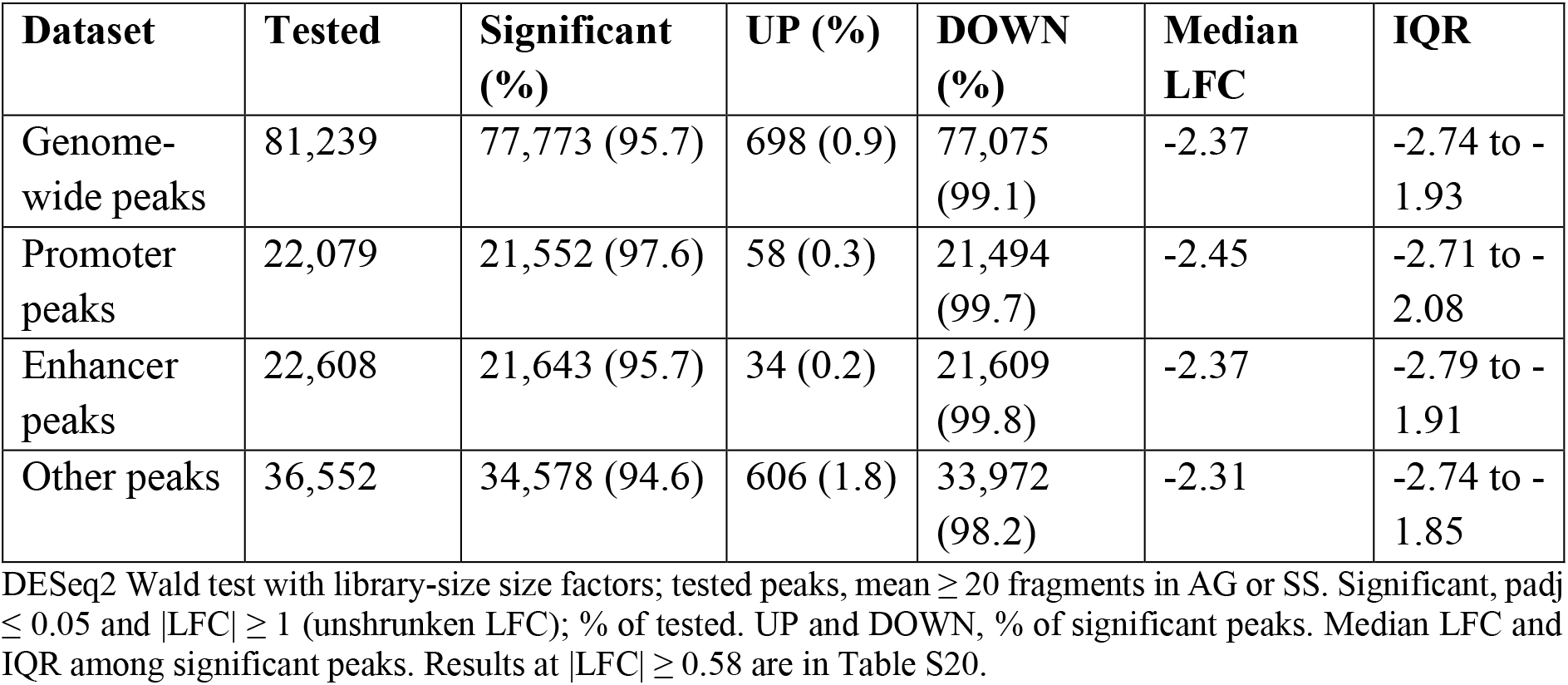
Summary of ATAC-seq differential accessibility analysis.

The median LFC among significant peaks was -2.37 (interquartile range (IQR): -2.74 to -1.93), corresponding to an approximately five-fold reduction in accessibility. Promoter-associated peaks (n = 22,079 tested; 97.6% significant, 99.7% of them decreased; median LFC -2.45) and enhancer-associated peaks (n = 22,608; 95.7% significant, 99.8% decreased; median LFC - 2.37) showed similar patterns, demonstrating a widespread decrease in chromatin accessibility at both proximal and distal regulatory elements. Results at a relaxed threshold (|LFC| ≥ 0.58) were essentially identical (78,815 significant peaks, 99.1% decreased; Table S20).

Collectively, these results demonstrate that starvation elicits a dramatic and highly unidirectional reduction in chromatin accessibility at both proximal promoters and distal enhancers. This pattern is consistent with the large-scale chromatin compaction reported during severe stress in previous studies, where compaction was established using orthogonal methods [16–18]. However, reduced ATAC-seq accessibility alone does not constitute direct evidence of compaction in the present study.

### Cellular poly(A)+ RNA content is unchanged by starvation

Median-of-ratios normalisation assumes that the median gene is unchanged between conditions, an assumption that may be violated by a substantial shift in the total cellular mRNA pool. We therefore measured cellular poly(A)+ RNA content directly. Fluorometric quantification of poly(A)+ RNA purified from defined numbers of cells showed no difference between conditions (SS/AG = 1.01, 95% CI 0.87-1.17; n = 3 cultures per condition). An independent measurement of poly(A)+ RNA as a proportion of input total RNA gave a concordant result (SS/AG = 1.01, 95% CI 0.89-1.14). In both experiments, equivalence testing excluded changes greater than 25% in either direction (p < 0.01). The lower confidence limit for the per-cell measurement corresponds to a maximum uniform displacement of 0.20 in LFC, well below the |LFC| ≥ 1 threshold applied throughout. Thus, cellular mRNA content is preserved during starvation, supporting the RNA-seq analyses reported below without correction for a global shift in mRNA abundance (Supplementary Data, Section 3, and **Table S11**).

### Starvation induces selective and asymmetric changes in mRNA abundance

RNA-seq analysis (two biological replicates per condition; see also Section 2 Supplementary Data) identified 2,664 differentially expressed genes (DEGs; 12.4% of the expressed transcriptome; padj < 0.05, |LFC| ≥ 1; **Figure 4A, B, Table 4**). PCA demonstrated clear separation between growth and starvation conditions (**Figure 4C**). The gene-expression response was strongly asymmetric, with 1,719 genes (64.5%) exhibiting increased RNA abundance and 945 genes (35.5%) exhibiting reduced RNA abundance in starved cells.

**Figure 4.**
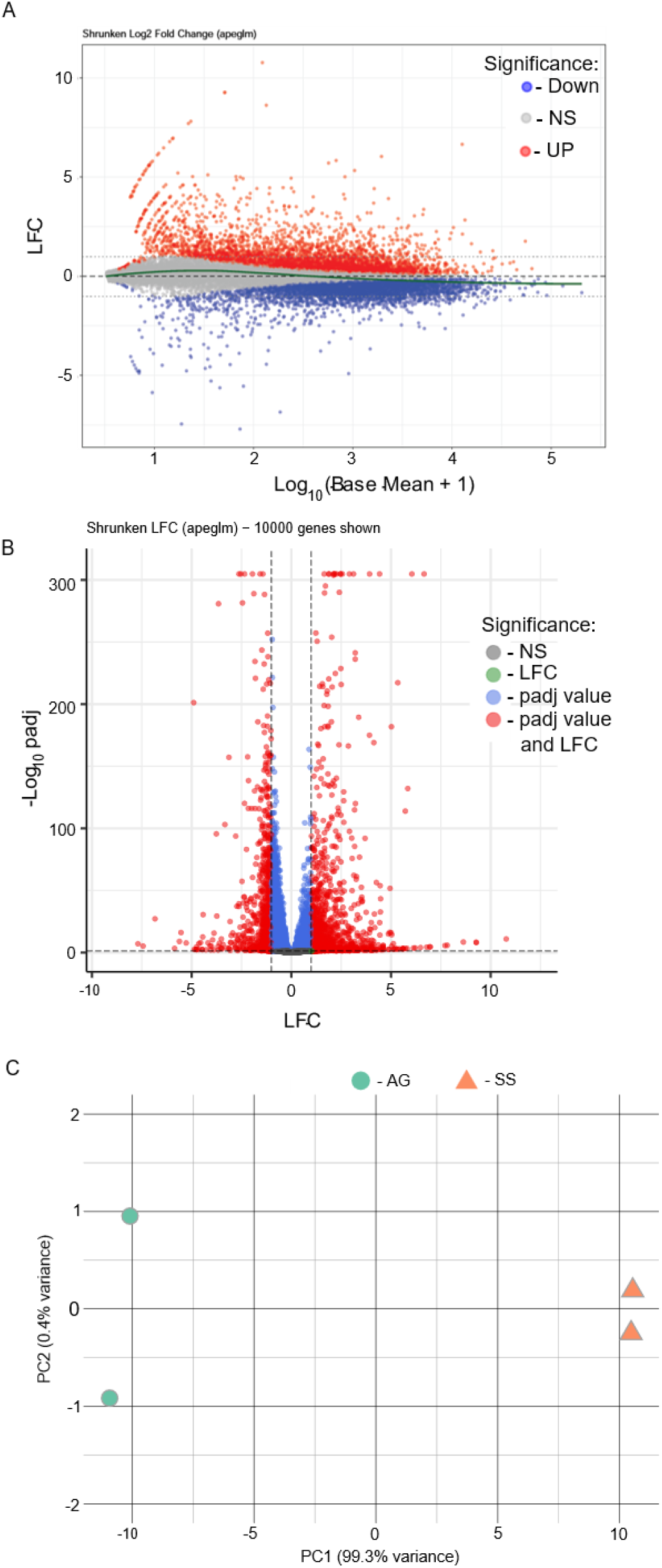
Transcriptome-wide analysis reveals extensive reprogramming upon starvation. (A) MA plot of apeglm-shrunken LFC (SS/AG) against log10(baseMean + 1) for all expressed genes. Genes are coloured by differential expression status (DESeq2 Wald test; padj ≤ 0.05 and |LFC| ≥ 1, called on unshrunken LFC; Table 4): increased in SS (UP, red), decreased in SS (Down, blue) and not significant (NS, grey). Because significance is called on unshrunken LFC, some significant genes lie inside the ±1 lines after shrinkage. Dashed line, LFC = 0; dotted lines, LFC = ±1; green line, loess trend. (B) Volcano plot of apeglm-shrunken LFC against −log10 padj (10,000 genes shown). Genes are coloured by the thresholds they pass: neither (NS, grey), |LFC| ≥ 1 only (LFC, green), padj ≤ 0.05 only (padj value, blue) or both (padj value and LFC, red). Dashed lines mark |LFC| = 1 and padj = 0.05. Genes with padj below the numerical limit (< 10^−300^) are plotted at the top of the axis. (C) PCA of variance-stabilised (VST) counts of the 2,664 DEGs. AG (green circles) and SS (orange triangles) samples separate along PC1 (99.3% of variance); PC2 explains 0.4%. Each point is one biological replicate (n = 2 per condition).

**Table 4.** Summary of RNA-seq differential expression analysis.

| **Dataset** | **Expressed** | **Significant\*** | **UP (%)** | **DOWN (%)** |
|---|---|---|---|---|
| SS vs AG | 21505 | 2664 | 64.5 | 35.5 |

\* **-** significance criteria: padj ≤ 0.05, \|LFC\| ≥ 1; no baseMean filtering

### Pathway analysis reveals coordinated gene-expression reprogramming

Preranked fast gene set enrichment analysis (fGSEA) across Gene Ontology (GO), Hallmark and transcription factor target (TFT) collections identified 1,078 significantly enriched gene sets (FDR < 0.01), comprising 325 positively enriched and 753 negatively enriched pathways (**Table S4**). To reduce redundancy, pathways were clustered into 41 biological categories based on leading-edge gene overlap, and category-level enrichment was quantified using weighted normalised enrichment scores (wNES) and Stouffer’s combined Z-scores (see Methods; **Figure 5, Table S5**).

**Figure 5.**
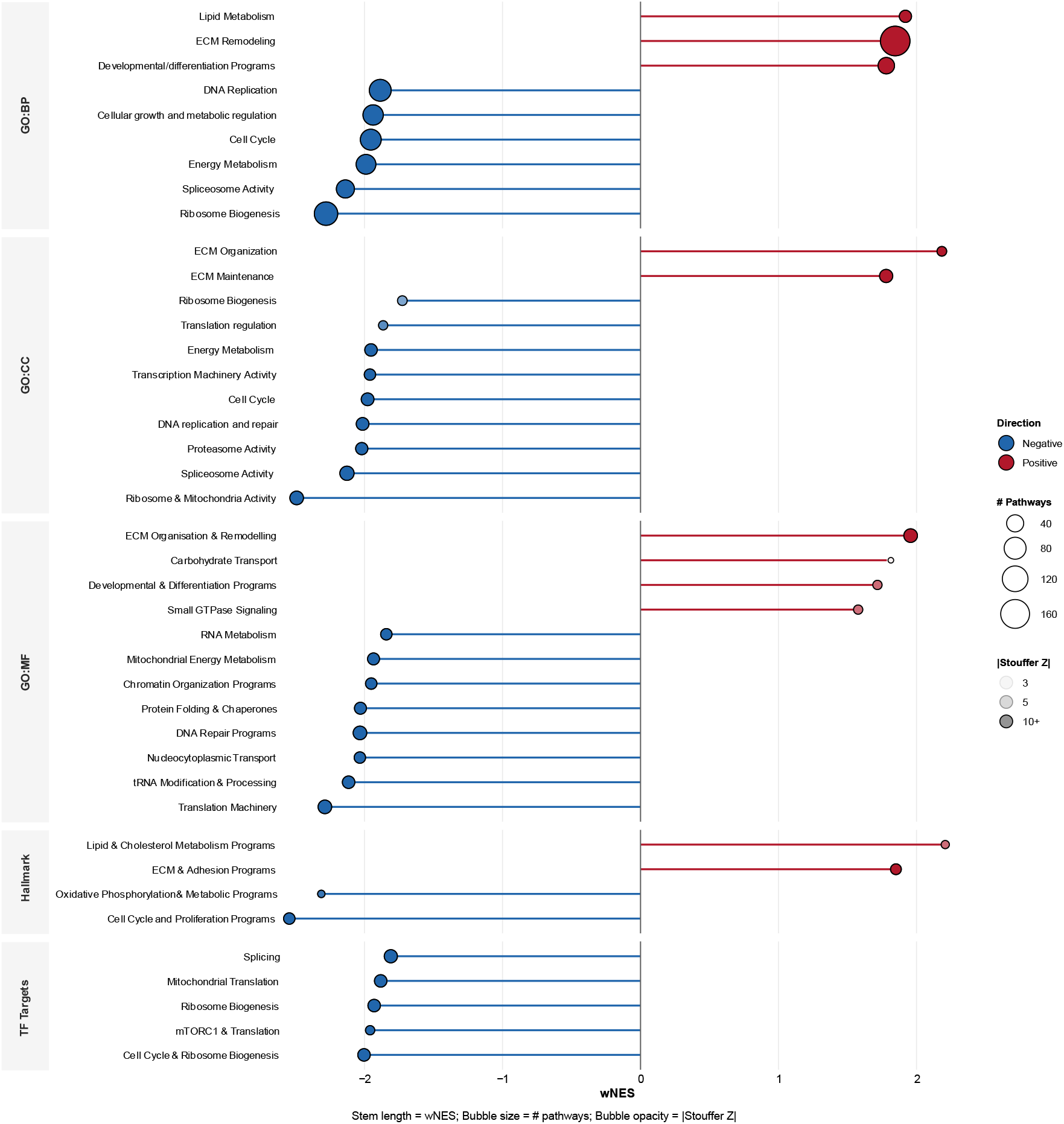
Differential regulation of high-level biological categories under starvation. Lollipop plot showing curated biological categories derived from fGSEA analysis (*p*adj ≤ 0.01). Categories are organised by gene set collection (GO:BP, GO:CC, GO:MF, Hallmark, TF Targets). Stem length and position represent weighted normalised enrichment score (wNES), with positive values (red) indicating upregulation and negative values (blue) indicating downregulation in starved cells. Bubble size corresponds to the number of individual pathways within each category. Bubble opacity reflects the magnitude of Stouffer’s combined Z-score (key: |Z| = 3, 5 and 10 or more).

Starvation was associated with strong and consistent negative enrichment of biosynthetic and proliferative programmes (30 of 41 categories), including ribosome biogenesis, cell cycle, DNA replication, oxidative phosphorylation and spliceosome activity (**Figure S5**). All five significant TFT categories also showed negative enrichment, indicating reduced steady-state abundance of mRNAs associated with proliferation-linked transcriptional regulators.

In contrast to this widespread negative enrichment, 11 biological categories showed positive enrichment in SS, with extracellular matrix (ECM)-related programmes displaying the strongest positive enrichment across multiple ontologies. Lipid and cholesterol metabolism, developmental signalling and small GTPase pathways were also positively enriched. The enrichment of ECM organisation and remodelling genes under starvation is consistent with recent reports [57, 58].

### Regulator enrichment nominates Polycomb-associated, MYC/MAX-linked and module-specific regulatory programmes

To identify candidate regulatory signatures associated with the starvation RNA-abundance response, we analysed transcription factor, perturbation and chromatin-state enrichment among starvation-regulated genes. Genes with increased RNA abundance in SS cells were strongly enriched for Polycomb-associated targets, including EZH2 (padj = 1.2 × 10^−26^), SUZ12 (padj = 8.7 × 10^−16^), RNF2/RING1B (padj = 5.2 × 10^−10^), and JARID2 (**Table S13**). Consistent with this pattern, H3K27me3-associated gene sets were also strongly enriched among upregulated genes (padj = 3.2 × 10^−33^ across 25 tissue/cell-line terms). Additional enrichment for SMAD4 (padj = 9.1 × 10^−11^) and SMAD3 (padj = 4.4 × 10^−9^) targets linked the RNA-up response to TGF-β/BMP-associated regulatory programmes. Together, these results suggest that starvation-upregulated genes are enriched for Polycomb-associated and developmental/stress-responsive regulatory contexts.

Genes with reduced RNA abundance showed a contrasting enrichment pattern. MYC (padj = 1.5 × 10^−21^) and MAX (padj = 2.5 × 10^−19^) were among the strongest regulator-associated enrichments, accompanied by USF1/USF2 and E2F6 (**Table S13**). Perturbation signatures reinforced this assignment, with MYC loss-of-function representing the strongest genome-wide signature (padj = 6.3 × 10⁻⁵³), together with HSF1-associated signatures (padj = 1.1 × 10^−41^). In contrast to the upregulated set, downregulated genes were preferentially associated with active chromatin marks, including H3K27ac, H3K36me3 and H3K79me2 (**Table S15**). These results support the interpretation that starvation suppresses an actively transcribed, MYC/MAX-linked biosynthetic programme while selectively increasing RNA abundance from a distinct set of regulatory contexts.

Programme-level analysis refined these assignments (**Table S14**). PRC2/PRC1 target enrichment and H3K27me3-associated enrichment were strongest in the ECM Remodelling and Developmental/Differentiation programmes, and remained detectable in the smaller Small GTPase Signalling and Lipid/Cholesterol programmes. The Lipid/Cholesterol programme showed a distinct enrichment for SREBF1 (padj = 2.3 × 10^−25^, OR = 18.0) and SREBF2 (padj = 5.3 × 10^−12^), supported by SREBF2 perturbation signatures (padj = 1.4 × 10^−22^, OR = 40). Thus, the adaptive RNA-up response appears to combine a broad Polycomb-associated component with module-specific regulatory signatures, most clearly SREBF1/2 in lipid and cholesterol metabolism.

### Exon-intron analyses support altered RNA production during starvation

To assess whether starvation-induced changes in steady-state RNA abundance reflect altered RNA production, transcript stability, or both, we examined genome-wide exon-intron concordance and formal Exon-Intron Split Analysis (EISA) [35], with published transcript half-lives as context. Because steady-state RNA abundance integrates synthesis and turnover, these indirect analyses cannot exclude post-transcriptional regulation or quantify transcription directly. Rather, they test whether the observed changes are compatible with widespread mRNA stabilisation as the predominant explanation.

In published half-life measurements from nutrient-replete HeLa cells [59], most genes with increased RNA abundance were short-lived transcripts (88.7% with half-life ≤ 9 h, half of the starvation period; Supplementary **Figure S4**). Without continued production or marked stabilisation, such transcripts would be depleted during 18 h of starvation, which argues against passive persistence of pre-existing RNA, although these external data cannot distinguish increased production from selective stabilisation.

We next analysed exon-intron concordance using the ribosomal RNA-depleted total RNA-seq dataset. Exonic reads were quantified directly, whereas intronic counts were derived by subtracting exon counts from gene-body counts; the two compartments were then analysed separately for differential abundance. Across 12,734 filtered protein-coding genes, exonic and intronic LFCs were positively correlated (Pearson r = 0.690; Spearman ρ = 0.625; both p < 1 × 10^−300^; Supplementary **Figure S8A**). Across all filtered genes, irrespective of statistical significance, 71.9% showed exonic and intronic changes in the same direction. Independent classification of each compartment as UP, DOWN, or UNCHANGED (padj ≤ 0.05, |log2FC| ≥ 0.5) gave Cohen’s κ = 0.328 across the complete 3 × 3 table (69.0% observed agreement versus 53.8% expected by chance; Supplementary Table S9). This value was strongly influenced by the 10,989 genes classified as UNCHANGED in one or both compartments. Among genes with significant directional calls in both compartments (n = 1,745; 13.7% of filtered genes), 98.0% changed concordantly (1,710/1,745; label-shuffled permutation p < 1 × 10^−4^; Supplementary **Figure S8B, C**). The genome-wide median difference between exonic and intronic responses (Δ = exon LFC - intron LFC) was +0.003, close to zero (Supplementary Figure S8C). Increased exonic abundance was generally accompanied by increased, rather than reduced, intronic signal. Because intronic changes provide an indirect proxy for changes in pre-mRNA production, this pattern is consistent with altered RNA production contributing substantially to the response, but does not directly measure nascent transcription.

To assess post-transcriptional regulation at individual genes, we applied the EISA interaction model [35, 60]. The additional internal filtering used by EISA retained 10,881 genes, a subset of the 12,734 genes included in the concordance analysis. Summary statistics from the two analyses are therefore not directly comparable. Within the EISA gene set, exonic and intronic fold changes were strongly correlated (Pearson r = 0.720), and the median interaction term (Dex - Din) was +0.018 (IQR -0.273 to +0.350), again centred close to zero (Supplementary **Figure S8D, E**).

At FDR < 0.05 and |Dex - Din| ≥ 0.5, 2,746 genes (25.2%) showed significant exon-intron divergence, comprising 1,609 positive-interaction stabilisation candidates and 1,137 negative-interaction destabilisation candidates. A further 1,620 genes had significant interactions below the effect-size threshold, while 6,515 were assigned to the concordant/background class (median Dex - Din = +0.045). Applying the more stringent threshold of FDR < 0.05 and |Dex - Din| ≥ 1.0 identified 542 high-confidence candidates (5.0%): 437 stabilisation candidates (median Dex - Din = +1.21) and 105 destabilisation candidates (median Dex - Din = -1.17). A positive EISA interaction does not by itself demonstrate active transcript stabilisation. During reduced RNA production, intronic signal can decline faster than the pre-existing mature RNA pool is cleared, causing Dex to exceed Din. Consistent with this kinetic interpretation, only 33 of the 437 stabilisation candidates showed a significant increase in intronic signal; the remainder showed reduced or unchanged intronic signal. Among genes with significantly increased exonic abundance (n = 2,005), approximately 11% were high-confidence stabilisation candidates. Thus, most genes with increased exonic abundance were not assigned to this class at the stringent threshold.

Reduced abundance of spliceosome-related genes during starvation (Supplementary **Figure S5**) raises the possibility that impaired splicing or intron retention could affect the exon-intron signal. We therefore performed two sensitivity analyses. First, among genes retained by EISA (n = 10,881), we tested whether negative-interaction destabilisation candidates were enriched among genes with many introns. Intron retention increases intronic counts and therefore shifts Dex - Din in the negative direction, so such enrichment would be expected if retention were a major source of the signal. Instead, the proportion of destabilisation candidates decreased across increasing intron-count quartiles (1.5%, 0.8%, 0.9%, and 0.5%; χ^2^ = 17.5, p = 0.0004). This inverse association remained significant after adjustment for baseline expression (logistic regression, p < 0.001; Supplementary **Figure S9A**). Second, we recalculated exon-intron concordance after excluding genes with high splicing complexity. Excluding either the top intron-count quartile (n = 3,012) or all genes with ≥20 introns (n = 2,173) had little effect: Pearson r changed from 0.690 to 0.671-0.675, the same-sign fraction from 71.9% to 71.0-71.2%, and concordance among directionally classified genes from 98.0% to 98.2% (Supplementary **Figure S9B**; **Table S10**).

These sensitivity analyses argue that neither the broader exon-intron concordance pattern nor the negative-interaction class is driven primarily by genes with high splicing complexity. However, intron count is only an indirect proxy for intron retention and cannot exclude transcript-specific retention. Intron retention would shift Dex - Din in the direction opposite to the larger positive-interaction class, but EISA alone cannot distinguish active stabilisation from kinetic persistence of mature transcripts following reduced RNA production.

Taken together, the exon-intron concordance and EISA analyses support altered RNA production as a major contributor to starvation-associated changes in RNA abundance. EISA additionally identified candidate post-transcriptional regulation in 5.0% of tested genes at the stringent threshold. The predominance of positive-interaction candidates, most of which had reduced or unchanged intronic signal, is compatible with persistence of mature transcripts during reduced RNA production. These results are consistent with transcript-specific post-transcriptional regulation superimposed on broader changes in RNA production during stress [60], rather than a uniform, transcriptome-wide shift in mRNA stability. Genome-wide assays such as PRO-seq, GRO-seq and SLAM-seq would measure nascent RNA directly, rather than infer changes in RNA production from intronic reads as in EISA. We examine this relationship experimentally at selected *loci* using paired RT-qPCR measurements of nascent and total RNA, as described below.

### Weak global coupling between chromatin accessibility and gene expression

To quantify the relationship between chromatin accessibility and mRNA-abundance changes, we integrated RNA-seq and ATAC-seq. Of 16,484 expressed genes, 11,792 had at least one promoter-associated peak and 6,632 had at least one linked enhancer peak, and 5,805 had both (**Table 5A, C**).

**Table 5.**
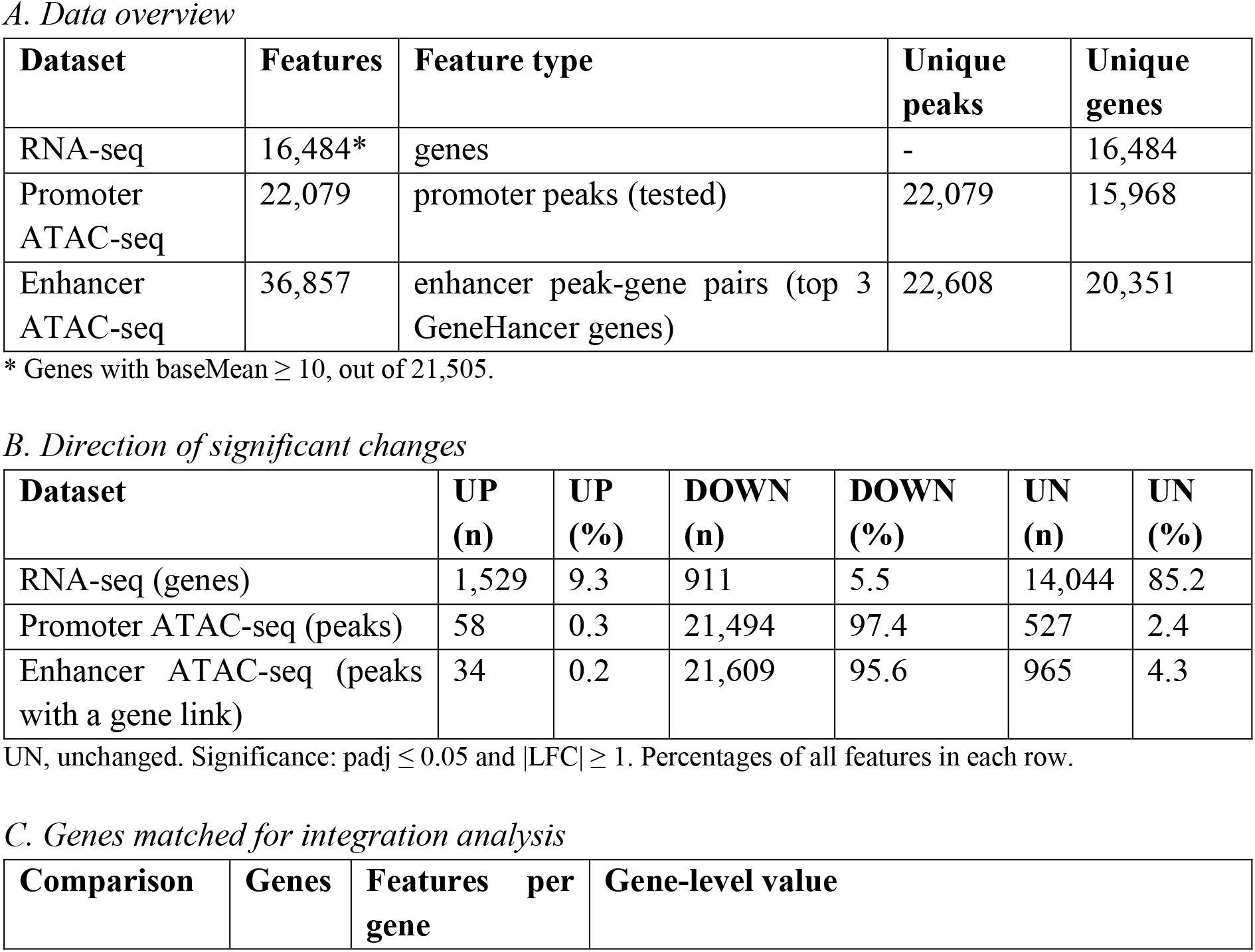
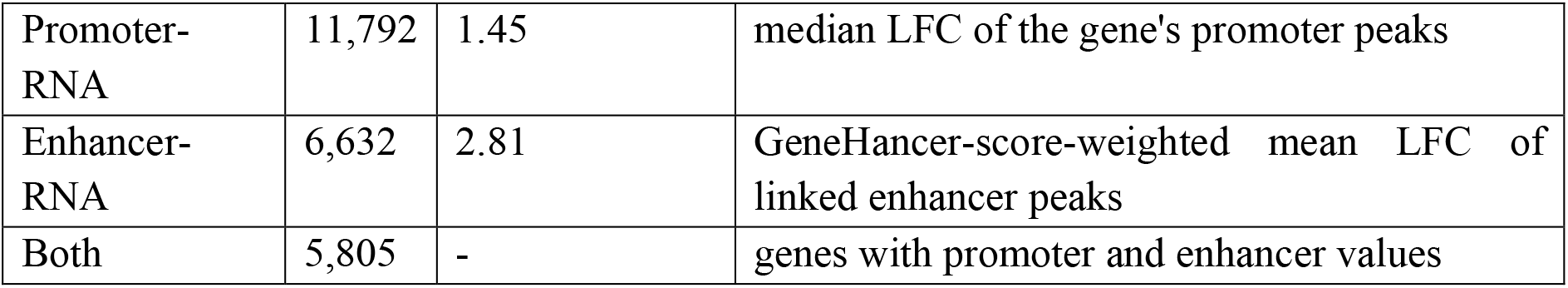
Summary of ATAC-seq and RNA-seq statistics.

| *A. Data overview* |  |  |  |  |  |  |
| **Dataset** | **Features** | **Feature type** | **Unique peaks** | **Unique genes** |  |  |
|---|---|---|---|---|---|---|
| RNA-seq | 16,484* | genes | - | 16,484 |  |  |
| Promoter ATAC-seq | 22,079 | promoter peaks (tested) | 22,079 | 15,968 |  |  |
| Enhancer ATAC-seq | 36,857 | enhancer peak-gene pairs (top 3 GeneHancer genes) | 22,608 | 20,351 |  |  |
| *B. Direction of significant changes* |  |  |  |  |  |  |
| **Dataset** | **UP (n)** | **UP (%)** | **DOWN (n)** | **DOWN (%)** | **UN (n)** | **UN (%)** |
| RNA-seq (genes) | 1,529 | 9.3 | 911 | 5.5 | 14,044 | 85.2 |
| Promoter ATAC-seq (peaks) | 58 | 0.3 | 21,494 | 97.4 | 527 | 2.4 |
| Enhancer ATAC-seq (peaks with a gene link) | 34 | 0.2 | 21,609 | 95.6 | 965 | 4.3 |
| *C. Genes matched for integration analysis* |  |  |  |  |  |  |
| **Comparison** | **Genes** | **Features per gene** | **Gene-level value** |  |  |  |
| Promoter-RNA | 11,792 | 1.45 | median LFC of the gene's promoter peaks |  |  |  |
| Enhancer-RNA | 6,632 | 2.81 | GeneHancer-score-weighted mean LFC of linked enhancer peaks |  |  |  |
| Both | 5,805 | - | genes with promoter and enhancer values |  |  |  |

\* Genes with baseMean ≥ 10, out of 21,505.
UN, unchanged. Significance: padj ≤ 0.05 and \|LFC\| ≥ 1. Percentages of all features in each row.

While 97% of promoter peaks and 96% of enhancer peaks showed significantly decreased accessibility, only 15% of expressed genes were differentially regulated (**Figure 6A**, **Table 5B**). This disparity suggests that the widespread reduction in chromatin accessibility does not translate directly into a proportional reduction in steady-state mRNA levels.

**Figure 6.**
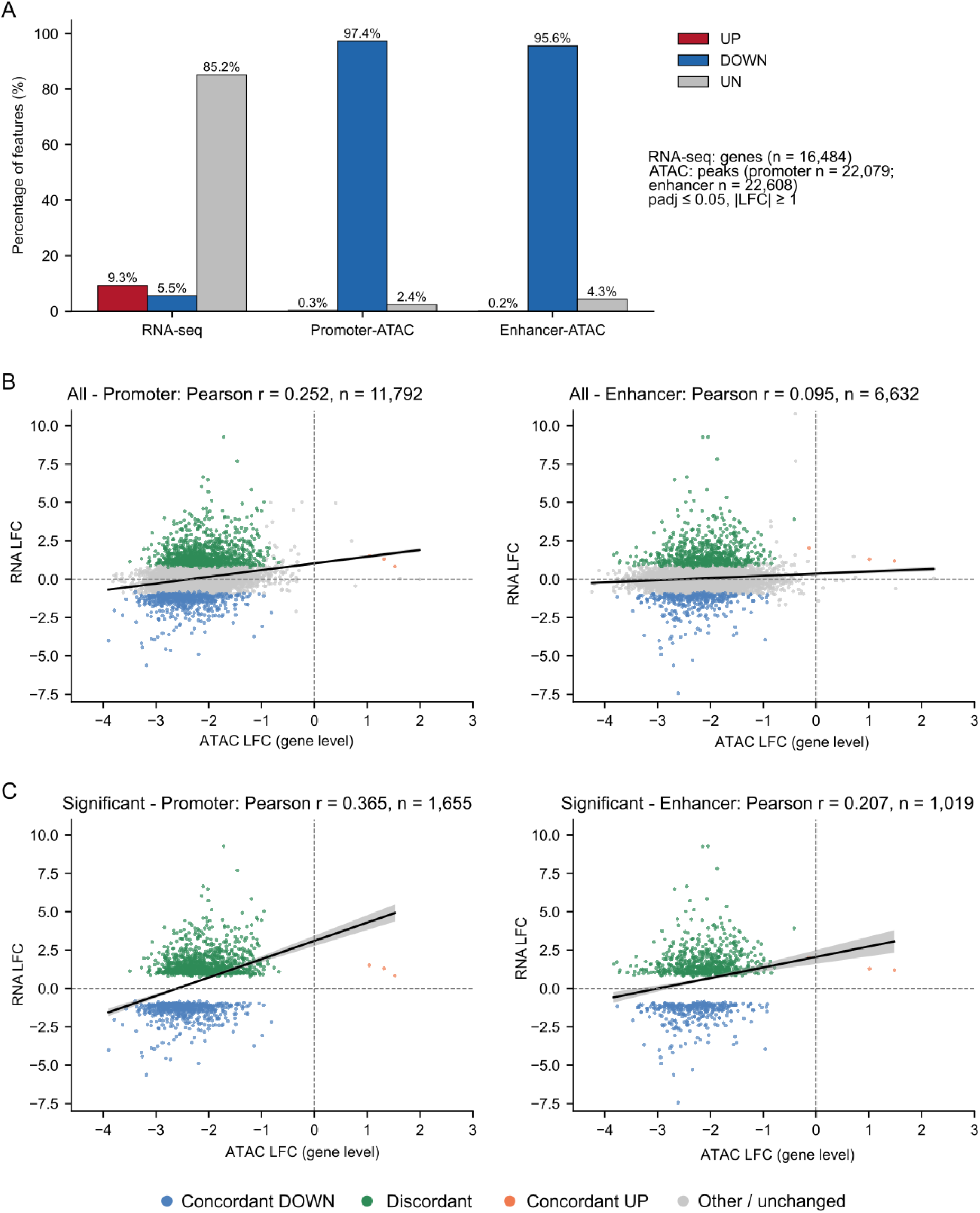
Integrated analysis reveals modest coupling between ATAC-seq and RNA-seq changes during starvation. (A) Direction of significant changes (padj ≤ 0.05, |LFC| ≥ 1) for RNA-seq (genes, n = 16,484), promoter ATAC-seq peaks (n = 22,079) and enhancer ATAC-seq peaks linked to a gene (n = 22,608): UP (red), DOWN (blue) and unchanged (UN, grey). RNA abundance changes are selective, whereas promoter and enhancer accessibility changes are overwhelmingly decreases. (B) Gene-level accessibility changes versus mRNA changes for all matched genes: promoter (left; median LFC of the gene’s promoter peaks; n = 11,792; Pearson r = 0.252) and enhancer (right; GeneHancer-weighted mean LFC of linked enhancer peaks; n = 6,632; r = 0.095). apeglm-shrunken LFC. Black lines show linear regression fits with 95% confidence bands. Points are coloured by gene-level concordance: concordant decrease (blue), discordant (green), concordant increase (orange), other or unchanged (grey). (C) As (B), restricted to genes called significant in both RNA-seq and ATAC-seq (padj ≤ 0.05, |LFC| ≥ 1): promoter (left; n = 1,655; r = 0.365) and enhancer (right; n = 1,019; r = 0.207). This selection on significance inflates the correlations (secondary analysis; Table 6).

**Table 6.**
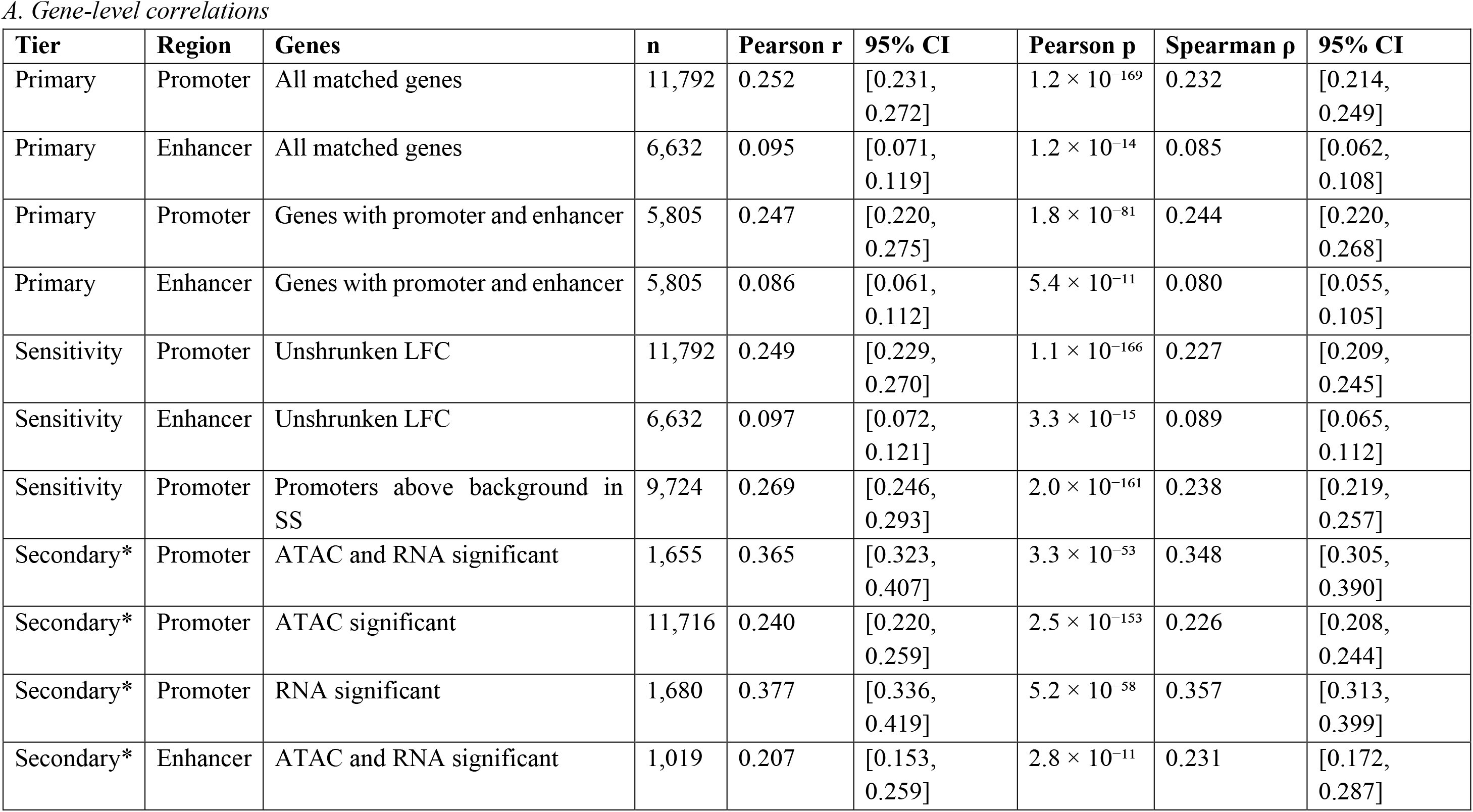
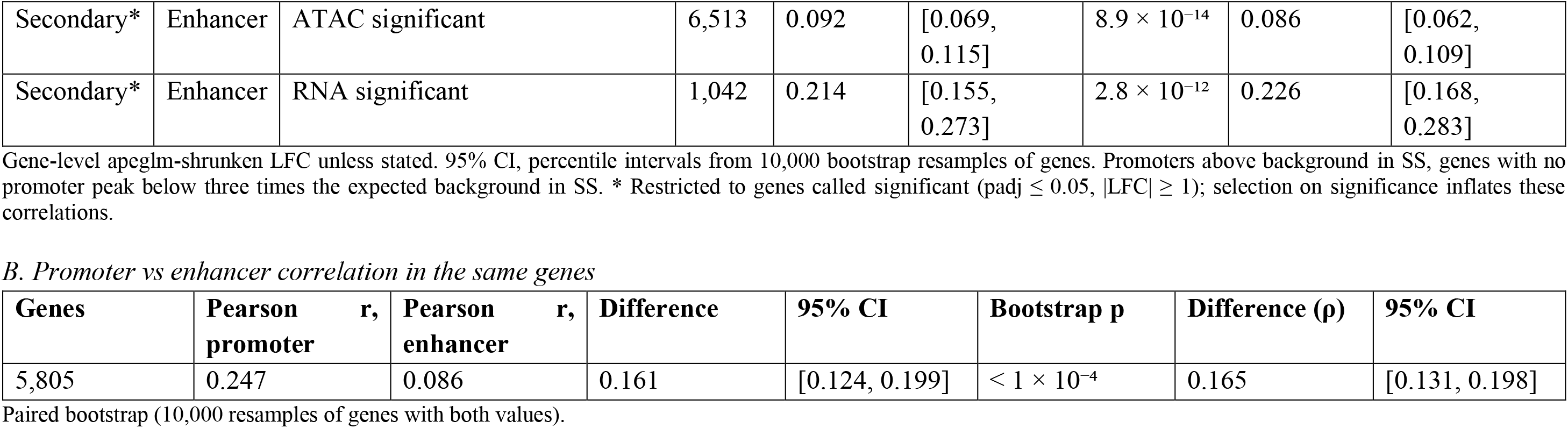
Correlation between chromatin accessibility and gene expression changes.

| **Tier** | **Region** | **Genes** | **n** | **Pearson r** | **95% CI** | **Pearson p** | **Spearman ρ** | **95% CI** |
|---|---|---|---|---|---|---|---|---|
| Primary | Promoter | All matched genes | 11,792 | 0.252 | [0.231, 0.272] | 1.2 × 10<sup>−169</sup> | 0.232 | [0.214, 0.249] |
| Primary | Enhancer | All matched genes | 6,632 | 0.095 | [0.071, 0.119] | 1.2 × 10<sup>−14</sup> | 0.085 | [0.062, 0.108] |
| Primary | Promoter | Genes with promoter and enhancer | 5,805 | 0.247 | [0.220, 0.275] | 1.8 × 10<sup>−81</sup> | 0.244 | [0.220, 0.268] |
| Primary | Enhancer | Genes with promoter and enhancer | 5,805 | 0.086 | [0.061, 0.112] | 5.4 × 10<sup>−11</sup> | 0.080 | [0.055, 0.105] |
| Sensitivity | Promoter | Unshrunken LFC | 11,792 | 0.249 | [0.229, 0.270] | 1.1 × 10<sup>−166</sup> | 0.227 | [0.209, 0.245] |
| Sensitivity | Enhancer | Unshrunken LFC | 6,632 | 0.097 | [0.072, 0.121] | 3.3 × 10<sup>−15</sup> | 0.089 | [0.065, 0.112] |
| Sensitivity | Promoter | Promoters above background in SS | 9,724 | 0.269 | [0.246, 0.293] | 2.0 × 10<sup>−161</sup> | 0.238 | [0.219, 0.257] |
| Secondary* | Promoter | ATAC and RNA significant | 1,655 | 0.365 | [0.323, 0.407] | 3.3 × 10<sup>−53</sup> | 0.348 | [0.305, 0.390] |
| Secondary* | Promoter | ATAC significant | 11,716 | 0.240 | [0.220, 0.259] | 2.5 × 10<sup>−153</sup> | 0.226 | [0.208, 0.244] |
| Secondary* | Promoter | RNA significant | 1,680 | 0.377 | [0.336, 0.419] | 5.2 × 10<sup>−58</sup> | 0.357 | [0.313, 0.399] |
| Secondary* | Enhancer | ATAC and RNA significant | 1,019 | 0.207 | [0.153, 0.259] | 2.8 × 10<sup>−11</sup> | 0.231 | [0.172, 0.287] |
| Secondary* | Enhancer | ATAC significant | 6,513 | 0.092 | [0.069, 0.115] | 8.9 × 10<sup>−14</sup> | 0.086 | [0.062, 0.109] |
| Secondary* | Enhancer | RNA significant | 1,042 | 0.214 | [0.155, 0.273] | 2.8 × 10<sup>−12</sup> | 0.226 | [0.168, 0.283] |
| *B. Promoter vs enhancer correlation in the same genes* |  |  |  |  |  |  |  |  |
| **Genes** | **Pearson r, promoter** | **Pearson r, enhancer** | **Difference** | **95% CI** | **Bootstrap p** | **Difference (ρ)** | **95% CI** |  |
| 5,805 | 0.247 | 0.086 | 0.161 | [0.124, 0.199] | < 1 × 10<sup>−4</sup> | 0.165 | [0.131, 0.198] |  |

Gene-level apeglm-shrunken LFC unless stated. 95% CI, percentile intervals from 10,000 bootstrap resamples of genes. Promoters above background in SS, genes with no promoter peak below three times the expected background in SS. \* Restricted to genes called significant (padj ≤ 0.05, \|LFC\| ≥ 1); selection on significance inflates these correlations.
Paired bootstrap (10,000 resamples of genes with both values).

Across all matched genes, promoter accessibility changes correlated modestly with mRNA-abundance changes (Pearson r = 0.25, 95% CI 0.23-0.27; Spearman ρ = 0.23; n = 11,792), whereas enhancer accessibility changes showed a weaker correlation (r = 0.10, 95% CI 0.07-0.12; n = 6,632; **Figure 6B**, **Table 6**). In the 5,805 genes with both values, the promoter correlation exceeded the enhancer correlation by 0.16 (95% CI 0.12-0.20). The promoter correlation was unchanged when genes with weak, near-background promoter peaks were excluded (r = 0.27; n = 9,724) or when unshrunken fold changes were used (r = 0.25), indicating that the modest coupling is not explained by noise at weak peaks. Restricting the analysis to genes with significant changes in both assays increased the correlations (promoter r = 0.37; enhancer r = 0.21; **Figure 6C**), as expected when genes are selected for large changes, but the associations remained modest.

Within-condition promoter accessibility-expression correlations were positive but modest in both states (AG: Pearson r = 0.28, Spearman ρ = 0.27; SS: Pearson r = 0.25, Spearman ρ = 0.25; Supplementary **Table S8**). These values are similar in magnitude to the correlation between promoter accessibility and mRNA changes, indicating that promoter accessibility and mRNA abundance are only modestly coupled within each condition and remain so during the starvation response.

Directional agreement between accessibility and mRNA changes was limited: 5.8% of genes showed concordant changes at promoters and 5.6% at enhancers (**Figure 7A**, **Table 7**), and these were almost exclusively coordinated decreases; concordant increases were rare (3 genes). Because 99.5% of matched genes showed reduced promoter accessibility, these categorical comparisons are dominated by the global loss, and we therefore examined the relationship on a continuous scale in the next section. Together, these analyses show that starvation-associated changes in chromatin accessibility and mRNA abundance are only modestly coupled at the level of individual genes, consistent with previous reports of limited accessibility-expression coupling under cellular perturbation [9, 10].

**Figure 7.**
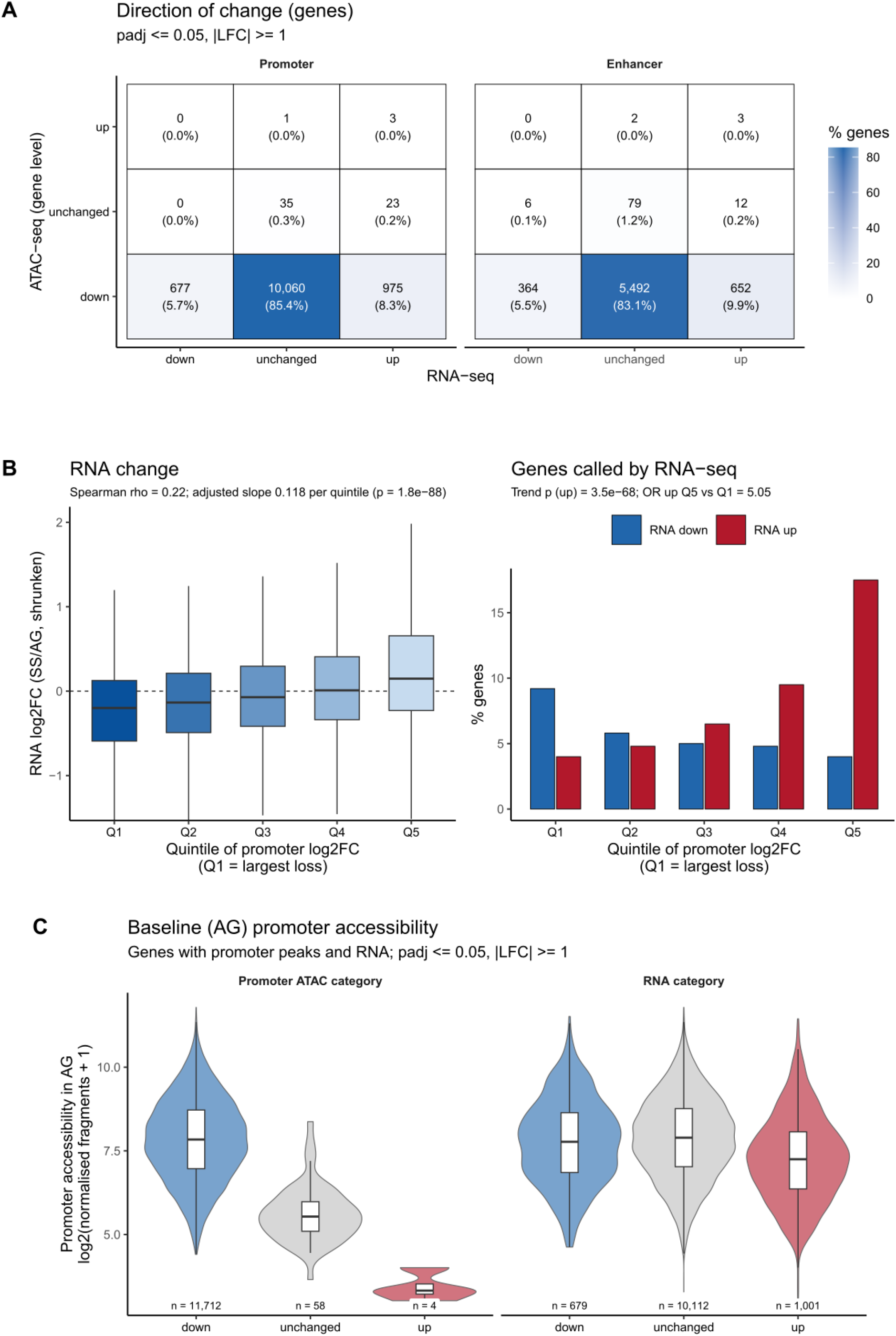
Accessibility-stratified analysis of gene-expression changes. (A) Gene-level contingency heatmaps of the direction of change in accessibility (rows) and mRNA abundance (columns) for promoters (left; n = 11,774 genes) and enhancers (right; n = 6,610 genes); padj ≤ 0.05, |LFC| ≥ 1. Cells show numbers and percentages of genes; colour shows the percentage. Odds ratios are given in Table 7. (B) mRNA changes across quintiles of promoter accessibility change (Q1, largest loss; Q5, smallest loss; 11,792 genes). Left, RNA LFC (shrunken) per quintile (box plots; Spearman ρ = 0.22; slope adjusted for baseline accessibility 0.118 per quintile, p = 1.8 × 10^−88^). Right, percentage of genes with increased (red) or decreased (blue) mRNA per quintile (Cochran-Armitage trend test for increased mRNA, p = 3.5 × 10^−68^; odds ratio Q5 vs Q1 = 5.05). (C) Baseline promoter accessibility in AG cells, log2(median normalised fragments + 1) of the gene’s promoter peaks, by promoter accessibility category (left; reduced n = 11,712, unchanged n = 58, increased n = 4) and by mRNA category (right; decreased n = 679, unchanged n = 10,112, increased n = 1,001). Violin plots with box plots showing median and interquartile range. Genes with unchanged promoters have lower baseline accessibility than genes with reduced promoters (Wilcoxon p = 1.7 × 10^−29^).

**Table 7.**
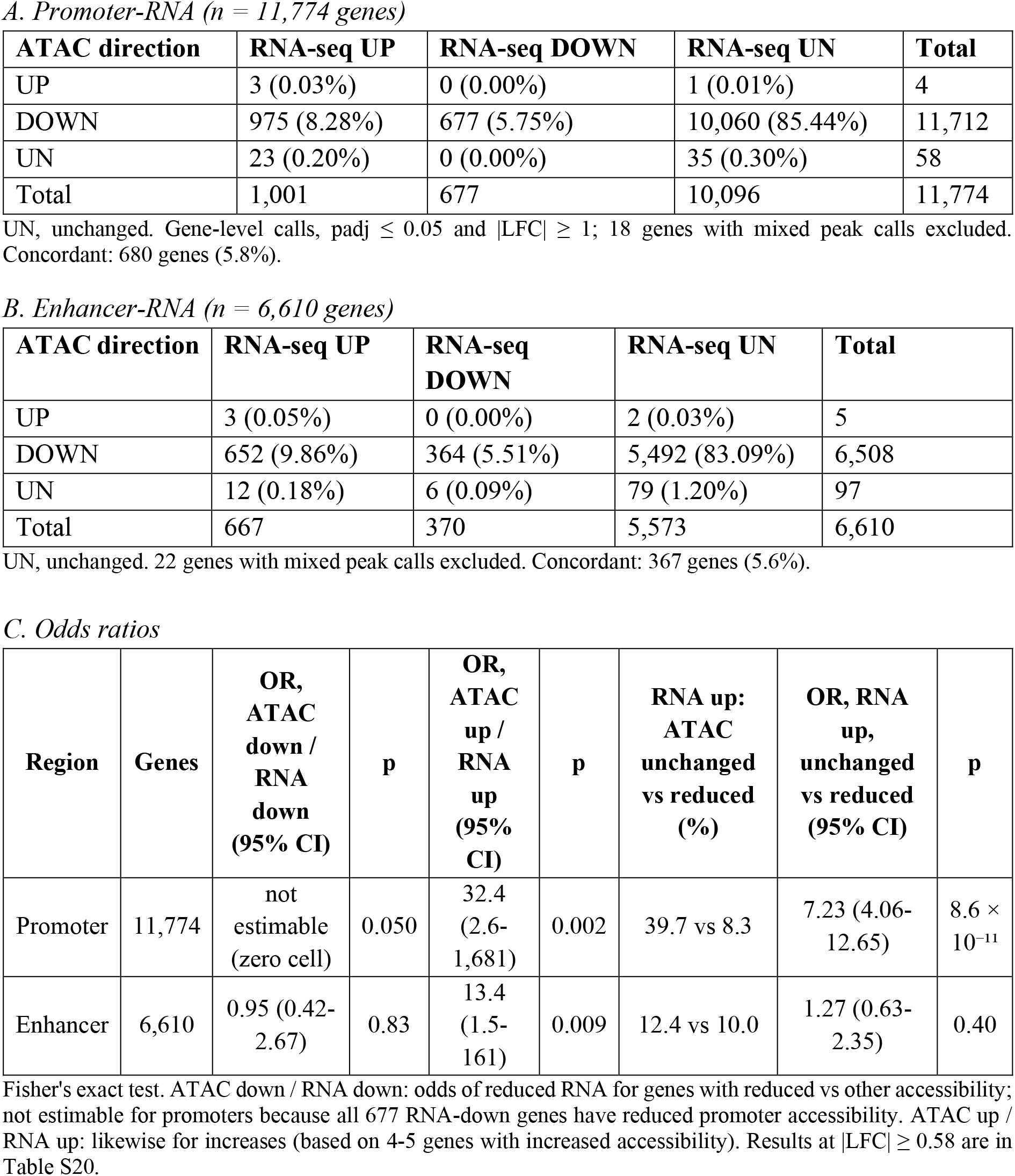
Contingency analysis of directional changes in chromatin accessibility and gene expression.

| *A. Promoter-RNA (n = 11,774 genes)* |  |  |  |  |  |  |  |  |
| **ATAC direction** | **RNA-seq UP** | **RNA-seq DOWN** | **RNA-seq UN** | **Total** |  |  |  |  |
|---|---|---|---|---|---|---|---|---|
| UP | 3 (0.03%) | 0 (0.00%) | 1 (0.01%) | 4 |  |  |  |  |
| DOWN | 975 (8.28%) | 677 (5.75%) | 10,060 (85.44%) | 11,712 |  |  |  |  |
| UN | 23 (0.20%) | 0 (0.00%) | 35 (0.30%) | 58 |  |  |  |  |
| Total | 1,001 | 677 | 10,096 | 11,774 |  |  |  |  |
| *B. Enhancer-RNA (n = 6,610 genes)* |  |  |  |  |  |  |  |  |
| **ATAC direction** | **RNA-seq UP** | **RNA-seq DOWN** | **RNA-seq UN** | **Total** |  |  |  |  |
| UP | 3 (0.05%) | 0 (0.00%) | 2 (0.03%) | 5 |  |  |  |  |
| DOWN | 652 (9.86%) | 364 (5.51%) | 5,492 (83.09%) | 6,508 |  |  |  |  |
| UN | 12 (0.18%) | 6 (0.09%) | 79 (1.20%) | 97 |  |  |  |  |
| Total | 667 | 370 | 5,573 | 6,610 |  |  |  |  |
| *C. Odds ratios* |  |  |  |  |  |  |  |  |
| **Region** | **Genes** | **OR, ATAC down / RNA down (95% CI)** | **p** | **OR, ATAC up / RNA up (95% CI)** | **p** | **RNA up: ATAC unchanged vs reduced (%)** | **OR, RNA up, unchanged vs reduced (95% CI)** | **p** |
| Promoter | 11,774 | not estimable (zero cell) | 0.050 | 32.4 (2.6-1,681) | 0.002 | 39.7 vs 8.3 | 7.23 (4.06-12.65) | 8.6 × 10<sup>−11</sup> |
| Enhancer | 6,610 | 0.95 (0.42-2.67) | 0.83 | 13.4 (1.5-161) | 0.009 | 12.4 vs 10.0 | 1.27 (0.63-2.35) | 0.40 |

UN, unchanged. Gene-level calls, padj ≤ 0.05 and |LFC| ≥ 1; 18 genes with mixed peak calls excluded. Concordant: 680 genes (5.8%).
UN, unchanged. 22 genes with mixed peak calls excluded. Concordant: 367 genes (5.6%).
Fisher's exact test. ATAC down / RNA down: odds of reduced RNA for genes with reduced vs other accessibility; not estimable for promoters because all 677 RNA-down genes have reduced promoter accessibility. ATAC up / RNA up: likewise for increases (based on 4-5 genes with increased accessibility). Results at |LFC| ≥ 0.58 are in Table S20.

### Accessibility-stratified analysis reveals selective expression responses

Genes were first stratified by the direction of promoter accessibility change. Genes whose promoters retained accessibility were more often up-regulated: 39.7% of the 58 genes with unchanged promoter accessibility showed increased mRNA abundance, compared with 8.3% of genes with reduced accessibility (odds ratio 7.23, 95% CI 4.06-12.65; p = 8.6 × 10^−11^; **Table 7**), and their median RNA LFC was positive (0.61 vs -0.06; Supplementary **Figure S12**). Genes with unchanged promoters were, however, few and had low baseline accessibility in AG cells (median log2 normalised fragments 5.5 vs 7.8; p = 1.7 × 10^−29^; **Figure 7C**), and accessibility at the most accessible promoters fell the most (Spearman ρ = -0.55 between baseline accessibility and promoter LFC).

We therefore tested the relationship across all genes, ranked into quintiles of promoter accessibility change (**Figure 7B**). mRNA changes increased steadily from the quintile with the largest loss of promoter accessibility to the quintile with the smallest loss (median RNA LFC -0.20 to 0.15). The proportion of up-regulated genes rose from 4.0% to 17.5% (trend p = 3.5 × 10^−68^; odds ratio 5.05 for the fifth vs the first quintile), and the proportion of down-regulated genes fell from 9.2% to 4.0% (trend p = 1.0 × 10^−13^). The gradient was independent of baseline promoter accessibility (adjusted change in RNA LFC per quintile 0.118, p = 1.8 × 10^−88^; adjusted odds ratio for up-regulation 1.40 per quintile, 95% CI 1.32-1.48). A weaker gradient was observed for enhancer accessibility (odds ratio 2.11 for the fifth vs the first quintile; Supplementary **Figure S13**).

Multivariable regression for genes with matched promoter and enhancer data (n = 5,805) confirmed that promoters were approximately four-fold more predictive than enhancers (standardised β = 0.240 vs 0.059; **Table 8**). Adding baseline promoter accessibility did not improve the model (β = 0.005, p = 0.75), and the combined model explained only ∼6% of the variance in mRNA-abundance changes (R^2^ = 0.064), indicating that chromatin accessibility accounts for only a limited fraction of gene-level expression changes during starvation.

**Table 8.** Multivariable regression of gene expression on promoter and enhancer accessibility.

| **Model** | **Predictor** | **β (raw)** | **β (standardised)** | **SE** | **t** | **p** |
|---|---|---|---|---|---|---|
| 1 | Intercept | 1.246 | - | 0.064 | 19.35 | 7.5 × 10<sup>−81</sup> |
| 1 | Promoter LFC | 0.443 | 0.240 | 0.024 | 18.82 | 1.0 × 10<sup>−76</sup> |
| 1 | Enhancer LFC | 0.085 | 0.059 | 0.018 | 4.61 | 4.1 × 10<sup>−6</sup> |
| 2 | Intercept | 1.231 | - | 0.079 | 15.51 | 3.6 × 10<sup>−53</sup> |
| 2 | Promoter LFC | 0.448 | 0.243 | 0.029 | 15.70 | 1.9 × 10<sup>−54</sup> |
| 2 | Enhancer LFC | 0.084 | 0.059 | 0.018 | 4.57 | 5.0 × 10<sup>−6</sup> |
| 2 | Baseline AG promoter accessibility | 0.003 | 0.005 | 0.010 | 0.32 | 0.75 |
| 3 | Intercept | 0.995 | - | 0.046 | 21.41 | 8.2 × 10<sup>−100</sup> |
| 3 | Promoter LFC | 0.452 | 0.259 | 0.019 | 24.13 | 1.3 × 10<sup>−125</sup> |
| 3 | Baseline AG promoter accessibility | 0.008 | 0.013 | 0.007 | 1.23 | 0.22 |
| **Model** | **Formula** | **n (genes)** | **R²** | **Adjusted R²** | **F (df)** | **p** |
| 1 | RNA ~ promoter + enhancer | 5,805 | 0.064 | 0.064 | 199.95 (2, 5802) | 1.1 × 10<sup>−84</sup> |
| 2 | RNA ~ promoter + enhancer + baseline | 5,805 | 0.064 | 0.064 | 133.31 (3, 5801) | 1.6 × 10<sup>−83</sup> |
| 3 | RNA ~ promoter + baseline (all promoter genes) | 11,792 | 0.063 | 0.063 | 399.10 (2, 11789) | 1.9 × 10<sup>−168</sup> |

RNA and ATAC values are gene-level apeglm-shrunken LFC. Baseline AG promoter accessibility, log2(median normalised fragments + 1) of the gene's promoter peaks in AG cells. Standardised coefficients allow comparison of predictors.

Together, these analyses indicate that, against a background of genome-wide loss, increased mRNA abundance occurs preferentially at genes whose promoters retain relatively more accessibility, whereas genes with the largest losses show a modest shift towards reduced expression.

### Classification of regulatory response patterns

Expressed genes with promoter or enhancer accessibility data (n = 12,619) were classified according to the direction of their RNA-seq and ATAC-seq changes, combining promoter and enhancer calls (**Figure 7A**). The largest category (85.0%) consisted of genes with reduced accessibility and no detectable change in mRNA abundance. The second-largest category (8.5%) comprised genes with increased mRNA abundance despite reduced accessibility, followed by concordant decreases (5.8%). Changes in mRNA abundance without a change in accessibility were rare (0.1%; 12 genes), as were concordant increases (2 genes), and no gene showed reduced mRNA abundance with increased accessibility. Because almost all genes lost accessibility, this distribution is shaped by the global reduction; it nevertheless reinforces the conclusion that starvation-induced changes in chromatin accessibility are largely not accompanied by detectable changes in mRNA abundance, and it identifies a sizeable group of genes that are up-regulated despite reduced accessibility.

Applying stringent criteria (padj ≤ 0.01 and |LFC| ≥ 2 in RNA-seq and in at least one promoter or enhancer peak, with the same direction) identified 99 genes with coordinated accessibility-expression changes, all of which showed concordant decreases (Supplementary **Table S6**).

### Gene-level integration reveals expression-chromatin discordance in adaptive programmes

To examine gene-level changes within functionally coherent programmes, we integrated expression and accessibility data for leading-edge genes from enriched pathways (Supplementary **Table S7**; **Figure 8A, B**). In every programme, 95-100% of leading-edge genes with a promoter peak showed significantly reduced promoter accessibility, irrespective of the direction of their mRNA change. The programmes differed instead in the magnitude of the loss, and this analysis revealed two distinct regulatory patterns.

**Figure 8.**
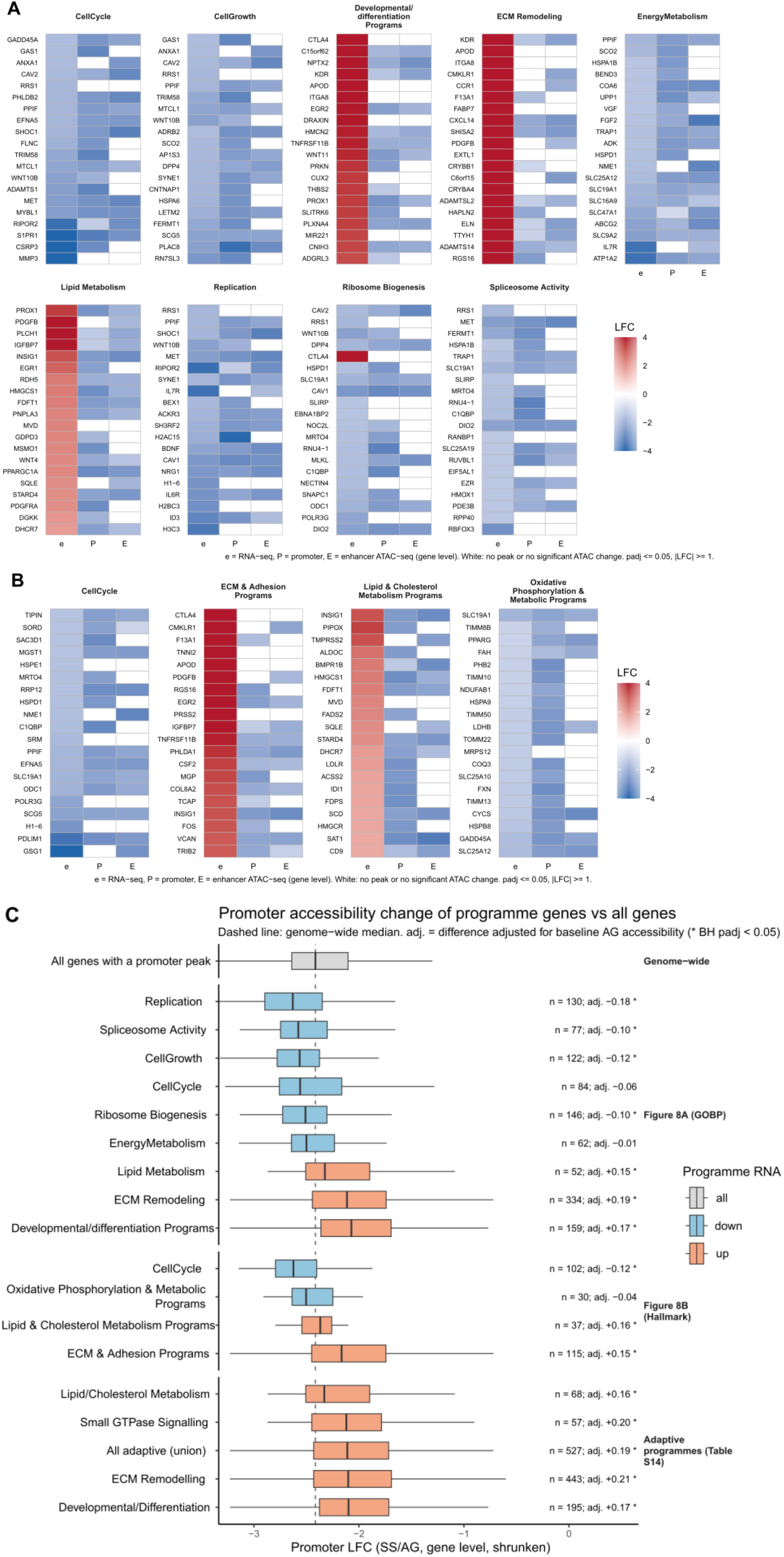
Integrated transcriptomic and chromatin-accessibility profiles of leading-edge genes from functionally clustered pathways. (A, B) Heatmaps of gene-expression and chromatin-accessibility changes between AG and SS cells for leading-edge genes of functionally clustered pathway categories from (A) GO:BP and (B) Hallmark collections. For each gene, three measurements are shown: RNA LFC (e), promoter ATAC-seq LFC (P) and enhancer ATAC-seq LFC (E; gene-level values as in Figure 6). The top 20 leading-edge genes per category are shown, ranked by absolute RNA fold change. Colour scale represents LFC from -4 to +4. White cells indicate no peak or no significant ATAC-seq change. Thresholds: padj ≤ 0.05, |LFC| ≥ 1. (C) Promoter accessibility change (gene-level, apeglm-shrunken LFC) of leading-edge genes of each category in (A, B) and of the four adaptive programmes (Table S7) compared with all genes with a promoter peak (n = 11,792; grey). Box plots coloured by the direction of the programme’s mRNA change (down, blue; up, orange). The dashed line marks the genome-wide median (-2.41). For each set, n and the difference from all other genes adjusted for baseline AG promoter accessibility are shown; * Benjamini-Hochberg padj < 0.05 (linear regression of promoter LFC on set membership and baseline accessibility).

First, genes driving down-regulated biosynthetic and proliferative programmes were suppressed together with a strong reduction in promoter accessibility (Supplementary **Table S7**), equal to or greater than the genome-wide reduction (**Figure 8C**).

Second, and more strikingly, genes within the four up-regulated adaptive programmes (ECM remodelling, lipid/cholesterol metabolism, small GTPase signalling and developmental/differentiation programmes; Supplementary **Table S7**) displayed pronounced expression-accessibility discordance. Their mRNA levels increased (Supplementary **Table S7**), although ∼98% of those with a promoter peak showed significantly reduced promoter accessibility and enhancer accessibility was similarly reduced (**Figure 8A, B**). Their loss of promoter accessibility was, however, smaller than that of other genes (**Figure 8C**). The difference persisted after adjustment for baseline promoter accessibility in AG cells and was observed in each of the four programmes.

Thus, adaptive-programme genes were up-regulated despite reduced promoter signal per mapped fragment, but their promoters retained about 1.2-fold more accessibility than those of other genes. This comparison between groups of genes does not depend on the normalisation, and relative to the remaining peak signal it corresponds to preferential preservation of promoter accessibility. The pattern is in line with the genome-wide gradient between promoter accessibility change and mRNA change (**Figure 7B**) and suggests selective prioritisation of survival-critical functions through mechanisms that operate largely independently of bulk chromatin accessibility.

### Nascent and total RNA measurements at selected loci agree with RNA-seq and confirm discordance between transcription and chromatin accessibility at adaptive genes

Genes in the upregulated adaptive programmes showed increased RNA abundance despite reduced promoter and enhancer accessibility (**Figure 8**). To test whether this reflects increased RNA production, we pulse-labelled AG and SS cells with 5-ethynyluridine (5-EU) for 30 min and quantified labelled (nascent) and total RNA by RT-qPCR. The panel comprised 17 leading-edge genes from adaptive and biosynthetic programmes, plus UBXN2A as an unchanged control (Supplementary **Tables S7 and S17**).

All 17 regulated genes showed significant total RNA changes in the RNA-seq direction, whereas UBXN2A showed no significant change (**Figure 9A**). Total RNA LFCs correlated with RNA-seq across all 18 genes (Spearman ρ = 0.77; **Figure 9C**).

**Figure 9.**
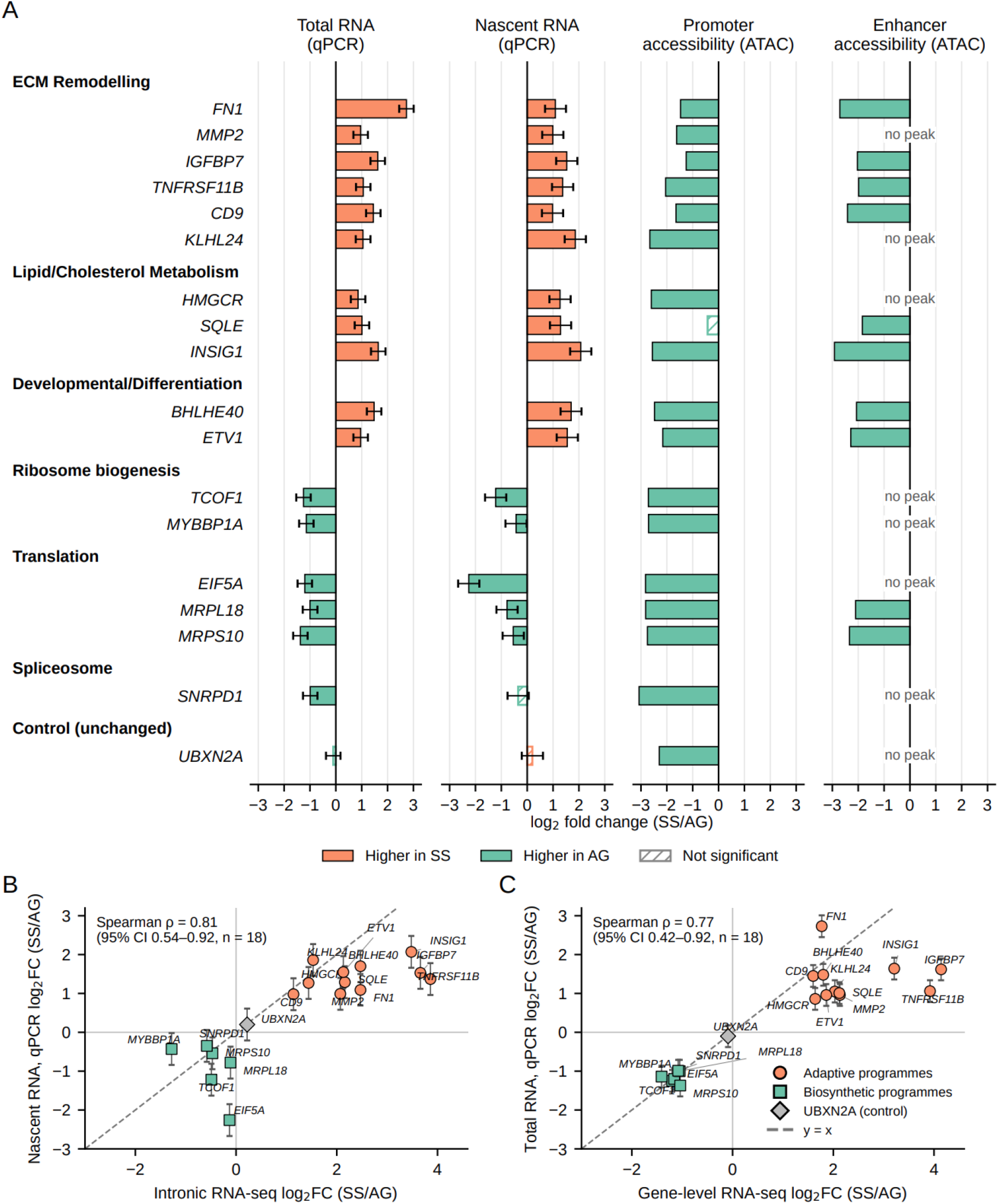
Nascent and total RNA changes at selected adaptive and biosynthetic genes in starved U2OS cells compared with RNA-seq and ATAC-seq. (A) LFCs (SS/AG) in total RNA and 5-EU-labelled nascent RNA measured by RT-qPCR, shown alongside changes in promoter and enhancer accessibility assigned to the same genes by ATAC-seq. The panel includes 11 leading-edge genes from upregulated adaptive programmes, six leading-edge genes from downregulated biosynthetic programmes (Table S7) and the unchanged control UBXN2A. Genes are grouped by programme. Orange bars indicate higher values in starved (SS) cells and green bars higher values in actively growing (AG) cells; hatched bars indicate non-significant changes. qPCR values are means from three independent experiments, with error bars representing 95% confidence intervals. Total RNA was normalised to 28S rRNA, and nascent RNA was corrected for spike-in recovery. Significance was defined as q < 0.05 using a limma moderated paired test with Benjamini-Hochberg correction. ATAC-seq values are gene-level LFCs (DESeq2, library-size normalisation; n = 3 per condition): promoter, median of the gene’s promoter peaks; enhancer, GeneHancer-score-weighted mean of linked enhancer peaks. ATAC-seq changes were called significant at padj ≤ 0.05 and |LFC| ≥ 1 for at least one of the gene’s peaks. “No peak” indicates the absence of a linked enhancer peak. (B) Nascent RNA changes measured by RT-qPCR compared with intronic RNA-seq changes calculated from intronic counts (DESeq2 with apeglm-shrunken LFCs; n = 2 per condition). (C) Total RNA changes measured by RT-qPCR compared with gene-level RNA-seq changes (DESeq2 Wald test; n = 2 per condition). In (B) and (C), error bars represent 95% confidence intervals for the qPCR estimates. Circles denote adaptive-programme genes, squares biosynthetic-programme genes, and the diamond UBXN2A. The dashed line indicates y = x. Spearman correlation coefficients were calculated across all 18 genes, with 95% confidence intervals obtained from 10,000 bootstrap resamples of genes. Values for individual genes are provided in Supplementary Table S17.

Nascent RNA increased for all 11 genes representing the ECM remodelling, lipid/cholesterol metabolism and developmental/differentiation programmes and decreased for all five ribosome biogenesis and translation genes (**Figure 9A**). The decrease for the spliceosome gene SNRPD1 was not significant. Nascent RNA changes correlated with intronic RNA-seq changes across all 18 genes (Spearman ρ = 0.81; **Figure 9B**).

All 11 adaptive genes had a promoter-associated ATAC-seq peak, and ten showed significantly reduced promoter accessibility; the exception, SQLE, showed a small, non-significant decrease (LFC -0.42). Eight had GeneHancer-linked enhancer peaks, and enhancer accessibility was significantly reduced for all eight, including SQLE (**Figure 9A**). Every adaptive gene therefore showed reduced accessibility at its promoter, its linked enhancers or both. Promoter accessibility was similarly reduced at all six biosynthetic genes and at the unchanged control UBXN2A (LFC -2.29). Thus, increased RNA production occurred despite reduced Tn5 accessibility at these *loci*.

The panel included seven genes with an EISA post-transcriptional call: three stabilisation candidates (IGFBP7, CD9 and KLHL24) and four destabilisation candidates (FN1, TCOF1, EIF5A and MRPL18), all with modest interaction terms (|Dex - Din| 0.54 - 0.82). A change in transcript stability would alter the nascent-to-total ratio; a negative ratio LFC indicates that total RNA increased more, or decreased less, than RNA production. FN1 was the only gene with a significant change in this ratio (LFC −1.73, q = 0.017), which is opposite in direction to its EISA destabilisation call. None of the other six EISA candidates showed a significant ratio change.

Nascent 28S rRNA signal decreased 7.4-fold, consistent with reduced rRNA synthesis in starved cells, whereas total 28S rRNA per µg of RNA, used to normalise total RNA, decreased by only 10%, suggesting limited effect of starvation on the total rRNA pool. Notably, spike-in recovery did not differ significantly between conditions (Supplementary **Table S18**), and all retained standard curves and gene assays met the predefined performance criteria after one pooled-slope refit (28S, experiment 1) and the exclusion of one assay (EFEMP2, efficiency 83.4%, Supplementary **Table S19**). The spike-in does not correct for differences in cellular 5-EU uptake or incorporation, which could affect the magnitude of nascent RNA changes. A global difference in labelling would, however, shift all nascent measurements in the same direction and cannot account for the opposite changes observed in the same samples, in which adaptive transcripts increased while biosynthetic transcripts and 28S rRNA decreased.

## Discussion

### Limited ATAC-RNA coupling during starvation

The present study provides a comprehensive characterisation of the relationship between chromatin accessibility and gene expression during severe nutrient deprivation. Our integrated analysis reveals a striking asymmetry. Starvation induces a widespread reduction in chromatin accessibility, with 99% of significantly changed peaks showing reduced accessibility. In contrast, the gene-expression response is more limited and selective: only 12.4% of genes are differentially expressed, and 64% of these show increased rather than decreased mRNA abundance (**Table 4**). This contrast between global loss of chromatin accessibility and selective increases in mRNA abundance is a central finding of the study and has important implications for interpreting chromatin accessibility during stress-responsive gene regulation.

Consistent with this asymmetry, accessibility and expression changes were only modestly correlated at promoters and enhancers (r = 0.25 and r = 0.10, respectively; **Figure 6**), and only about 6% of genes showed concordant directional changes (**Figure 7A**). These values are consistent with previous reports of modest ATAC-RNA coupling across perturbations and cell types, including promoter-level correlations reported by Kiani et al. [9]. Thus, even under extensive chromatin remodelling, changes in accessibility provide only limited information about the direction and magnitude of mRNA abundance changes.

Within-condition analyses placed these cross-condition relationships in context. Promoter accessibility and steady-state expression were similarly modestly correlated in actively growing and starved cells (AG: r = 0.28; SS: r = 0.25; Supplementary **Table S8**), comparable to the cross-condition promoter correlation (r = 0.25; **Figure 6B**). Starvation therefore does not disrupt a previously strong accessibility-expression relationship. Rather, chromatin remodelling occurs within a regulatory framework in which promoter accessibility is already only modestly predictive of mRNA abundance.

This model provides a stringent test of ATAC-RNA coupling because the chromatin response is strongly directional, unlike perturbations that produce more balanced mixtures of accessibility gains and losses. Despite coordinated genome-wide accessibility loss, transcriptional responses remained selective and weakly associated with accessibility changes, with promoter and enhancer accessibility together explaining approximately 6% of the variance in mRNA abundance changes. The association that remained was graded rather than categorical: genes whose promoters lost the least accessibility were more often up-regulated than those that lost the most (17.5% vs 4.0%; **Figure 7B**), independently of their accessibility before starvation. Similar modest accessibility-expression relationships have been reported under oxidative stress, heat stress and other perturbations [10–12]. Targeted nascent RNA measurements further showed that this discordance extends to RNA production: at the adaptive-programme loci tested, nascent RNA increased despite reduced accessibility at the promoter, the linked enhancers or both (**Figure 9**). Together, these findings support a model in which chromatin accessibility acts as a permissive rather than deterministic component of transcriptional regulation during nutrient stress.

### Potential mechanisms underlying the genome-wide reduction in chromatin accessibility

The genome-wide reduction in ATAC-seq accessibility in starved cells raises an important interpretative question: whether it reflects a genuine biological decrease in Tn5 accessibility or a technical artefact of library preparation. Several lines of evidence discussed below argue strongly against a technical artefact, but, without a spike-in experiment, this possibility cannot be excluded completely.

Firstly, AG and SS libraries were prepared in parallel from equal numbers of nuclei and tagmented with the same amount of Tn5 from a single batch for the same time, so differences in transposition conditions do not explain the reduction. Furthermore, the number of PCR cycles during library pre-amplification was also equal. Secondly, the structure of the starved libraries is consistent with a biological reduction in Tn5 accessibility (see also Supplementary Data Section 1). Starved libraries retained characteristic ATAC-seq fragment architecture, including a shared short-fragment mode, nucleosome-associated fragment classes and detectable ∼10 bp Tn5 rotational periodicity (**Figure 1A**, Supplementary **Figure S2B-D, Table S1B**). Starvation reduced the NFR-sized fragment fraction while preserving mono-and dinucleosomal size classes, increasing the Mono/NFR and Di/NFR ratios and the median insert size, which is the opposite of the pattern expected from indiscriminate over-tagmentation. Mapping quality, proper pairing, duplication and estimated library size were comparable between conditions (Supplementary **Table S1A**), and the condition-specific structure was reproducible across replicates, which argues against under-tagmentation. Peak-to-background analysis showed that the dominant change was a loss of summit-centred signal, and that the smaller increases in flank and random-background signal matched those expected when fragments no longer captured in peaks are distributed over the rest of the genome (Supplementary **Figure S10, Table S12**). Thus, the starvation phenotype is best interpreted as a biological reduction in focal Tn5 accessibility rather than failure of library preparation.

The magnitude of this reduction requires one qualification. Library-size normalisation expresses the signal at each peak per mapped fragment. Because the libraries were amplified and sequenced to similar depth, the number of mapped fragments does not record the total yield of tagmented fragments per cell, and without an exogenous reference, such as spike-in chromatin, the approximately five-fold reduction per mapped fragment cannot be converted into an absolute reduction per cell. The reduction extended to housekeeping-gene promoters and distal CTCF-motif sites, which are usually regarded as constitutively accessible (**Figure 2C**), so these sites could not serve as internal references either. Normalisation to fragments in consensus peaks removes the overall reduction by construction and instead describes how the remaining peak-associated signal is distributed among regions (Supplementary **Table S3**). The two scales differ almost entirely by a constant. Correlations, rankings and comparisons between groups of genes therefore do not depend on the choice, whereas the numbers of gained and lost peaks and the directional classification of genes are reported on the per-fragment scale. The most parsimonious biological explanation is compaction-associated chromatin reorganisation. Global chromatin condensation is a conserved response to nutrient limitation and related stress states, including quiescence in *Saccharomyces cerevisiae* [14, 15], starvation in *Caenorhabditis elegans* [16], and arginine starvation or transient ischaemia in mammalian cells [17, 18]. Our results are compatible with this framework: accessibility loss was strongly directional, affected promoters, enhancers and constitutively accessible sites alike, and produced very few starvation-specific peaks, suggesting that starvation preserves a subset of pre-existing regulatory elements within a globally less accessible chromatin environment.

However, ATAC-seq alone cannot formally establish physical chromatin compaction, and other biological mechanisms may contribute. One plausible mechanism is altered linker histone H1 behaviour. H1 mobility is reduced and chromatin retention increased under nutrient deprivation, and H1 binds preferentially at nucleosomal positions targeted by Tn5 [18, 61]. Because H1 can suppress nucleosome unwrapping without directly excluding transcription-factor binding [62], a locus could register as less Tn5-accessible while remaining transcriptionally competent. A more speculative possibility is that starvation alters nuclear envelope or nuclear pore properties, for example through stress-responsive regulation of nuclear pore composition [63], although a direct link to Tn5 accessibility remains untested.

A distinct alternative is local or global chromatin decompaction, but this is unlikely to explain the observed library structure. Stress-induced chromatin opening is well documented, especially during heat shock, where transcriptional activation can involve local nucleosome loss, histone displacement and three-dimensional genome reorganisation [64–67]. Local accessibility gains have also been reported in Drosophila and human heat-shock models [19, 20]. Nevertheless, a global decompaction model would predict increased NFR-sized fragments, preserved or increased FRiP and TSS enrichment, and widespread formation of starvation-specific peaks. We observe the opposite pattern. Taken together, the fragment architecture, retained Tn5 periodicity, peak-centred signal loss, loss at constitutively accessible sites and peak-provenance analysis support a genuine genome-wide reduction in Tn5 accessibility, with compaction-associated chromatin reorganisation as the most likely, but not exclusive, mechanistic explanation.

### Transcriptional and post-transcriptional contributions to the starvation response

A key question for interpreting ATAC-RNA coupling is how much of the observed change in steady-state mRNA abundance reflects altered RNA production rather than transcript stability or RNA processing. Nascent-transcription assays such as PRO-seq, GRO-seq, SLAM-seq or related approaches provide the most direct readout of RNA synthesis, but they do not replace conventional steady-state RNA-seq, which captures the accumulated mRNA pool used for expression and pathway-level analyses. In the absence of genome-wide paired nascent and steady-state data, exon-intron concordance, EISA and intron-retention sensitivity analyses provide an indirect genome-wide assessment, which we complemented with direct measurements of nascent RNA at selected *loci* (**Figure 9**).

We observed an increase in the nascent RNA levels for all eleven adaptive-programme genes tested and decrease for the ribosome biogenesis and translation genes, showing that starvation combines reduced production of growth-related RNAs with induction of selected adaptive transcripts, and that reduced Tn5 accessibility does not preclude transcriptional induction at these *loci*. Nascent RNA changes also tracked intronic RNA-seq changes across the panel, supporting the use of intronic reads as a proxy for RNA production and, by extension, the genome-wide exon-intron concordance (Supplementary **Figure S8**). Total RNA measured by qPCR reproduced the direction of all RNA-seq changes and their ranking across genes, although qPCR effect sizes were generally smaller (Supplementary **Table S17**).

The 28S rRNA measurements provide an internal positive control. Nascent 28S rRNA decreased 7.4-fold, whereas the mature 28S pool changed only modestly. This is consistent with the downregulated ribosome biogenesis programme (**Figure 5**) and with the established repression of RNA polymerase I transcription under nutrient limitation, mediated in part by mTOR-dependent inactivation of the Pol I initiation factor TIF-IA [68] and with our previous observations under these starvation conditions [22].

Exon-intron concordance and EISA suggested a selective post-transcriptional component rather than a transcriptome-wide alternative explanation (Supplementary **Figures S8, S9**). At the stringent threshold, 542 genes, representing 5.0% of tested genes, showed a significant interaction term, comprising 437 stabilisation and 105 destabilisation candidates. The stabilisation class was dominated by genes whose intronic signal declined or remained unchanged, the pattern expected when production falls while the mature transcript pool persists, rather than one indicating selective transcript protection. Explaining increased steady-state abundance by stabilisation alone would also require unchanged or reduced transcriptional output, whereas increased exonic abundance was generally accompanied by increased intronic signal, and only around one in nine genes with significantly increased exonic abundance was a stabilisation candidate. Transcript-specific stabilisation does occur during nutrient stress, as illustrated by AUF1-dependent stabilisation of FST mRNA during glucose deprivation [69], but such examples represent selective control rather than a transcriptome-wide mechanism.

A related concern is that reduced expression of spliceosome-related genes during starvation (**Figure 5**) could generate apparent post-transcriptional signals through intron retention. However, exon-intron concordance remained strong after excluding genes most vulnerable to intron-retention artefacts, indicating that the dominant signal is not driven by splicing-complexity effects. Moreover, intron retention in mammalian cells is often regulated and transcript-selective, as illustrated by detained introns and stress-responsive kinase control of SR splicing factors [70, 71].

The nascent-to-total measurements did not independently confirm the seven EISA calls tested. Six candidates had no significant ratio change, while FN1 showed an apparent opposite-direction change requiring independent confirmation. The modest EISA interaction effects (|Dex - Din| 0.54-0.82), limited precision of ratio estimates and differences between the RNA populations measured may contribute to this disagreement. Intronic abundance depends on splicing and intron turnover as well as production, and the nascent-to-total ratio does not directly measure RNA stability. Gene-level EISA calls should therefore be regarded as candidates, and direct measurements of RNA decay will be needed to establish whether transcript stability is altered.

Together, these analyses indicate that changes in RNA production make the major contribution to starvation-associated changes in RNA abundance, whereas the post-transcriptional contribution suggested by EISA is selective and was not confirmed at the *loci* tested. This pattern is consistent with mammalian stress responses, in which post-transcriptional regulation often provides transcript-specific control superimposed on broader transcriptional programmes [60]. The nascent RNA measurements have limitations: the genes were selected from RNA-seq results, so agreement in total RNA is partly expected, and the panel comprised only 18 *loci*. Genome-wide nascent RNA profiling and direct measurement of RNA decay in starved cells will be required to establish how generally these conclusions apply.

### Mechanistic implications of ATAC-RNA discordance during starvation

A central question raised by limited ATAC-RNA coupling is how specific genes, particularly leading-edge genes of the four upregulated adaptive programmes (**Figure 8**), achieve increased expression despite reduced promoter and enhancer accessibility. Several non-mutually exclusive mechanisms may contribute, independently of the post-transcriptional component addressed above.

First, the chromatin landscape established before stress may pre-configure which genes retain regulatory potential during starvation. Peak-provenance analysis showed that most accessible regions detected in starved cells were not formed de novo and that approximately 95% of starvation-accessible peaks corresponded to regions already present under growth conditions (**Figure 1H**, **Table 1**). Starvation therefore appears to act primarily through selective retention of pre-existing regulatory elements. In line with this, genes whose promoters retained relatively more accessibility were more often up-regulated, although the effect was modest (Figure 7B). This is consistent with reports that heat-shock transcriptional responses are substantially pre-wired by promoter and enhancer architecture, with rapidly induced genes often possessing pre-loaded polymerase and accessible chromatin before stress [72].

Second, transcriptional machinery already engaged at adaptive-programme genes may continue to operate even as bulk accessibility declines. Paused RNA polymerase II complexes, present at many mammalian promoters, can be rapidly released into productive elongation without requiring de novo chromatin opening [73]. Similarly, pioneer transcription factors can engage nucleosomal DNA directly, and their occupancy can precede, and is not strictly dependent on, detectable accessibility changes [74, 75]. Such mechanisms would be compatible with reduced bulk Tn5 signal at transcriptionally responsive *loci*.

Third, accessibility loss and mRNA changes may not be temporally synchronous. Our design captures a single 18-hour timepoint and cannot resolve the order of chromatin and transcriptional changes. Finally, bulk ATAC-seq and RNA-seq average over the full cell population and may obscure cell-to-cell heterogeneity in chromatin-transcription coupling. Single-cell multi-omic approaches capable of jointly profiling accessibility and expression in the same cell would directly test this possibility. These mechanisms are not mutually exclusive and may act in combination to sustain expression of survival-critical genes within a less accessible chromatin landscape.

### An integrated model of starvation-adaptive regulatory programmes

Starvation induces profound pathway-level regulatory reprogramming, characterised by a clear asymmetry: global biosynthetic and proliferative programmes are broadly suppressed in parallel with reduced chromatin accessibility, whereas a restricted subset of adaptive pathways becomes upregulated despite the globally less accessible chromatin environment (**Figures 5, 8 and Table S5**). These include small GTPase signalling, extracellular matrix organisation, lipid and cholesterol metabolism, and developmental pathways. Although these pathways are characteristically enriched in embryonic and developmental contexts, where morphogenetic remodelling and cellular plasticity are prominent [76–78], their co-activation under starvation does not necessarily imply reversion to a developmental regulatory state. Instead, it may reflect convergent deployment of conserved signalling modules serving distinct adaptive purposes. Regulator and chromatin-state enrichment analyses add a candidate mechanistic layer to this model. The enrichment of PRC2/PRC1 targets and H3K27me3-associated gene sets among RNA-up genes, particularly within the ECM Remodelling and Developmental/Differentiation programmes (Supplementary **Tables S14 and S15**), is consistent with the possibility that starvation-responsive RNA-abundance increases involve genes embedded in Polycomb-associated regulatory contexts. This does not demonstrate loss of Polycomb occupancy or H3K27me3 in starved U2OS cells, but it provides a plausible framework for understanding why increased RNA abundance can occur without detectable gains in ATAC-seq accessibility: Polycomb-regulated promoters are not necessarily nucleosome-occluded, and relief or bypass of Polycomb-associated repression can alter transcriptional output without requiring proportional increases in bulk Tn5 accessibility [79, 80]. Conversely, enrichment of MYC/MAX targets among RNA-down genes is consistent with repression of a MYC-linked biosynthetic programme, in line with the established role of MYC in ribosome biogenesis, protein synthesis and metabolic growth control [81, 82]. Finally, programme-specific enrichment of SREBF1/2 and SMAD3/4 (supplementary **Table S14**) links the lipid/cholesterol and developmental/ECM-associated modules to sterol homeostasis and TGF-β/BMP signalling, respectively [83, 84]. These assignments should be interpreted as candidate regulatory hypotheses rather than resolved mechanisms, but they sharpen the proposed starvation-adaptive model by nominating specific factors and chromatin-state contexts for future perturbation studies. Strikingly, leading-edge genes within all upregulated programmes exhibited strong increases in mRNA abundance despite reduced accessibility at their promoters and enhancers, representing the most pronounced examples of ATAC-RNA discordance in our dataset (**Figure 8A, B**). Their promoters lost less accessibility than those of other genes (**Figure 8C**), so that, relative to the remaining peak signal, their promoter accessibility was preferentially preserved. This difference is, however, too small to account for the increases in mRNA abundance.

Restricting exon-intron concordance and EISA analyses to these genes showed that they provide the strongest evidence for transcriptional activation in the dataset: among exon-upregulated genes, adaptive-programme leading-edge genes were substantially more likely than the remaining exon-upregulated genes to show a significant concordant increase in intronic signal (66.2% versus 34.0%; odds ratio 3.80, Fisher’s exact p = 2.4 × 10^-41^; Supplementary **Figure S11A**). Direct measurements confirmed increased RNA production for all eleven adaptive-programme genes tested (**Figure 9**). The same leading-edge genes were also enriched for the EISA stabilisation class relative to background (Supplementary **Figure S11B**), raising the possibility of an additional post-transcriptional layer. However, none of the three adaptive-programme stabilisation candidates tested (IGFBP7, CD9 and KLHL24) showed a significant change in the nascent-to-total ratio, so this layer remains a possibility rather than an established feature of the adaptive response.

Together, these analyses provide direct evidence at selected *loci*, and indirect genome-wide evidence, that increased transcription contributes to the induction of the adaptive programmes. Thus, the adaptive programmes most clearly illustrate the central conclusion of our analysis: selective mRNA induction during starvation can occur in a less accessible chromatin environment.

We propose a hypothetical “starvation-adaptive state” integrating all four upregulated programmes (**Figure 10**). The coordinated upregulation of Rho/Rab/Rap GTPase regulators and effectors [85–87], ECM components and integrins suggests potential coupling between cytoskeletal dynamics and cell-matrix interactions, with Rap1 signalling providing an established link through integrin activation [88–91]. Concurrent activation of lipid and cholesterol biosynthesis, including SREBF2 targets and mevalonate pathway enzymes [92], may support membrane remodelling while providing isoprenoid substrates required for GTPase prenylation [93].

**Figure 10.**
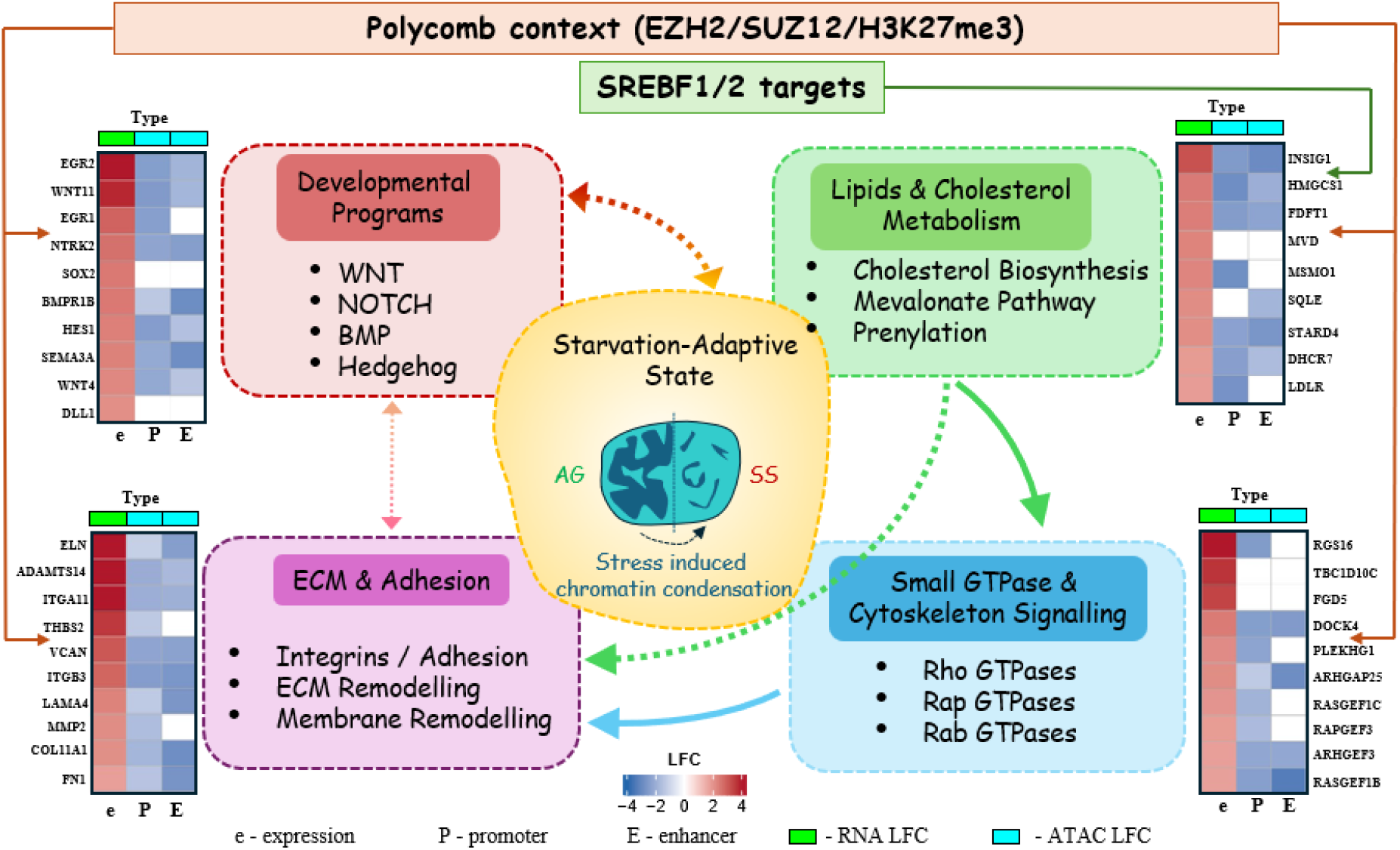
Integrated model of a hypothetical starvation-adaptive state. Schematic representation of four interconnected biological programmes upregulated in starved U2OS cells: developmental signalling, lipid and cholesterol metabolism, small GTPase and cytoskeleton signalling, and ECM and adhesion. The central schematic depicts a proposed starvation-adaptive state associated with reduced chromatin accessibility. Dashed arrows indicate proposed regulatory and functional relationships between programmes. The Polycomb-associated frame indicates enrichment of adaptive programme genes for EZH2/SUZ12 targets and H3K27me3-associated loci (Supplementary Tables S14 and S15). The SREBF1/2 banner indicates enrichment of the Lipid and Cholesterol Metabolism programme for SREBF1/2 targets (Supplementary Table S14). The mevalonate pathway may provide isoprenoid substrates required for GTPase prenylation, while GTPase signalling may support cytoskeletal dynamics, ECM engagement and macropinocytosis. Although several pathways overlap with embryonic or developmental signatures, their co-activation under starvation is interpreted as convergent deployment of conserved adaptive modules rather than reversion to a developmental state. Heatmaps show LFC values for representative genes in each programme: RNA expression (e), promoter accessibility (P) and enhancer accessibility (E), as in Figure 8. Colour scale ranges from -4 to +4; white cells indicate no peak or no significant ATAC-seq change. The recurrent pattern of increased RNA abundance with reduced promoter and enhancer accessibility illustrates ATAC-RNA discordance across the four adaptive programmes. Thresholds: |LFC| ≥ 1, padj ≤ 0.05.

ECM remodelling during starvation is consistent with recent reports showing that ECM components can serve as nutrient reservoirs through macropinocytic uptake [57], and ECM gene induction has also been observed during serum starvation in fibroblasts [58]. The additional engagement of developmental signalling pathways, including WNT, NOTCH and BMP, may reflect stress-induced cellular plasticity rather than lineage commitment [94–99]. This model should be viewed as hypothesis-generating rather than mechanistically resolved. The proposed relationships and their consequences for membrane dynamics, ECM interactions or nutrient acquisition remain to be tested through targeted genetic perturbations and functional assays.

## Conclusion

This study shows that severe nutrient deprivation produces a genome-wide reduction in Tn5 accessibility while allowing selective increases in mRNA abundance. The accessibility loss is strongly directional, affects promoters, enhancers and constitutively accessible sites, and is supported by fragment architecture, peak-centred signal and peak-provenance analyses as a biological reduction in focal Tn5 accessibility rather than a library-preparation artefact, although its exact magnitude cannot be established without an exogenous reference. The most likely interpretation is compaction-associated chromatin reorganisation, although ATAC-seq alone cannot formally establish physical chromatin compaction.

Integrated ATAC-seq and RNA-seq analyses show that chromatin accessibility and mRNA abundance remain only modestly coupled during starvation. Exon-intron concordance and EISA indicate that changes in RNA production are the major contributor to altered RNA abundance, and direct measurement of nascent RNA at selected loci confirmed increased RNA production of adaptive-programme genes despite reduced promoter and enhancer accessibility, alongside reduced production of biosynthetic transcripts and rRNA. A post-transcriptional component suggested by EISA for a subset of genes was not confirmed at the loci tested. Together, these findings support a model in which selective transcriptional induction can occur within a globally less accessible chromatin environment. The proposed starvation-adaptive state, involving extracellular matrix remodelling, lipid and cholesterol metabolism, small GTPase signalling and developmental pathways, should be viewed as a hypothesis-generating framework requiring genome-wide nascent RNA profiling, single-cell multi-omics and functional perturbation for direct mechanistic testing.

## Supporting information

Abbreviation list

## List of abbreviations

is available in Supplementary information.

## Declarations

### Ethics approval and consent to participate

Not applicable.

### Consent for publication

Not applicable.

### Availability of data and materials

The datasets used and/or analysed during the current study are available at the GEO repository, accession number GSE327351.

### Competing interests

The authors declare that they have no competing interests.

## Funding

This work was not funded.

### Authors’ contributions

XQ and SAR performed all wet-lab experiments, conducted the initial data analysis, and contributed to writing the manuscript. SM analysed the data and contributed to writing the manuscript. KP conceived the study, analysed the data, and was the major contributor to writing the manuscript.

## Acknowledgements

The authors acknowledge the staff of the Genomics CTU (Courtney Ward and Sara El Jadid), Faculty of Medicine, Health and Life Sciences, Queen’s University Belfast, for their assistance with RNA- and ATAC-sequencing and support with initial data analysis. The authors also thank former KP student Daniella Ramalho McCourt for her contribution to ATAC-seq data analysis, Dr Fuquan Liu for kindly providing the pSpike plasmid, Professor Harry Hardiman (QUB) for assistance with data analysis and critical reading of the manuscript, and Professor Chris Creevey (QUB) for critical reading of the manuscript. Xiyuan Qi acknowledges the Queen’s-China Scholarship Council for financial support during her PhD.

## Notes

### Competing Interest Statement

The authors have declared no competing interest.

