## Supplementary figures for "Genome-wide loss of chromatin accessibility coexists with selectively elevated RNA abundance in nutrient-starved U2OS cells"

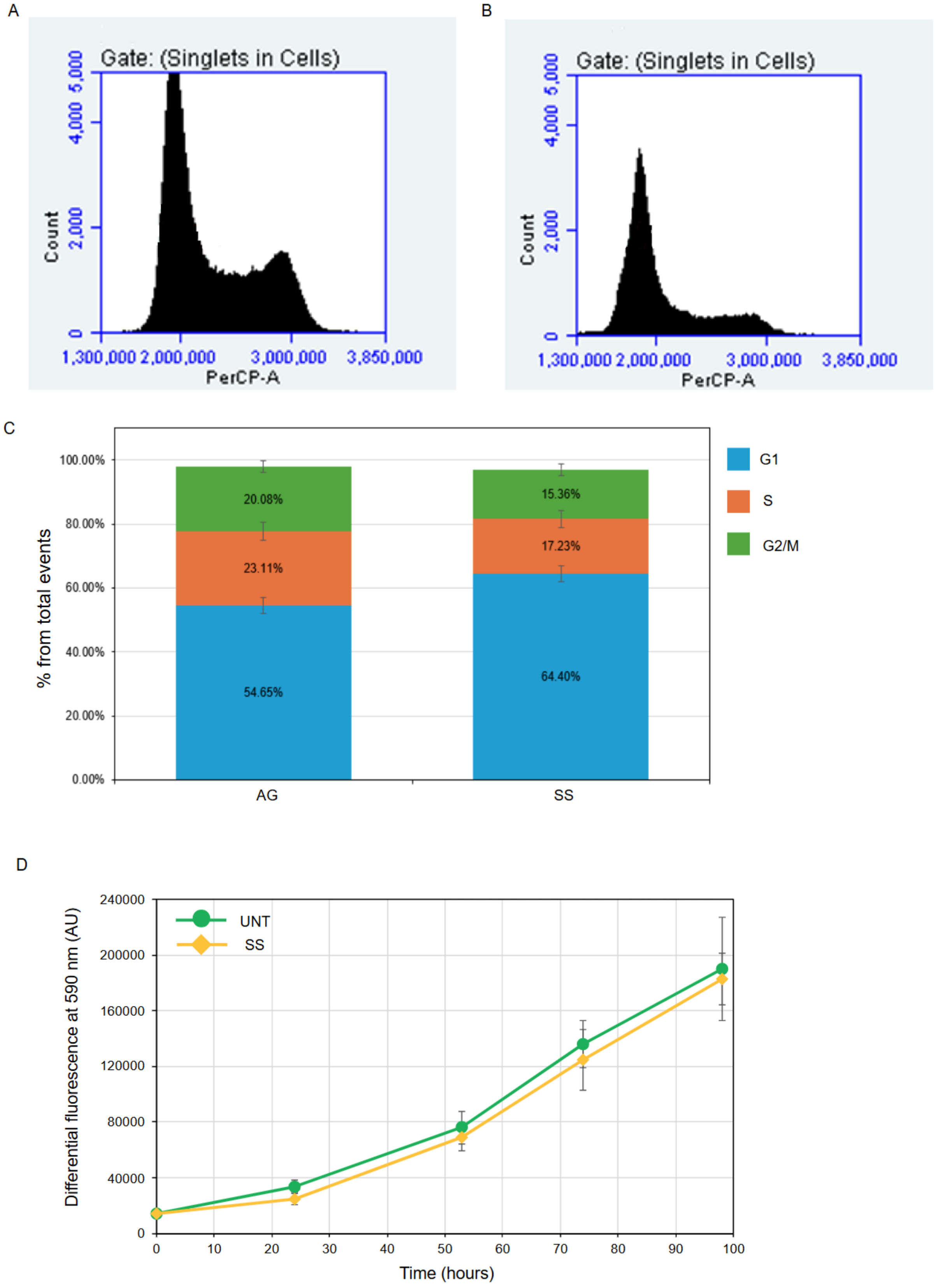


**Figure S1. Starvation induces G1 accumulation without compromising cell viability or proliferative recovery in U2OS cells.**
(A, B) Representative flow cytometry histograms of propidium iodide (PI)-stained U2OS cells gated on singlets. The x-axis shows PI fluorescence intensity (PerCP-A channel) and the y-axis

shows cell count. (A) Actively growing (AG) cells display a characteristic asynchronous cell cycle profile with distinct G1 and G2/M peaks. (B) Starved (SS) cells show a shift in cell cycle distribution with an increased proportion of cells in G1 phase relative to AG controls, consistent with starvation-induced G1 accumulation.

(C) Quantification of cell cycle phase distribution from PI-FACS analysis. Stacked bar chart shows the percentage of total events in G1 (blue), S (orange), and G2/M (green) phases for AG and SS conditions. AG cells exhibit 54.65% G1, 23.11% S, and 20.08% G2/M. Starved cells show an increased G1 fraction (64.40%) with concomitant reductions in S phase (17.23%) and G2/M (15.36%), indicating partial G1 arrest upon nutrient deprivation. Error bars represent standard deviation (n=3).

(D) Proliferative recovery assessed by PrestoBlue metabolic activity assay. Equal numbers of untreated (UNT; green squares) and previously starved (SS; orange diamonds) cells were replated in complete medium, and metabolic activity was measured by differential fluorescence at 590 nm (arbitrary units, AU) over a time course from 0 to 96 hours. Both conditions show progressive increases in fluorescence signal, indicating active cell proliferation following replating. Starved cells display comparable metabolic activity relative to untreated controls across all time points, demonstrating that proliferative capacity is fully retained following 18 h of starvation. Error bars represent standard deviation from biological replicates (n=3).

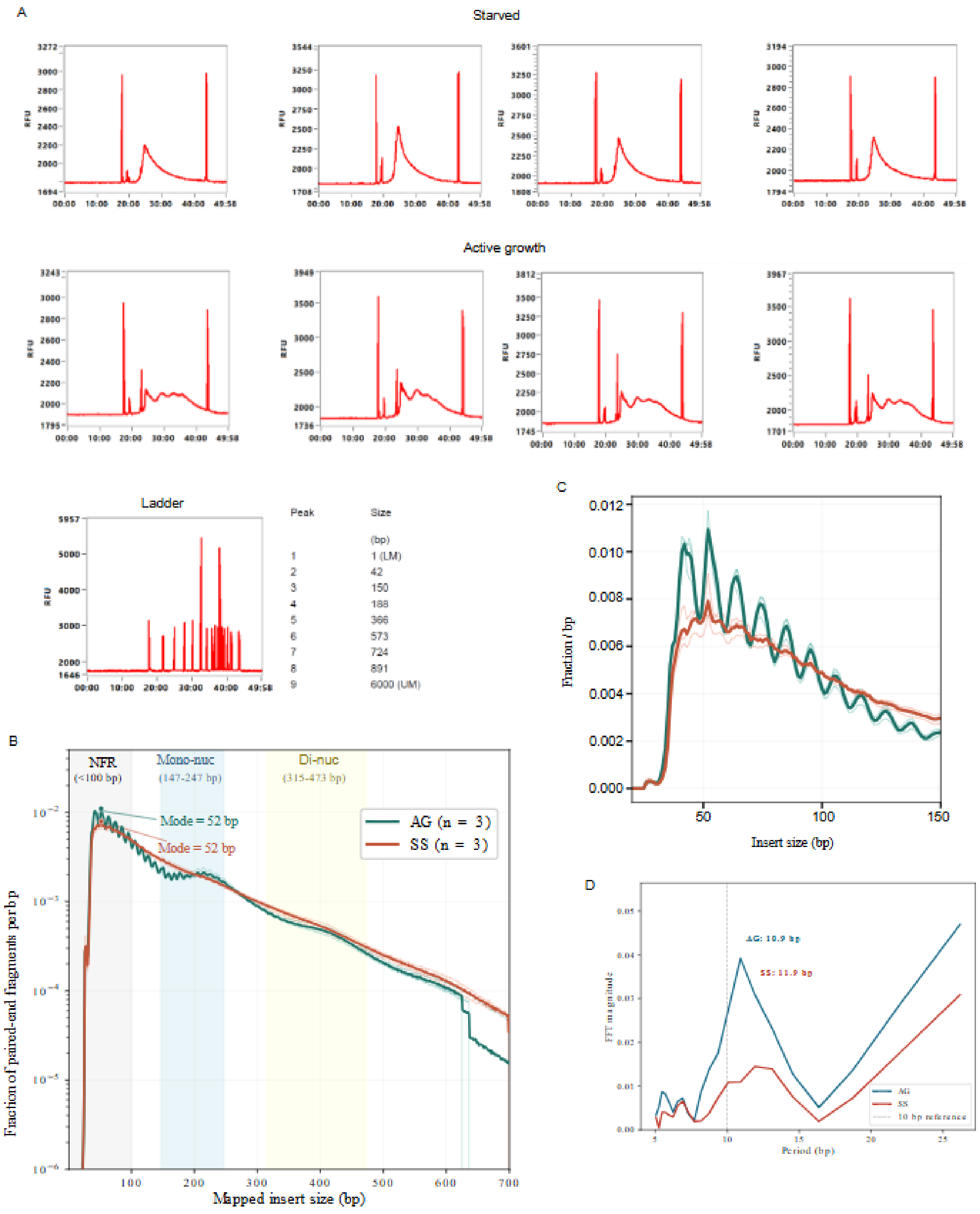


**Figure S2. Fragment Analyzer profiles and mapped ATAC-seq insert-size distributions in actively growing and starved U2OS cells.**

(A) Representative pre-sequencing electropherogram traces generated using the Agilent 5200 Fragment Analyzer for ATAC-seq libraries prepared from actively growing (AG) and starved (SS) U2OS cells. AG samples display sharply resolved mono- and dinucleosomal periodicity, whereas SS samples show broader, less discrete fragment distributions. Both conditions retain characteristic ATAC-seq fragment architecture, including a prominent nucleosome-free region (NFR) peak. Fragment Analyzer traces measure full library molecules, including approximately 130 bp of sequencing-adapter sequence; mapped genomic inserts of approximately 50 bp, 200 bp and 400 bp therefore correspond to library-molecule peaks at approximately 180 bp, 330 bp and 530 bp, respectively. The lower-right panel shows the Agilent DNF-474 size standard used for calibration and is not an ATAC-seq sample trace.

(B) Mapped paired-end insert-size distributions for AG and SS ATAC-seq libraries, plotted from 1 to 700 bp. Thin lines show individual biological replicates and bold lines show condition means. Shaded regions indicate the fragment-class windows used in Figure 1A: NFR (≤100 bp), mononucleosome (147–247 bp) and dinucleosome (315–473 bp). The modal insert size was 52 bp in both conditions.

(C) Linear-scale enlargement of the NFR sub-distribution (20–150 bp) showing the characteristic approximately 10 bp Tn5-associated rotational periodicity in both AG and SS libraries.

(D) Fast Fourier transform (FFT) periodogram of the NFR sub-distribution shown in panel C. The dominant spectral component remained close to the canonical approximately 10 bp periodicity in both conditions, with no significant difference in dominant period between AG and SS libraries ($p = 0.212$, Welch's t-test). FFT magnitude was significantly reduced in SS relative to AG, consistent with lower overall ATAC-seq signal in starved cells, but the periodicity itself was retained. This pattern is inconsistent with indiscriminate over-tagmentation, which would be expected to degrade or abolish this structural signature.

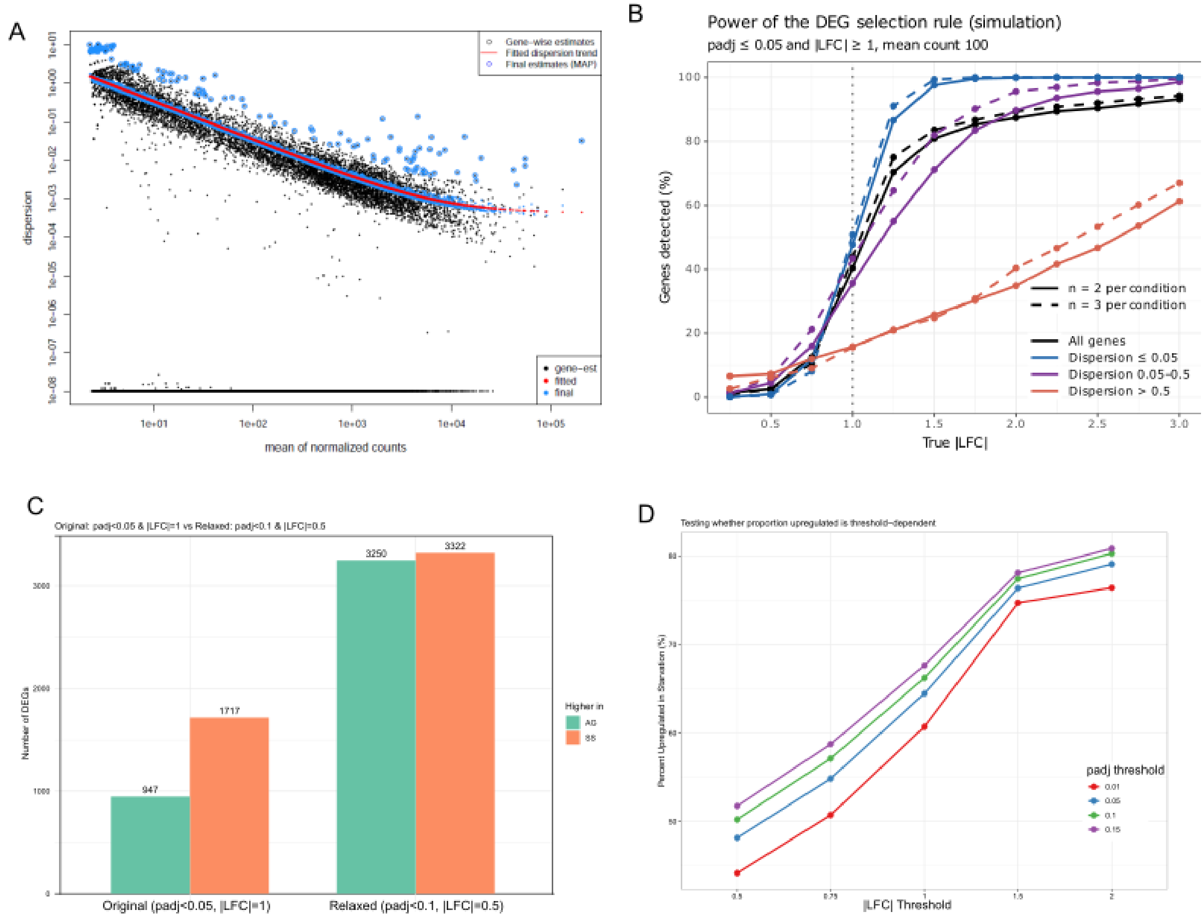


**Figure S3. Statistical characterization of RNA-seq differential expression analysis with n=2 replicates.**

(A) DESeq2 dispersion estimates: gene-wise estimates (black points), fitted mean-dispersion trend (red line) and final maximum a posteriori (MAP) estimates (blue points). With two replicates, DESeq2 relies on empirical Bayes shrinkage towards the fitted trend. Median dispersion = 0.023.

(B) Power of the DEG selection rule (padj ≤ 0.05 and |LFC| ≥ 1) estimated by simulation. Negative binomial counts were simulated for 30,000 genes (mean normalised count 100, dispersions matched to the observed distribution, true changes in 25% of genes) and analysed with DESeq2 as in the main analysis. Lines show the percentage of genes with a true change of the given |LFC| that met the rule, for all genes (black) and by dispersion class (blue for dispersion ≤ 0.05, purple for 0.05 to 0.5 and red for > 0.5), with two (solid) or three (dashed) replicates per condition. The dotted line marks |LFC| = 1. Because the rule requires an estimated |LFC| ≥ 1, power at a true |LFC| of 1 is limited to about 50% even for genes with low dispersion.

(C) Number of DEGs higher in AG (green) or SS (orange) with the original thresholds (padj < 0.05, |LFC| ≥ 1) and with relaxed thresholds (padj < 0.1, |LFC| ≥ 0.5). The number of DEGs increased from 2,664 to 6,572 and the proportion of genes increased in SS decreased from 64.5% to 50.5%.

(D) Percentage of DEGs increased in SS across |LFC| thresholds (0.5 to 2.0) and padj thresholds (0.01 to 0.15). The proportion of increased genes rises with the fold-change threshold at every padj threshold, showing that the excess of increased genes is confined to larger changes.

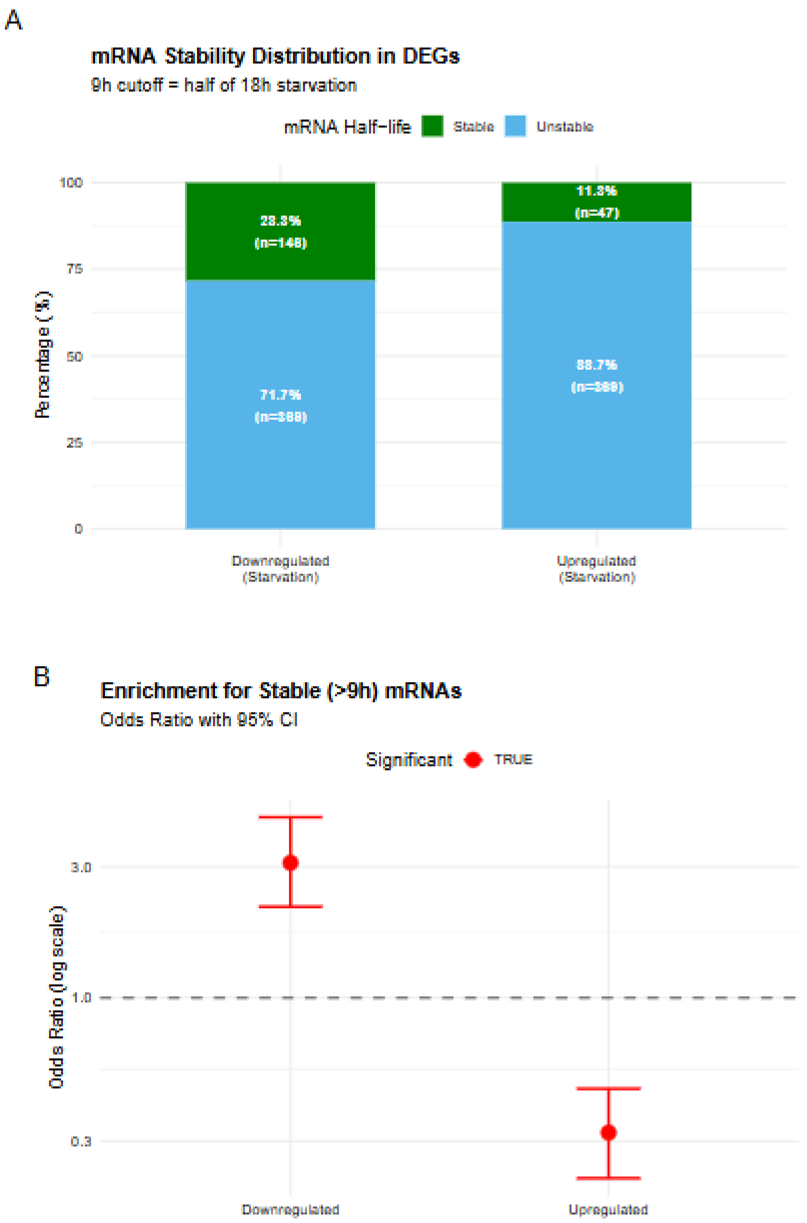


**Figure S4. Enrichment of short-lived mRNAs among upregulated genes, using published HeLa half-lives.**

(A) Stacked bar chart showing mRNA stability distribution in differentially expressed genes. Upregulated genes (n=416) and downregulated genes (n=515) are stratified by mRNA half-life using a 9-hour threshold (half the 18-hour starvation duration). Green bars represent stable mRNAs (half-life >9h); blue bars represent unstable mRNAs (half-life ≤9h). Percentages and gene counts are displayed within each segment. Upregulated genes show strong enrichment for unstable mRNAs (88.7%, n=369), whereas downregulated genes are enriched for stable mRNAs (28.3%, n=146). Under first-order decay kinetics, mRNAs with half-lives ≤9h would be reduced to ≤25% of initial abundance after 18 hours without new synthesis, consistent with a predominantly transcriptional contribution to their elevated abundance, although selective stabilisation of a subset of transcripts cannot be fully excluded by half-life data alone.

(B) Odds ratio plot with 95% confidence intervals for enrichment of stable (>9h) mRNAs. Each point represents the odds ratio for a differential expression category, with error bars indicating 95% confidence intervals. The horizontal dashed line at OR=1 indicates no enrichment. Upregulated genes show significant depletion of stable mRNAs (OR=0.322, 95%

CI [0.225–0.461], $p=8.5\times10^{-11}$), corresponding to a 3.1-fold enrichment for unstable mRNAs. Downregulated genes show the reciprocal pattern with significant enrichment for stable mRNAs (OR=3.10, 95% CI [2.17–4.45], $p<10^{-10}$). Red points indicate statistically significant enrichment ($p<0.05$); grey points indicate non-significant results. Half-life data from [1]. Statistical analysis by Fisher's exact test.

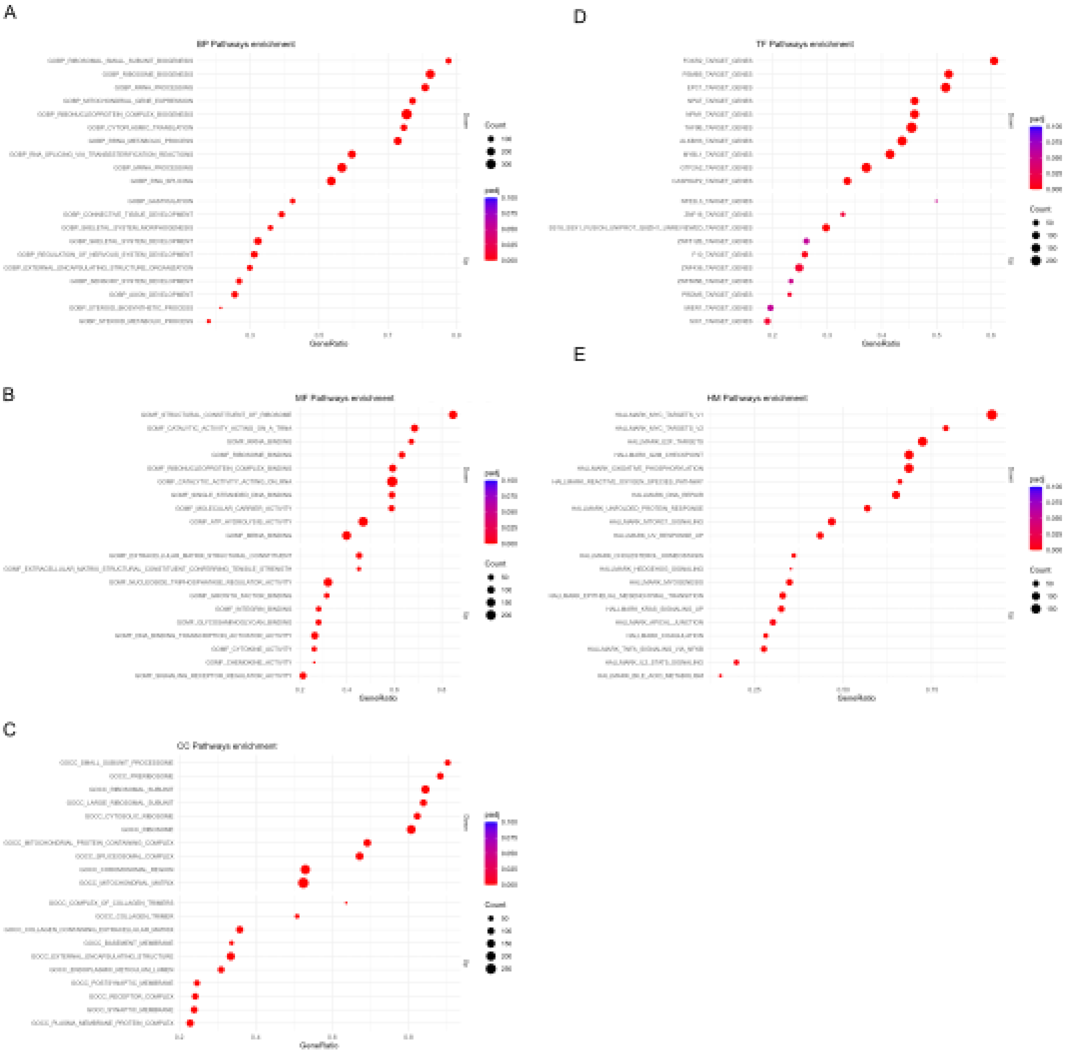


**Figure S5. Individual pathway enrichment results.**
(A) Dot plot of significantly enriched GO:BP pathways. (B) Dot plot of significantly enriched GO:MF pathways. (C) Dot plot of significantly enriched GO:CC pathways. (D) Dot plot of significantly enriched TF targets. (E**)** Dot plot of significantly enriched Hallmark pathways. Dot size represents leading-edge gene count; colour indicates padj value.

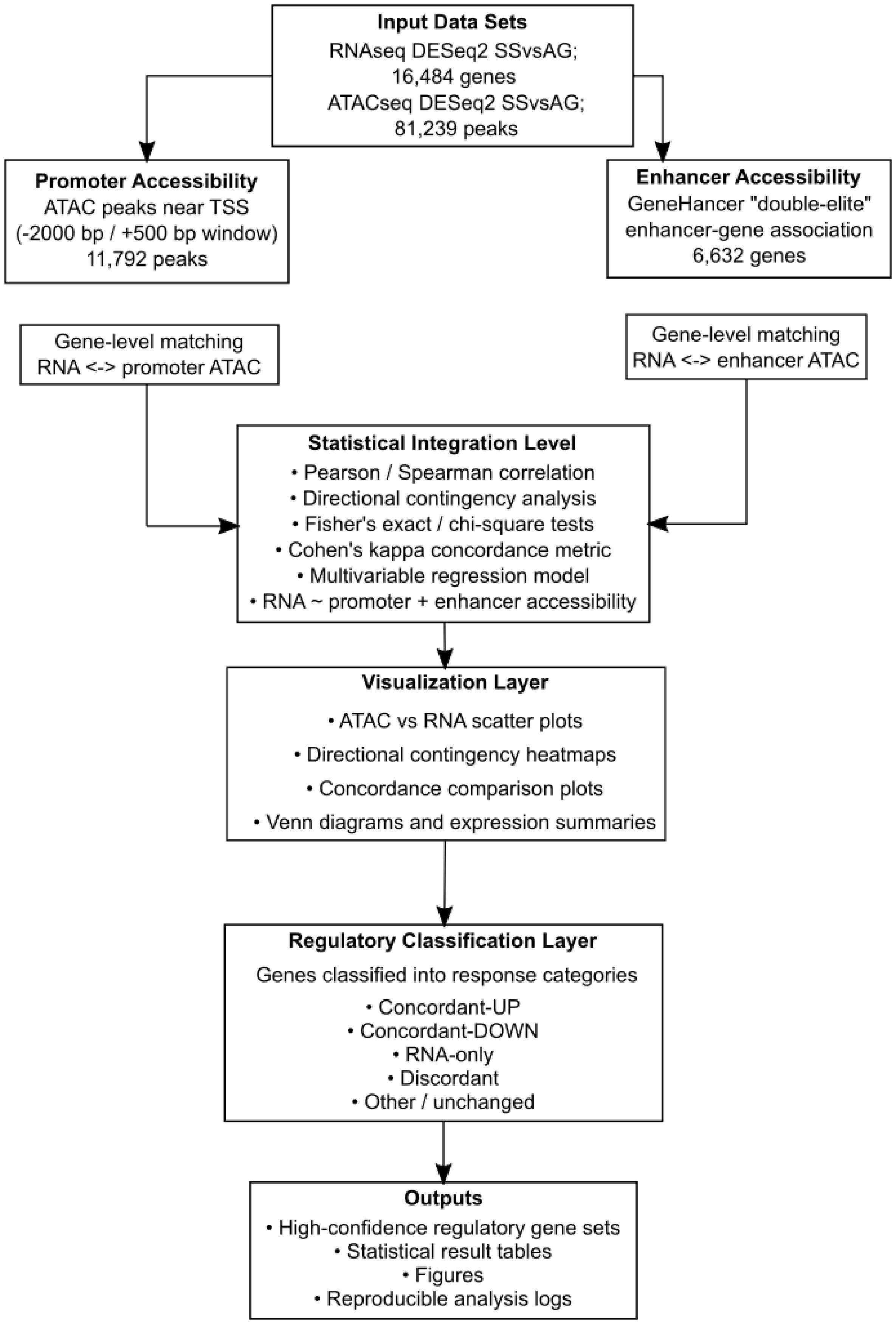
Input Data Sets
RNAseq DESeq2 SSvsAG;
16,484 genes
ATACseq DESeq2 SSvsAG;
81,239 peaks
Promoter Accessibility
ATAC peaks near TSS
(-2000 bp / +500 bp window)
11,792 peaks
Enhancer Accessibility
GeneHancer "double-elite"
enhancer-gene association
6,632 genes
Gene-level matching
RNA <-> promoter ATAC
Gene-level matching
RNA <-> enhancer ATAC
Statistical Integration Level
• Pearson / Spearman correlation
• Directional contingency analysis
• Fisher's exact / chi-square tests
• Cohen's kappa concordance metric
• Multivariable regression model
• RNA ~ promoter + enhancer accessibility
Visualization Layer
• ATAC vs RNA scatter plots
• Directional contingency heatmaps
• Concordance comparison plots
• Venn diagrams and expression summaries
Regulatory Classification Layer
Genes classified into response categories
• Concordant-UP
• Concordant-DOWN
• RNA-only
• Discordant
• Other / unchanged
Outputs
• High-confidence regulatory gene sets
• Statistical result tables
• Figures
• Reproducible analysis logs

**Figure S6. Architecture of the ATAC–RNA integration pipeline.** Differential expression results from RNA-seq and differential chromatin accessibility results from ATAC-seq were integrated using a modular R-based workflow. ATAC peaks were first annotated to genes using promoter proximity or high-confidence enhancer–gene associations from the GeneHancer database. RNA and ATAC features were matched at the gene level separately for promoter and enhancer regulatory layers. Statistical integration analyses quantified chromatin–transcription relationships using correlation analysis, contingency testing, concordance metrics, and multivariable regression models. Visualization modules generated exploratory and publication-quality plots summarizing regulatory patterns. Genes were subsequently classified into regulatory response categories including concordant regulation, ATAC-only, RNA-only, and discordant responses, and high-confidence regulatory events were identified for downstream analysis.

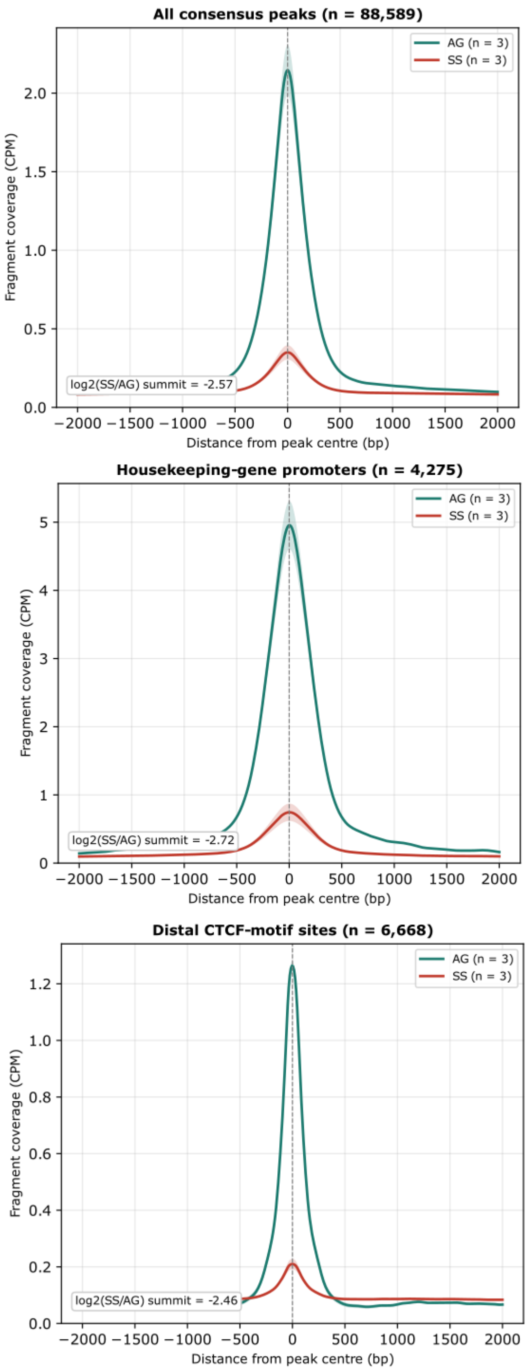


**Figure S7. Aggregate ATAC-seq signal metaplots centred on consensus peak summits and on constitutively accessible sites.**
Mean ATAC-seq fragment coverage, scaled to counts per million mapped fragments (CPM), ± SEM across biological replicates (n = 3 per condition), over a ±2 kb window centred on peak summits for all consensus peaks (left; n = 88,589), housekeeping-gene promoter peaks (middle; n = 4,275) and distal CTCF-motif sites (right; n = 6,668) in actively growing (AG, green) and starved (SS, red) cells. log2(SS/AG) at the summit: -2.57, -2.72 and -2.46, respectively. Replicate-level quantification of summit, flank and random-background signal is provided in Table S12 and Supplementary Figure S10.

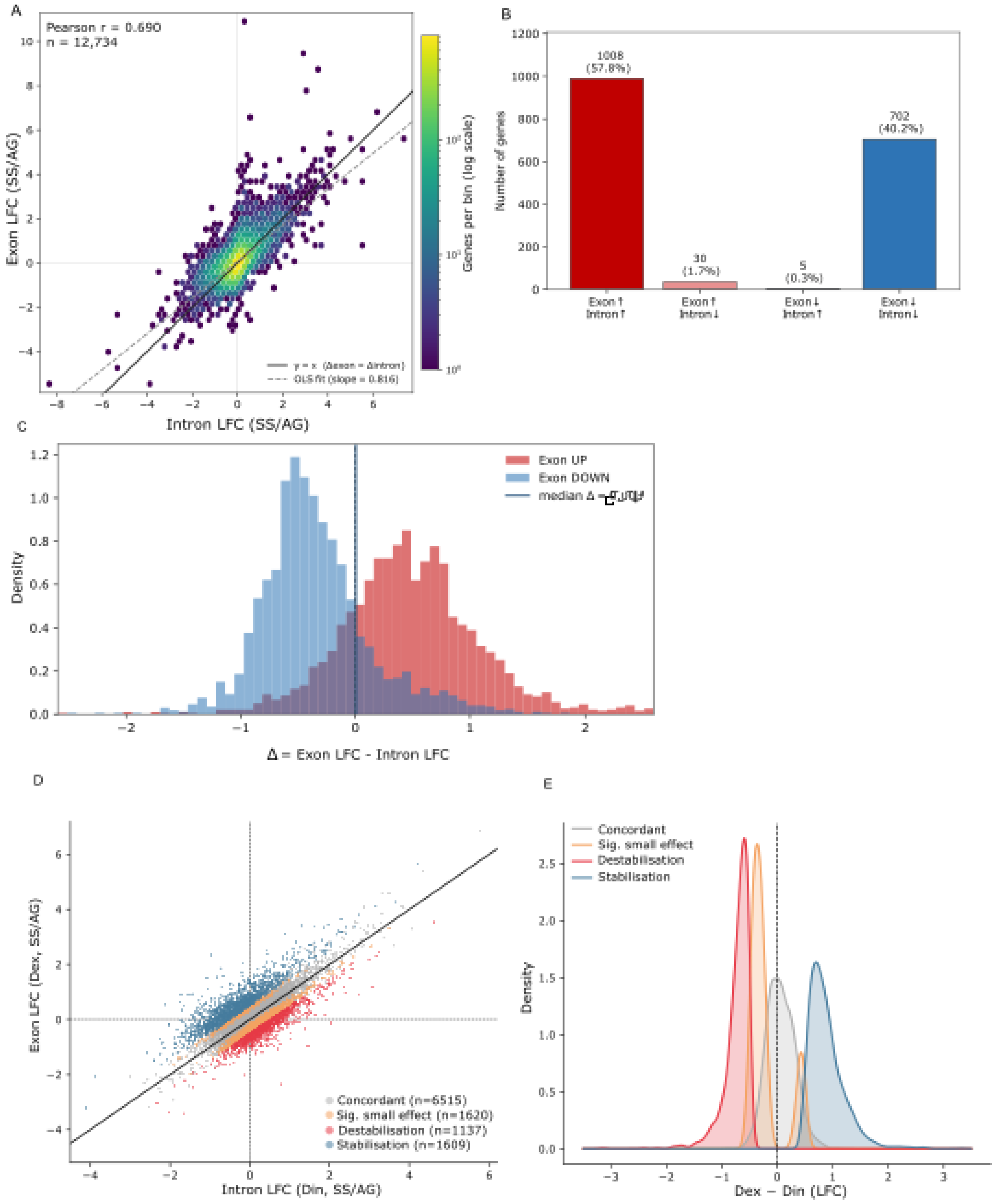


**Figure S8. Exon–intron concordance and EISA analysis of starvation-induced RNA-abundance changes.**

(A) Hex-binned density scatter of apeglm-shrunken exonic versus intronic log2 fold-changes (SS/AG) for 12,734 filtered protein-coding genes. The solid diagonal indicates y = x; the dashed line shows the ordinary least-squares regression fit. Exonic and intronic responses were positively correlated genome-wide (Pearson $r = 0.690$; Spearman $\rho = 0.625$; both $p < 1 \times 10^{-300}$), and 71.9% of genes showed the same sign of change.

(B) Directional classification of genes significant independently in both compartments (padj $\leq 0.05$, $|LFC| \geq 0.5$; $n = 1{,}745$). Most genes changed in the same direction in exons and introns: 1,008 were concordantly upregulated and 702 were concordantly downregulated, whereas only 35 genes were discordant. Overall same-direction concordance was 98.0% (1,710/1,745), exceeding the label-shuffled null expectation ($p < 1 \times 10^{-4}$).

(C) Density distributions of Δ = exon LFC - intron LFC for genes with significantly increased or decreased exonic signal. The genome-wide median Δ was +0.003, indicating no systematic excess of exonic over intronic response.

(D) Formal EISA scatter plot of exonic (Dex) versus intronic (Din) LFCs for 10,881 genes tested using eisaR. Points are coloured by EISA interaction class: stabilisation candidates (Dex - Din >= 0.5, FDR < 0.05, n = 1,609), destabilisation candidates (Dex - Din <= -0.5, FDR < 0.05, n = 1,137), significant small-effect interactions (n = 1,620) and concordant/background genes (n = 6,515). Sign convention follows Gaidatzis et al. {Gaidatzis, 2015 #5563}: a positive interaction term indicates that mature transcript abundance changed more than production and is interpreted as stabilisation; a negative term indicates destabilisation. Dex and Din were strongly correlated (Pearson r = 0.720). At |Dex - Din| >= 1.0 and FDR < 0.05, 542 genes (5.0%) showed evidence of post-transcriptional regulation, comprising 437 stabilisation and 105 destabilisation candidates.

(E) Density distributions of the EISA interaction estimate Dex - Din by interaction class. The concordant/background class was centred near zero; stabilisation candidates were shifted toward positive values and destabilisation candidates toward negative values, consistent with the sign convention in (D). Together, these analyses support a predominantly transcriptional component to the starvation RNA response. Significant interactions were detected in a minority of genes, mostly in the positive (stabilisation) direction; because most of these genes showed reduced or unchanged intronic signal, a positive interaction is also compatible with persistence of the mature transcript pool during reduced RNA production and does not by itself demonstrate active stabilisation.

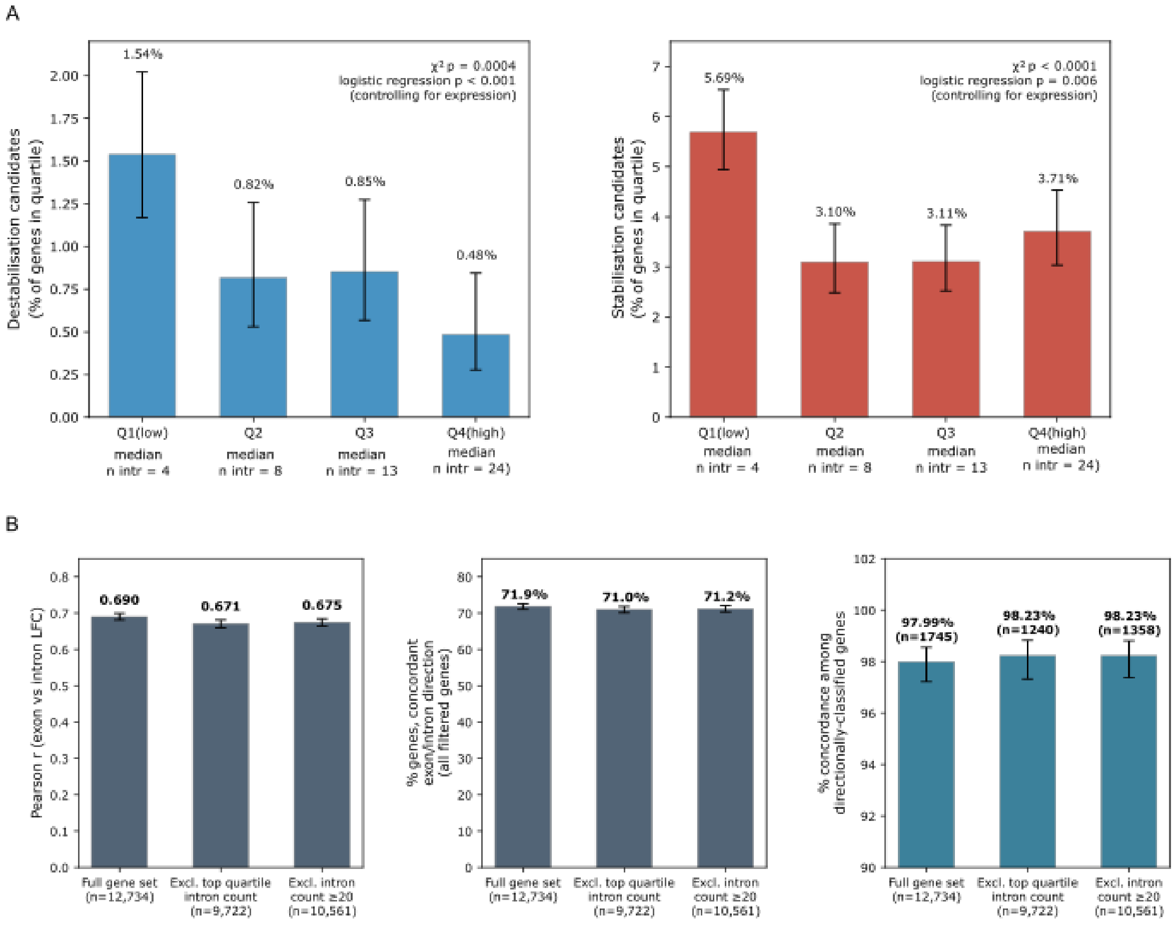

**Figure S9. Intron-count sensitivity analysis of EISA interaction classes and exon-intron concordance.**

(A) Genes passing EISA filtering (n = 10,881) were stratified into quartiles by intron count, and the percentage of genes meeting the stringent post-transcriptional threshold ($|Dex - Din| >= 1.0$, $FDR < 0.05$) was calculated separately for the destabilisation and stabilisation classes in each quartile. Because intron retention inflates intronic counts and would drive the interaction term negative, an intron-retention artefact would be expected to preferentially inflate the destabilisation class among genes with more introns. Contrary to this prediction, destabilisation candidates decreased with intron count (1.5%, 0.8%, 0.9% and 0.5% across quartiles; chi-squared = 17.5, $p = 0.0004$), and the association remained significant after adjustment for baseline expression level (logistic regression, $p < 0.001$). Stabilisation candidates, a class not expected to be inflated by intron retention, showed a broadly similar quartile pattern (5.7%, 3.1%, 3.1% and 3.7%; chi-squared $p < 0.0001$; logistic regression $p = 0.006$). Error bars show 95% Wilson score confidence intervals.

(B) Core exon–intron concordance metrics were recomputed after excluding genes with high intron counts: the top quartile by intron count (n = 3,012 excluded) or all genes with ≥20 introns (n = 2,173 excluded). Pearson correlation, same-sign concordance across all filtered genes and concordance among directionally classified genes remained essentially unchanged, indicating that the genome-wide exon–intron concordance pattern is not driven by genes most susceptible to intron-retention artefacts. Error bars: 95% CI (Fisher z-transformation for Pearson r; Wilson score interval for proportions). Gene sets are nested by construction (excluded sets are subsets of the full set), so a between-bar significance test is not reported.

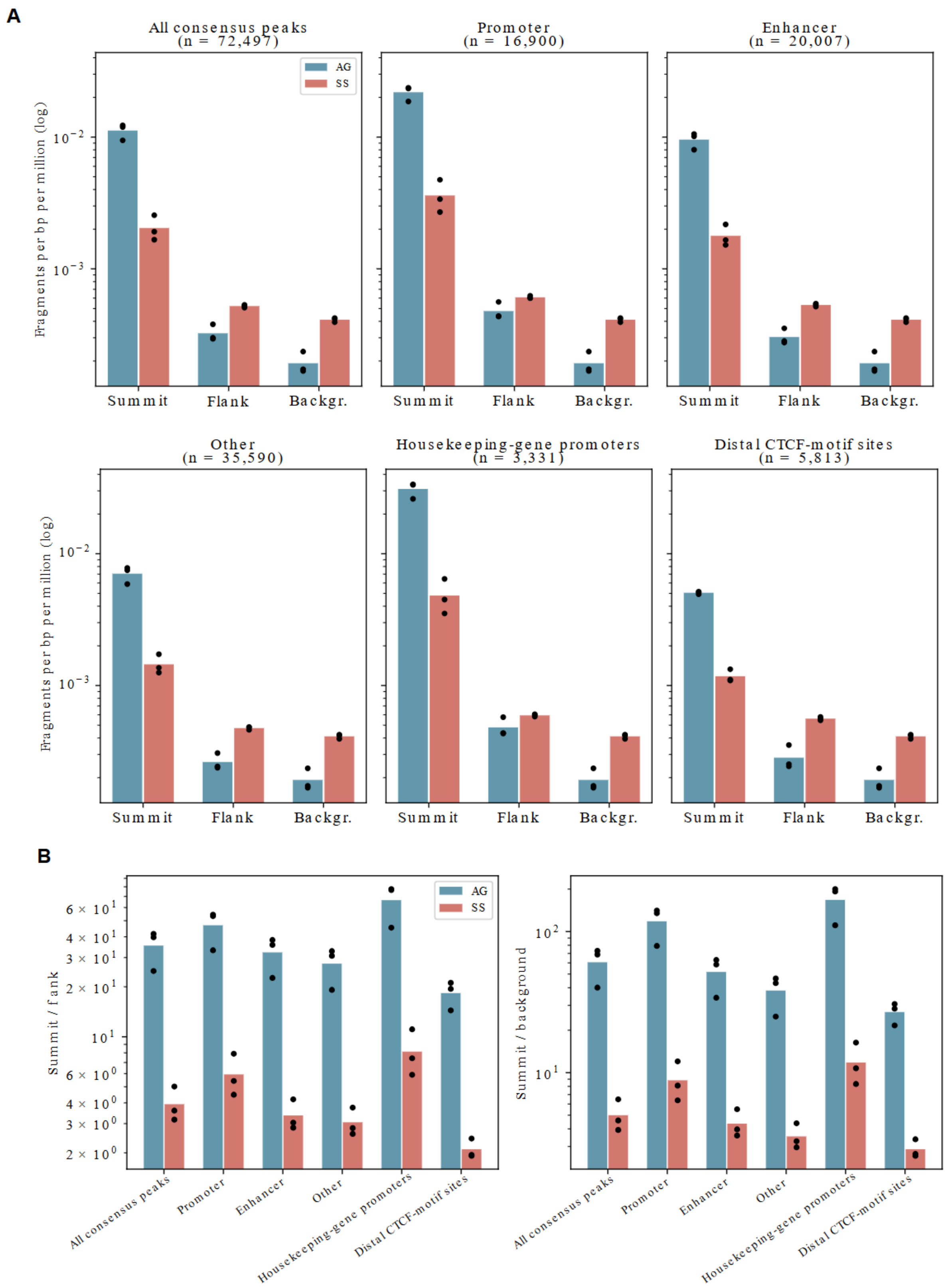


**Figure S10. Replicate-level quantification of summit, flank and background signal.**

(A) Mean signal (CPM per bp, log scale) at peak summits (±250 bp), local flanks (1-2 kb) and matched random-background regions in actively growing (AG) and starved (SS) cells, for all consensus peaks with clean flanks (n = 72,497) and for promoter, enhancer and other peaks, housekeeping-gene promoters and distal CTCF-motif sites (n shown above each panel). Bars, condition means; dots, individual biological replicates (n = 3 per condition). The random background is the same for all peak sets.
(B) Summit/flank (left) and summit/random-background (right) ratios for the same peak sets. Statistics are given in Table S12 and Supplementary Data Section 1.6.

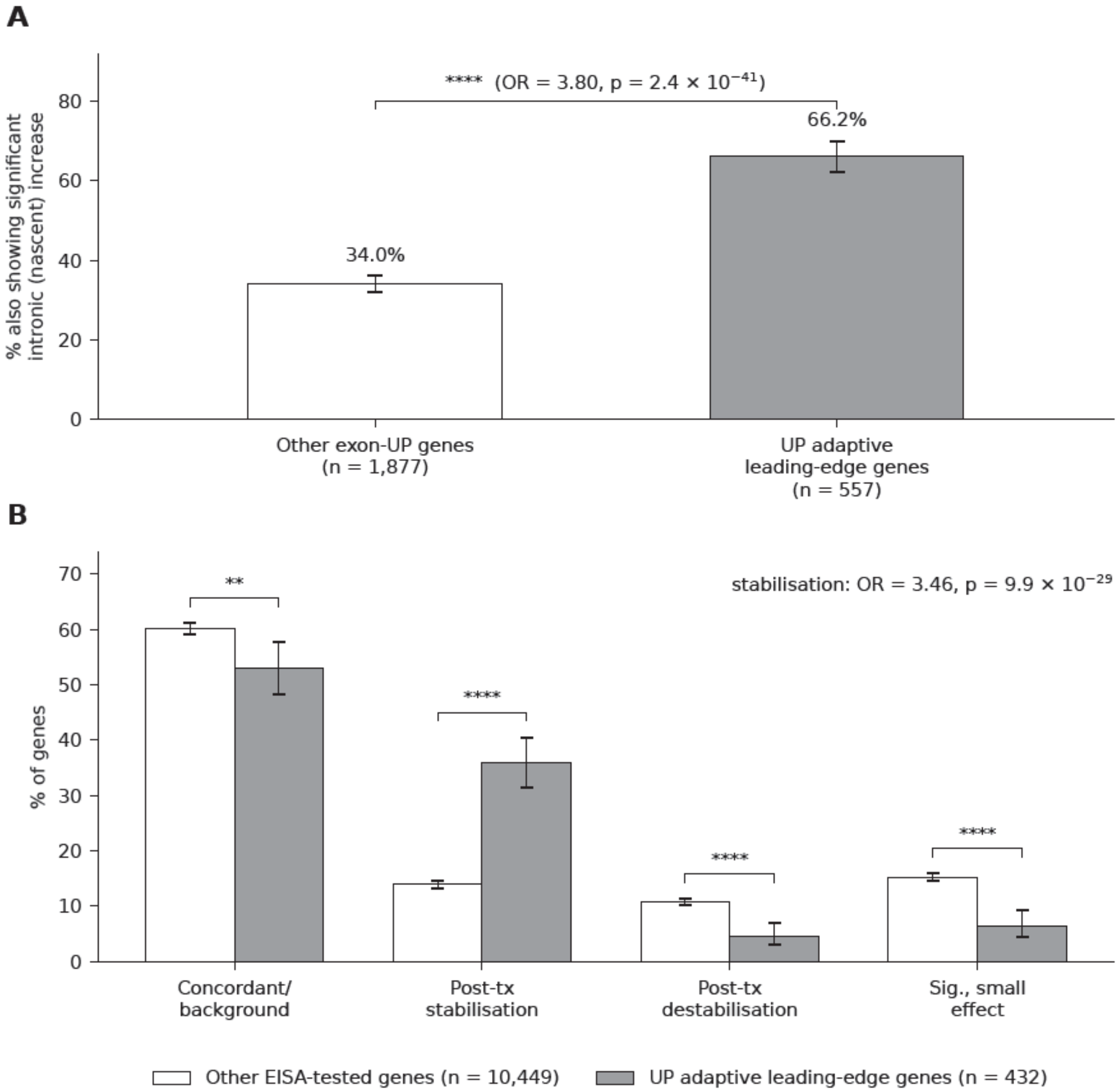


**Figure S11. Exon-intron concordance and EISA analysis of upregulated leading-edge genes in starvation-adaptive programmes.**

(A) Among exon-upregulated genes, adaptive-programme leading-edge genes were more likely than the remaining exon-upregulated genes to show a significant concordant increase in intronic signal (66.2% vs 34.0%, OR = 3.80, Fisher's exact $p = 2.4 \times 10^{-41}$), supporting increased RNA production. The two groups are disjoint.
(B) EISA interaction-class distributions for upregulated adaptive leading-edge genes (n = 432) and all other EISA-tested genes (n = 10,449). Leading-edge genes were enriched for the positive-interaction (stabilisation) class (35.9% vs 13.9%, OR = 3.46, Fisher's exact $p = 9.9 \times 10^{-29}$) and depleted of the destabilisation (4.6% vs 10.7%), small-effect (6.5% vs 15.2%) and concordant/background (53.0% vs 60.2%) classes. The concordant/background class (FDR ≥ 0.05) also contains genes with no change in exonic or intronic signal, so its depletion is expected for strongly changed genes. Together with (A), this indicates that adaptive genes are transcriptionally induced and raises the possibility of an additional post-transcriptional layer. However, none of the three stabilisation candidates tested by nascent RNA measurement (IGFBP7, CD9 and KLHL24) showed a significant change in the nascent-to-total ratio (**Figure 9, Table S17B**). Error bars show 95% Wilson confidence intervals. Brackets indicate Fisher's exact test significance (**$p < 0.01$, ****$p < 0.0001$).

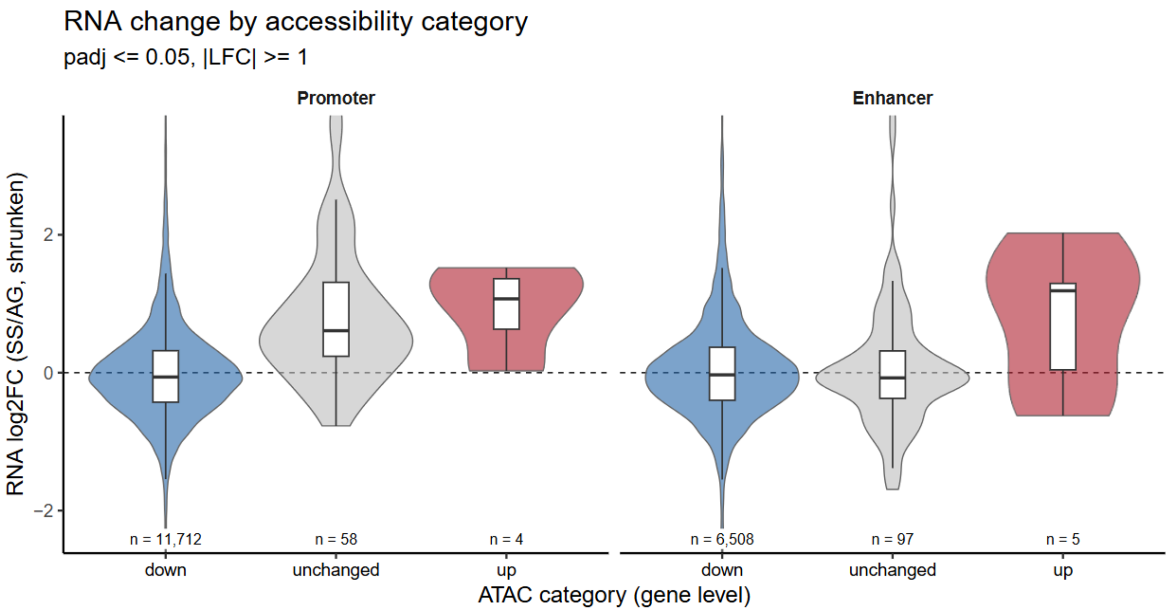


**Figure S12. mRNA changes by direction of accessibility change.**
RNA LFC (apeglm-shrunken) of genes grouped by gene-level promoter (left) and enhancer (right) accessibility category (reduced, unchanged, increased; padj ≤ 0.05, |LFC| ≥ 1). Violin plots with box plots showing median and interquartile range; numbers of genes are shown below each group. Genes with unchanged promoter accessibility (n = 58) had a median RNA LFC of 0.61, compared with -0.06 for genes with reduced promoter accessibility (n = 11,712).

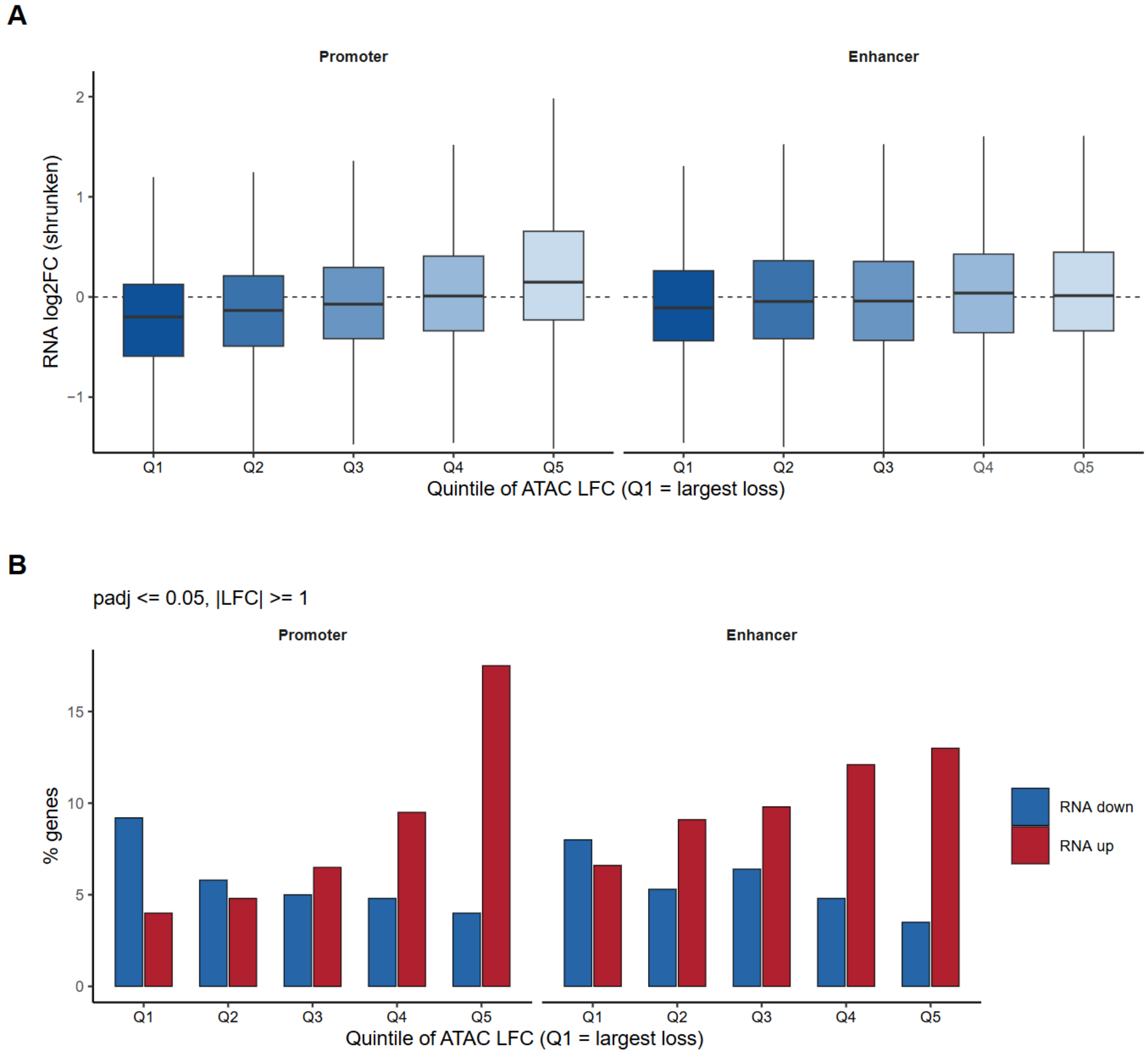


**Figure S13. mRNA changes across quintiles of promoter and enhancer accessibility change.**

(A) RNA LFC (apeglm-shrunken) across quintiles of gene-level promoter (left; n = 11,792) and enhancer (right; n = 6,632) accessibility change (Q1, largest loss). (B) Percentage of genes with decreased (blue) or increased (red) mRNA in each quintile (padj $\leq 0.05$, $|LFC| \geq 1$). The gradient is weaker for enhancers (odds ratio for increased mRNA, Q5 vs Q1: promoter 5.05, enhancer 2.11).

1. Tani H, Mizutani R, Salam KA, Tano K, Ijiri K, Wakamatsu A, Isogai T, Suzuki Y, Akimitsu N: **Genome-wide determination of RNA stability reveals hundreds of short-lived noncoding transcripts in mammals**. *Genome research* 2012, **22**(5):947-956.
2. Gaidatzis D, Burger L, Florescu M, Stadler MB: **Analysis of intronic and exonic reads in RNA-seq data characterizes transcriptional and post-transcriptional regulation**. *Nature biotechnology* 2015, **33**(7):722-729.
