## Supplementary tables and data for "Genome-wide loss of chromatin accessibility coexists with selectively elevated RNA abundance in nutrient-starved U2OS cells"

**Supplementary Data:**

**Tables S1B, S5, S6, S7, S13, S14 and S15 are in files**

**Table S1A. Library quality metrics for starved (SS) and growing (AG) U2OS cells.**

| Metric | AG | SS | p |
|---|---|---|---|
| *Before filtering (all aligned read pairs, including mitochondrial and duplicate reads)* | | | |
| Aligned read pairs (millions) | 35.6 ± 3.1 | 35.8 ± 0.6 | 0.95 |
| Mapped reads (%) | 99.4 ± 0.1 | 99.1 ± 1.0 | 0.64 |
| Properly paired reads (%) | 98.1 ± 0.4 | 97.8 ± 0.9 | 0.66 |
| Mitochondrial reads (% of mapped reads) | 10.6 ± 1.6 | 9.7 ± 0.4 | 0.42 |
| Duplicate read pairs, Picard (%) | 27.9 ± 2.2 | 24.7 ± 1.0 | 0.11 |
| Non-redundant fraction (NRF), Picard-based | 0.72 ± 0.02 | 0.75 ± 0.01 | 0.11 |
| Estimated library size, Picard (millions of distinct molecules) | 52.8 ± 6.6 | 61.9 ± 3.4 | 0.13 |
| *After filtering (final alignments used for peak calling)* | | | |
| Read pairs (millions) | 22.1 ± 1.9 | 22.8 ± 0.6 | 0.56 |
| Aligned read pairs retained (%) | 61.9 ± 2.1 | 63.9 ± 1.0 | 0.24 |
| Properly paired reads (%) | 98.8 ± 0.4 | 98.9 ± 0.3 | 0.75 |
| MAPQ = 60 (unique) reads (%) | ~89 | ~89 | NA |
| FRiP (%) | 56.4 ± 7.7 | 7.2 ± 1.9 | 0.006 |
| TSS enrichment score | 10.3 ± 1.4 | 3.9 ± 0.8 | 0.005 |

Mean ± SD of three independent libraries per condition. p, Welch's two-sided t-test, unadjusted.
Before filtering: values from Picard MarkDuplicates and samtools on the alignments of each library before removal of mitochondrial, duplicate and low-quality reads (MultiQC report of the alignment workflow). Mitochondrial reads are the reads aligned to chrM as a percentage of all mapped reads. NRF is (read pairs − duplicate read pairs) / read pairs, calculated on all aligned pairs including mitochondrial reads. The preferred ENCODE value is above 0.9. The estimated library size is Picard's estimate of the number of distinct molecules in the library.
After filtering: values from the final alignments after removal of mitochondrial, duplicate and low-quality reads. The mitochondrial fraction and the duplicate rate are zero in these alignments by construction, and NRF and PBC1 are 1 by construction, so none of them is informative about library complexity and they are not listed. PBC1 and PBC2 were not calculated before filtering because they require read counts per genomic position.
FRiP, fraction of fragments in the IDR optimal peaks of the sample's own condition. TSS enrichment score, ratio of signal at transcription start sites to flanking signal.
Fragment-size, periodicity and per-sample metrics are in Table S1B (Excel file). NA, not applicable.

**Table S2. IDR reproducibility analysis for ATAC-seq peak calling.**

*A. True-replicate IDR (all pairwise comparisons)*

| Replicate pair | Active growth (AG) | Starvation (SS) |
|---|---|---|
| Rep1 vs Rep2 | 125,532 | 44,705 |
| Rep1 vs Rep3 | 120,213 | 36,868 |
| Rep2 vs Rep3 | 121,950 | 41,847 |
| Best pair (Nt) | 125,532 (Rep1 vs Rep2) | 44,705 (Rep1 vs Rep2) |

Peaks passing IDR ≤ 0.05.

*B. Self-consistency IDR (pseudoreplicates of each replicate)*

| Replicate | Active growth (AG) | Starvation (SS) |
|---|---|---|
| Rep1 | 96,052 | 23,748 |
| Rep2 | 94,651 | 37,098 |

| | | |
|---|---|---|
| Rep3 | 98,233 | 22,144 |
| Self-consistency ratio (max / min) | 1.04 | 1.68 |

Peaks passing IDR ≤ 0.05 between the two pseudoreplicates of each replicate (N1-N3).

*C. Peak set summary*

| **Peak set** | **Active growth (AG)** | **Starvation (SS)** | **Notes** |
|---|---|---|---|
| Best true-replicate pair, Nt | 125,532 | 44,705 | Panel A |
| Pooled pseudoreplicates, Np | 137,830 | 52,842 | Pooled pr1 vs pooled pr2 |
| Rescue ratio, max(Np, Nt) / min(Np, Nt) | 1.10 | 1.18 | ENCODE: ≤ 2 |
| Self-consistency ratio | 1.04 | 1.68 | ENCODE: ≤ 2 (panel B) |
| Reproducibility | pass | pass | See note b |
| Optimal peak set | 137,830 (pooled pseudoreplicates, Np) | 52,842 (pooled pseudoreplicates, Np) | Used to build the consensus set |
| Conservative peak set (Nt) | 125,532 | 44,705 | Not used downstream |
| Relaxed optimal set (IDR ≤ 0.10) | 146,543 | 57,941 | Not used downstream |
| Optimal peaks after width filter (50-2,000 bp) and blacklist | 137,673 | 52,762 | Annotation step (script 01) |
| Consensus peak set (AG and SS merged) | 88,589 | | Merge distance 50 bp; 81,239 peaks tested for differential accessibility |

[a.] n = 3 biological replicates per condition. Peaks were called with MACS2 on Tn5-shifted reads (+4 / -5 bp; -p 0.01 --nomodel --shift -75 --extsize 150 --keep-dup all --call-summits), keeping the 500,000 peaks with the lowest p values, on each replicate, the pooled replicates and self- and pooled pseudoreplicates generated by random read partitioning. IDR (--rank p.value --use-best-multisummit-IDR, pooled peaks as reference peak list) was run on all pairs; peaks with global IDR ≤ 0.05 were kept after removal of ENCODE blacklist regions (hg38 v2) and non-standard chromosomes.

[b.] ENCODE rules: the optimal set is the conservative set (Nt) if Nt > Np and the pooled-pseudoreplicate set otherwise; reproducibility passes when the rescue and self-consistency ratios are both ≤ 2, is borderline when one exceeds 2 and fails when both do. Np exceeded Nt in both conditions (rescue ratios 1.10 and 1.18), so the pooled-pseudoreplicate set was the optimal set; both conditions passed.

[c.] SS yielded 38% as many optimal peaks as AG. Its higher self-consistency ratio reflects fewer reproducible peaks in Rep1 and Rep3, the replicates with the lowest FRiP (6.7% and 5.5%, versus 9.3% in Rep2; Table S1A), consistent with low peak-centred signal rather than library failure.

**Table S3. Differential accessibility on the total-fragment and consensus-peak scales.**

| **Method** | **Role** | **Gain / loss (\|LFC\| ≥ 1)** | **Gain / loss (\|LFC\| ≥ 0.58)** | **Median LFC** | **r with library-size LFC** | **Median shift** |
|---|---|---|---|---|---|---|

| | | | | | | |
|---|---|---|---|---|---|---|
| Library size (total mapped fragments) | Primary | 698 / 77,075 | 740 / 78,075 | -2.34 | 1.000 | 0.00 |
| Fragments in consensus peaks | Relative representation within peak signal | 8,316 / 1,457 | 10,275 / 3,995 | 0.08 | 0.999 | 2.42 |

DESeq2 fits of the 81,239 tested peaks with each set of size factors. Gain / loss, peaks with padj ≤ 0.05 at the given |LFC| threshold. r, Pearson correlation of per-peak LFC with the library-size LFC. Median shift, median per-peak difference from the library-size LFC (log2 units). Library-size normalisation expresses each peak's signal per mapped fragment. Consensus-peak normalisation equalises the total peak-associated signal between conditions, so it measures each peak's share of the remaining peak signal. A positive LFC on this scale therefore indicates relative preservation within the peak pool and can occur while signal per mapped fragment decreases. The median shift of 2.42 matches the log2 ratio of the fraction of fragments in consensus peaks in AG and SS (56.5% and 10.6%, 2.41). Per-peak LFCs on the two scales therefore differ mainly by this constant. Correlations, rankings and comparisons between groups of peaks or genes are essentially unchanged, whereas the numbers of gained and lost peaks depend on the scale. TMM (edgeR) and median-of-ratios (DESeq2) normalisation on covered peaks gave results within 0.04 log2 units of consensus-peak normalisation (gain / loss 7,808 / 1,723 and 7,954 / 1,662 at |LFC| ≥ 1, r = 1.000 with library-size LFCs).

**Table S4. Summary of fGSEA pathway enrichment analysis (SS vs AG).**

| Gene Set Collection | Total | Up (NES≥0) | Down (NES≤0) | Top \|NES\| |
|---|---|---|---|---|
| GO:BP | 670 | 245 | 425 | 2.73 |
| GO:CC | 157 | 30 | 127 | 2.73 |
| GO:MF | 148 | 33 | 115 | 2.71 |
| Hallmark | 27 | 14 | 13 | 2.90 |
| TF Targets | 76 | 3 | 73 | 2.25 |
| Canonical Pathways | - | - | - | - |
| **Total** | **1078** | **325** | **753** | **-** |

FDR ≤ 0.01. Top |NES| indicates the highest absolute normalized enrichment score within each collection.

**Table S8. Within-condition correlations between promoter accessibility and gene expression.**

| Comparison | n genes | Pearson r (95% CI) | Spearman ρ (95% CI) | p (Pearson) | Type |
|---|---|---|---|---|---|
| Promoter ATAC vs RNA, AG | 11,792 | 0.284 (0.267-0.302) | 0.270 (0.252-0.287) | 1.3 × $10^{-217}$ | Within-condition |
| Promoter ATAC vs RNA, SS | 11,792 | 0.252 (0.235-0.270) | 0.245 (0.228-0.262) | 3.1 × $10^{-170}$ | Within-condition |

log2(normalised mean + 1) in each condition (ATAC, three replicates; RNA, two replicates); promoter accessibility = median over the gene's promoter peaks. 95% CI from 10,000 bootstrap resamples of genes.

**Table S9. Genome-wide agreement between exonic and intronic direction calls across the full filtered gene set.**

*(A) Contingency table of exonic vs intronic direction classification (n = 12,734 filtered protein-coding genes; padj ≤ 0.05, |LFC| ≥ 0.5 applied independently in each compartment).*

| Exon direction \ Intron direction | DOWN | UNCHANGED | UP | Row total |
|---|---|---|---|---|
| **DOWN** | 702 | 1,423 | 5 | **2,130** |
| **UNCHANGED** | 745 | 7,076 | 349 | **8,170** |
| **UP** | 30 | 1,396 | 1,008 | **2,434** |
| **Column total** | **1,477** | **9,895** | **1,362** | **12,734** |

*(B) Agreement statistics derived from the contingency table in (A).*

| Statistic | Observed agreement | Expected agreement (by chance) | Cohen's κ (unweighted) |
|---|---|---|---|
| Exon vs intron direction (DOWN/UNCHANGED/UP) | 69.0% | 53.8% | **0.328** |

Genes were classified independently in the exonic and intronic compartments as UP (padj ≤ 0.05 and log2FC ≥ 0.5), DOWN (padj ≤ 0.05 and log2FC ≤ -0.5), or UNCHANGED (all other genes), using the same thresholds applied throughout the exon–intron concordance analysis (Figure S8). The contingency table in (A) cross-tabulates these calls across all 12,734 filtered protein-coding genes. Cohen's κ in (B) is computed across all nine cells of this table (unweighted) and reflects agreement beyond that expected from the marginal distributions alone. The fair-to-moderate κ value (0.328) reflects the substantial proportion of genes classified as UNCHANGED in one or both compartments (10,989 of 12,734 genes (86.3%)), among which agreement is necessarily limited by the conservative significance and fold-change thresholds applied independently to each compartment; restricting to genes with a confident directional call in both compartments yields substantially higher concordance (98.0%; Figure S8B, main text).

**Table S10. Sensitivity of genome-wide exon-intron concordance to exclusion of high-intron-count genes.**

| Analysis | N | Pearson r (exon vs intron LFC) | % same sign (all filtered genes) | Directional-only concordance (%) | Directional-only N |
|---|---|---|---|---|---|
| **Full gene set** | 12,734 | 0.690 | 71.9% | 97.99% | 1,745 |
| **Excluding top quartile by intron count (intron count ≥ 14 removed)** | 9,722 | 0.671 | 71.0% | 98.23% | 1,240 |
| **Excluding genes with intron count ≥ 20** | 10,561 | 0.675 | 71.2% | 98.23% | 1,358 |

Core exon-intron concordance statistics (Figure S8A) were recomputed after excluding genes most plausibly vulnerable to intron-retention artefacts arising from reduced splicing capacity, using intron count as the exclusion criterion. "Directional-only concordance" denotes the percentage of genes with a statistically significant, large-magnitude directional call independently in both the exonic and intronic compartments (padj ≤ 0.05, |log2FC| ≥ 0.5) that changed in the same direction. All statistics were computed from the same exonic and intronic log2 fold-change values used throughout the exon-intron concordance analysis (Figure S8), on the same underlying gene set, differing only in which genes were included after exclusion. Corresponding bar charts are shown in Supplementary Figure S9B.

**Table S11. Cellular poly(A)+ RNA content in actively growing (AG) and starved (SS) U2OS cells.**

*A. Poly(A)+ RNA purified directly from cells*

| Replicate | Condition | Cells used (× $10^6$) | Poly(A)+ RNA (ng) | Poly(A)+ RNA per cell (pg) |
|---|---|---|---|---|
| 1 | AG | 19.0 | 13125 | 0.691 |
| 2 | AG | 19.0 | 13773 | 0.725 |
| 3 | AG | 19.0 | 15108 | 0.795 |
| Mean ± SD | AG | - | 14002 ± 1011 | 0.737 ± 0.053 |
| 1 | SS | 7.4 | 5621 | 0.760 |
| 2 | SS | 7.4 | 5144 | 0.695 |

| | | | | |
|---|---|---|---|---|
| 3 | SS | 7.4 | 5760 | 0.778 |
| Mean ± SD | SS | - | 5508 ± 323 | 0.744 ± 0.044 |

*B. Poly(A)+ RNA purified from total RNA*

| Replicate | Condition | Input total RNA (µg) | Poly(A)+ RNA (ng) | Yield (% of input) |
|---|---|---|---|---|
| 1 | AG | 200 | 15446 | 7.72 |
| 2 | AG | 200 | 17443 | 8.72 |
| 3 | AG | 200 | 15455 | 7.73 |
| Mean ± SD | AG | - | 16115 ± 1150 | 8.06 ± 0.58 |
| 1 | SS | 200 | 15543 | 7.77 |
| 2 | SS | 200 | 16549 | 8.27 |
| 3 | SS | 200 | 16534 | 8.27 |
| Mean ± SD | SS | - | 16209 ± 576 | 8.10 ± 0.29 |

*C. Summary statistics*

| Measurement | AG (mean ± SD) | SS (mean ± SD) | SS/AG ratio | 95% CI | t-test p | TOST p (± 25%) |
|---|---|---|---|---|---|---|
| Poly(A)+ RNA per cell (pg) | 0.737 ± 0.053 | 0.744 ± 0.044 | 1.01 | 0.87-1.17 | 0.86 | 0.008 |
| Poly(A)+ RNA yield (% of input total RNA) | 8.06 ± 0.58 | 8.10 ± 0.29 | 1.01 | 0.89-1.14 | 0.91 | 0.005 |

**Table S12. Peak summit, local flank and random-background signal in actively growing (AG) and starved (SS) U2OS cells.**

*A. Signal by region class (CPM per bp)*

| Peak set | Region class | AG mean ± SD | SS mean ± SD | Fold (SS/AG) | Welch p |
|---|---|---|---|---|---|
| All consensus peaks (n = 72,497) | Summit (±250 bp) | 0.01120 ± 0.00154 | 0.00204 ± 0.00046 | 0.18 | 0.006 |
| All consensus peaks | Local flank (1-2 kb) | 0.00032 ± 0.00005 | 0.00052 ± 0.00001 | 1.61 | 0.014 |
| All consensus peaks | Random background | 0.00019 ± 0.00004 | 0.00041 ± 0.00002 | 2.14 | 0.004 |
| Promoter (n = 16,900) | Summit (±250 bp) | 0.02188 ± 0.00282 | 0.00361 ± 0.00104 | 0.16 | 0.004 |
| Promoter | Local flank (1-2 kb) | 0.00048 ± 0.00007 | 0.00061 ± 0.00001 | 1.27 | 0.084 |
| Enhancer (n = 20,007) | Summit (±250 bp) | 0.00956 ± 0.00137 | 0.00178 ± 0.00035 | 0.19 | 0.007 |
| Enhancer | Local flank (1-2 kb) | 0.00030 ± 0.00004 | 0.00053 ± 0.00001 | 1.75 | 0.007 |
| Other (n = 35,590) | Summit (±250 bp) | 0.00706 ± 0.00103 | 0.00145 ± 0.00025 | 0.21 | 0.008 |
| Other | Local flank (1-2 kb) | 0.00026 ± 0.00004 | 0.00048 ± 0.00001 | 1.81 | 0.006 |

| | | | | | |
|---|---|---|---|---|---|
| Housekeeping-gene promoters (n = 3,331) | Summit (±250 bp) | 0.03101 ± 0.00425 | 0.00482 ± 0.00149 | 0.16 | 0.004 |
| Housekeeping-gene promoters | Local flank (1-2 kb) | 0.00048 ± 0.00008 | 0.00059 ± 0.00001 | 1.23 | 0.14 |
| Distal CTCF-motif sites (n = 5,813) | Summit (±250 bp) | 0.00506 ± 0.00012 | 0.00118 ± 0.00013 | 0.23 | 3.4 × $10^{-6}$ |
| Distal CTCF-motif sites | Local flank (1-2 kb) | 0.00028 ± 0.00006 | 0.00056 ± 0.00002 | 1.98 | 0.011 |

Random background is sampled genome-wide and is the same for all peak sets.

*B. Proportion of summit-proximal signal within the summit window*

| Peak set | AG mean ± SD (%) | SS mean ± SD (%) | Welch p |
|---|---|---|---|
| All consensus peaks | 89.4 ± 2.8 | 49.1 ± 5.9 | 0.002 |
| Promoter | 91.8 ± 2.2 | 58.9 ± 6.9 | 0.009 |
| Enhancer | 88.5 ± 3.1 | 45.3 ± 5.2 | 7.8 × $10^{-4}$ |
| Other | 86.8 ± 3.5 | 43.1 ± 4.7 | 3.3 × $10^{-4}$ |
| Housekeeping-gene promoters | 94.0 ± 1.8 | 66.1 ± 7.0 | 0.015 |
| Distal CTCF-motif sites | 81.8 ± 3.1 | 34.4 ± 3.1 | 4.6 × $10^{-5}$ |

*C. Derived ratio metrics*

| Peak set | Ratio | AG mean ± SD | SS mean ± SD | Fold (SS/AG) | Welch p |
|---|---|---|---|---|---|
| All consensus peaks | Summit/flank | 35.4 ± 9.2 | 3.9 ± 1.0 | 0.111 | 0.026 |
| All consensus peaks | Summit/random background | 60.6 ± 17.9 | 5.0 ± 1.3 | 0.082 | 0.032 |
| Promoter | Summit/flank | 46.9 ± 11.9 | 5.9 ± 1.8 | 0.126 | 0.025 |
| Promoter | Summit/random background | 118.2 ± 34.1 | 8.8 ± 2.9 | 0.075 | 0.030 |
| Enhancer | Summit/flank | 32.2 ± 8.4 | 3.4 ± 0.7 | 0.104 | 0.026 |
| Enhancer | Summit/random background | 51.8 ± 15.6 | 4.4 ± 1.0 | 0.084 | 0.034 |
| Other | Summit/flank | 27.6 ± 7.4 | 3.1 ± 0.6 | 0.111 | 0.028 |
| Other | Summit/random background | 38.2 ± 11.6 | 3.5 ± 0.7 | 0.093 | 0.035 |
| Housekeeping-gene promoters | Summit/flank | 66.5 ± 18.3 | 8.1 ± 2.7 | 0.122 | 0.029 |
| Housekeeping-gene promoters | Summit/random background | 167.7 ± 49.4 | 11.8 ± 4.1 | 0.070 | 0.031 |
| Distal CTCF-motif sites | Summit/flank | 18.3 ± 3.5 | 2.1 ± 0.3 | 0.115 | 0.015 |
| Distal CTCF-motif sites | Summit/random background | 26.9 ± 4.7 | 2.9 ± 0.4 | 0.107 | 0.012 |

Region definitions, normalisation and statistics are described in Supplementary Data Section 1.6. Signal scaled to counts per million mapped fragments and expressed per base pair; peaks whose 1-2 kb flanks overlap other consensus peaks or blacklisted regions excluded. Mean ± SD of replicate-level values (n = 3 per condition); two-tailed Welch's t-test.

**Table S16. qPCR assays used for total and nascent RNA quantification.**

| Target | Forward primer (5′–3′) | Reverse primer (5′–3′) | Type[a] | Melting peak Tm, °C[b] |
|---|---|---|---|---|
| FN1 | TCCCCTGGGGTCACCTATTAC | GTATCCTGTTATCTGGGCCCG | S | 83.1 ± 0.22 |
| MMP2 | GAAGGCTGTGTTCTTTGCAGG | TTTTGCTCCAGTTAAAGGCGG | S | 86.0 ± 0.02 |
| IGFBP7 | GTGGCCCAGAAAAGCATGAAG | CTGAAGCCTGTCCTTGGGAAT | U | 80.7 ± 0.16 |
| TNFRSF11B | CAGTGTCTTTGGTCTCCTGCT | GCAAACCTGAAGAATGCCTCC | S | 79.3 ± 0.80 |
| CD9 | TTCGCCATTGAAATAGCTGCG | TAGTGGATGGCTTTCAGCGTT | S | 81.1 ± 1.89 |
| KLHL24 | CACTGAGTGCTACGATCCTGT | AATCCAGACATCACGGCTGTT | S | 80.0 ± 0.20 |
| HMGCR | ACGTGAACCTATGCTGGTCAG | TGGGGTATCTGTTTCAGCCAC | U | 78.5 ± 0.26 |
| SQLE | ACATTAGAGGAGAAATGCCAAGGA | GAAGGAGGAAGGAAGCTTGCT | S | 79.7 ± 0.21 |
| INSIG1 | TACGCTGATCACGCAGTTTCT | ACAAGGGAGCCAAGAACGAAT | U | 77.6 ± 0.22 |
| BHLHE40 | GAGACGTGACCGGATTAACGA | GTGCTTTCACATGCTTCAAGGT | S | 81.1 ± 1.34 |
| ETV1 | TACCAGATAGCAGCTACCCCA | GAAGGGGATGTTTGGCTCAGA | S | 83.8 ± 0.18 |
| TCOF1 | ATCAGACAGTGAGGAAGCAGC | ACAACTTTTCCAGGGGCTGAA | S | 83.9 ± 0.20 |
| MYBBP1A | GTGACATCCAGACCTGCATCA | AGATCTCCACCAGCACCTCTA | U | 85.5 ± 0.19 |
| EIF5A | CCAAGGTCCATCTGGTTGGTA | TGGAGCAGTGATAGGTACCCA | U | 79.9 ± 0.15 |
| MRPL18 | CAAGTTATCCGGCAGCGAAAC | GGGCTTCACTAGGGATCATGG | S | 83.0 ± 0.19 |
| MRPS10 | ACCTCCAAGGAAAATAGAGCGA | ACATCTGCTGTGCTTCCAGTT | S | 77.6 ± 0.21 |
| SNRPD1 | GAAGAACGGAACACAGGTCCA | CGAATACTCAGCGTTTCCAGC | S | 79.6 ± 1.02 |
| UBXN2A | GTGCCAGTCAGCAGTTTTTGA | GGAACACAGGCTTCGTAGACA | S | 78.3 ± 0.29 |
| EFEMP2 | TTCCAGCTGGGGCCTAACAA | ACACAGGAAGGTCCCATAGGA | U | 84.2 ± 0.20[c] |
| 28S | GAGATTCCCACTGTCCCTACCTAC | CTGATTCCGCCAAGCCC | NA | 81.1 ± 0.22 |
| Sp | CGGATTCACCCAAGGTGATTAT | AACGGTTAGTCGAACCATCAG | NA | 78.2 ± 0.13 |

[a] S, amplicon spans an intron of ≥ ~1 kb or yields no genomic product, so mainly spliced RNA is detected; U, amplicon spans a short intron, so unspliced RNA can also be amplified; NA, not applicable (28S rRNA and spike-in).

[b] Temperature of the single melting peak obtained for each primer pair in a separate validation experiment.

[c] Assay excluded from analysis: amplification efficiency 83.4%, outside the 85–115% acceptance range (Table S19B).

**Table S17. Changes in total and nascent RNA in starved (SS) relative to actively growing (AG) U2OS cells, compared with RNA-seq and EISA.**

*A. Total and nascent RNA*

| Gene | RNA-seq LFC | padj | Total RNA LFC (qPCR) | 95% CI | q | Intronic LFC (RNA-seq) | padj | Nascent RNA LFC (qPCR) | 95% CI | q |
|---|---|---|---|---|---|---|---|---|---|---|
| *ECM Remodelling* | | | | | | | | | | |
| FN1 | 1.77 | $1.3\times10^{-46}$ | 2.73 | 2.45 to 3.01 | $1.2\times10^{-19}$ | 2.47 | $3.2\times10^{-70}$ | 1.09 | 0.69 to 1.50 | $5.9\times10^{-6}$ |
| MMP2 | 2.13 | $<1.0\times10^{-300}$ | 0.96 | 0.68 to 1.24 | $4.4\times10^{-8}$ | 2.07 | $9.1\times10^{-28}$ | 0.99 | 0.58 to 1.40 | $2.9\times10^{-5}$ |
| IGFBP7 | 4.14 | $1.0\times10^{-169}$ | 1.62 | 1.34 to 1.90 | $4.3\times10^{-13}$ | 3.66 | $1.2\times10^{-84}$ | 1.53 | 1.12 to 1.94 | $1.7\times10^{-8}$ |
| TNFRSF11B | 3.92 | $<1.0\times10^{-300}$ | 1.06 | 0.78 to 1.34 | $7.6\times10^{-9}$ | 3.86 | $5.7\times10^{-79}$ | 1.37 | 0.96 to 1.78 | $1.6\times10^{-7}$ |
| CD9 | 1.60 | $1.1\times10^{-156}$ | 1.45 | 1.17 to 1.73 | $6.0\times10^{-12}$ | 1.14 | $1.8\times10^{-9}$ | 0.98 | 0.57 to 1.39 | $3.4\times10^{-5}$ |
| KLHL24 | 2.03 | $1.4\times10^{-127}$ | 1.05 | 0.77 to 1.34 | $8.3\times10^{-9}$ | 1.53 | $7.1\times10^{-10}$ | 1.86 | 1.45 to 2.27 | $3.3\times10^{-10}$ |
| *Lipid/Cholesterol Metabolism* | | | | | | | | | | |
| HMGCR | 1.64 | $1.2\times10^{-136}$ | 0.86 | 0.58 to 1.14 | $3.4\times10^{-7}$ | 1.44 | $2.8\times10^{-6}$ | 1.27 | 0.86 to 1.68 | $6.0\times10^{-7}$ |
| SQLE | 2.12 | $<1.0\times10^{-300}$ | 1.01 | 0.73 to 1.29 | $2.0\times10^{-8}$ | 2.16 | $1.4\times10^{-16}$ | 1.29 | 0.88 to 1.70 | $4.8\times10^{-7}$ |
| INSIG1 | 3.21 | $4.8\times10^{-242}$ | 1.64 | 1.36 to 1.92 | $4.3\times10^{-13}$ | 3.48 | $1.6\times10^{-17}$ | 2.07 | 1.66 to 2.48 | $2.8\times10^{-11}$ |
| *Developmental/Differentiation* | | | | | | | | | | |
| BHLHE40 | 1.80 | $2.7\times10^{-211}$ | 1.48 | 1.20 to 1.76 | $4.1\times10^{-12}$ | 2.47 | $9.1\times10^{-11}$ | 1.70 | 1.29 to 2.10 | $2.5\times10^{-9}$ |
| ETV1 | 1.86 | $1.3\times10^{-13}$ | 0.96 | 0.68 to 1.24 | $4.6\times10^{-8}$ | 2.13 | $6.0\times10^{-4}$ | 1.55 | 1.14 to 1.96 | $1.7\times10^{-8}$ |
| *Ribosome biogenesis* | | | | | | | | | | |
| TCOF1 | −1.20 | $5.8\times10^{-94}$ | −1.25 | −1.53 to −0.97 | $2.1\times10^{-10}$ | −0.49 | 0.015 | −1.22 | −1.63 to −0.81 | $1.1\times10^{-6}$ |
| MYBBP1A | −1.41 | $1.2\times10^{-78}$ | −1.14 | −1.42 to −0.86 | $1.6\times10^{-9}$ | −1.28 | $2.5\times10^{-8}$ | −0.43 | −0.84 to −0.02 | 0.044 |
| *Translation* | | | | | | | | | | |
| EIF5A | −1.16 | $3.1\times10^{-153}$ | −1.20 | −1.48 to −0.92 | $5.4\times10^{-10}$ | −0.13 | 0.742 | −2.26 | −2.67 to −1.85 | $5.2\times10^{-12}$ |
| MRPL18 | −1.05 | $1.1\times10^{-37}$ | −1.00 | −1.28 to −0.71 | $2.5\times10^{-8}$ | −0.11 | 0.791 | −0.78 | −1.19 to −0.37 | $5.6\times10^{-4}$ |
| MRPS10 | −1.04 | $4.0\times10^{-31}$ | −1.37 | −1.65 to −1.09 | $2.2\times10^{-11}$ | −0.47 | 0.161 | −0.54 | −0.95 to −0.13 | 0.013 |
| *Spliceosome* | | | | | | | | | | |
| SNRPD1 | −1.08 | $5.2\times10^{-46}$ | −0.99 | −1.27 to −0.71 | $2.8\times10^{-8}$ | −0.58 | 0.026 | −0.35 | −0.76 to 0.06 | 0.096 |
| *Control (unchanged)* | | | | | | | | | | |
| UBXN2A | −0.09 | 0.508 | −0.10 | −0.38 to 0.18 | 0.489 | 0.22 | 0.481 | 0.20 | −0.21 to 0.61 | 0.327 |

LFC, LFC (SS/AG). RNA-seq: gene-level DESeq2 Wald test (n = 2 per condition). Intronic: DESeq2 on intronic counts, apeglm-shrunken LFC (n = 2 per condition). qPCR: n = 3 independent experiments, total RNA normalised to 28S rRNA per µg, nascent RNA corrected for spike-in recovery, limma moderated paired test with Benjamini–Hochberg correction within each analysis (q).

*B. Nascent-to-total ratio and EISA interaction*

| Gene | Nascent-to-total LFC (qPCR) | 95% CI | q | qPCR class[a] | EISA Dex − Din | FDR | EISA class[b] |
|---|---|---|---|---|---|---|---|
| *ECM Remodelling* | | | | | | | |
| FN1 | −1.73 | −2.33 to −1.12 | 0.017 | Divergent | −0.54 | $1.1\times10^{-5}$ | Destabilisation |
| MMP2 | −0.06 | −0.49 to 0.37 | 0.776 | Concordant | 0.24 | 0.583 | Concordant/background |
| IGFBP7 | −0.18 | −0.83 to 0.47 | 0.639 | Concordant | 0.62 | 0.009 | Stabilisation |
| TNFRSF11B | 0.22 | −0.20 to 0.64 | 0.459 | Concordant | 0.28 | 0.692 | Concordant/background |
| CD9 | −0.56 | −1.08 to −0.05 | 0.152 | Concordant | 0.58 | $4.9\times10^{-4}$ | Stabilisation |
| KLHL24 | 0.71 | 0.09 to 1.34 | 0.152 | Concordant | 0.66 | 0.011 | Stabilisation |
| *Lipid/Cholesterol Metabolism* | | | | | | | |
| HMGCR | 0.31 | −0.36 to 0.99 | 0.459 | Concordant | 0.36 | 0.526 | Concordant/background |
| SQLE | 0.19 | −0.33 to 0.71 | 0.532 | Concordant | 0.16 | 0.944 | Concordant/background |
| INSIG1 | 0.34 | −0.52 to 1.21 | 0.521 | Concordant | 0.31 | 0.508 | Concordant/background |
| *Developmental/Differentiation* | | | | | | | |
| BHLHE40 | 0.12 | −0.37 to 0.61 | 0.653 | Concordant | −0.29 | 0.015 | Small effect |
| ETV1 | 0.50 | −0.06 to 1.06 | 0.205 | Concordant | – | – | Not tested[c] |
| *Ribosome biogenesis* | | | | | | | |
| TCOF1 | −0.06 | −0.66 to 0.54 | 0.811 | Concordant | −0.55 | $3.3\times10^{-6}$ | Destabilisation |
| MYBBP1A | 0.62 | −0.42 to 1.66 | 0.407 | Concordant | 0.00 | 0.871 | Concordant/background |
| *Translation* | | | | | | | |
| EIF5A | −1.15 | −2.24 to −0.06 | 0.152 | Concordant | −0.82 | $1.6\times10^{-4}$ | Destabilisation |
| MRPL18 | 0.12 | −0.45 to 0.69 | 0.670 | Concordant | −0.75 | $4.2\times10^{-5}$ | Destabilisation |
| MRPS10 | 0.74 | 0.13 to 1.35 | 0.152 | Concordant | −0.30 | 0.213 | Concordant/background |
| *Spliceosome* | | | | | | | |
| SNRPD1 | 0.55 | −0.10 to 1.19 | 0.210 | Unresolved | −0.29 | 0.053 | Concordant/background |
| *Control (unchanged)* | | | | | | | |
| UBXN2A | 0.21 | −0.23 to 0.65 | 0.459 | Unresolved | −0.21 | 0.121 | Concordant/background |

Nascent-to-total LFC = (nascent LFC corrected for nascent spike-in recovery) − (total LFC corrected for total spike-in recovery). A negative value indicates that total RNA increased more (or decreased less) than RNA production. EISA Dex − Din has the opposite sign convention.

[a] Concordant, total and nascent RNA both changed significantly in the same direction without a significant nascent-to-total change. Divergent, significant nascent-to-total change. Unresolved, neither. Classes were assigned at $q < 0.05$.

[b] EISA (eisaR, Gaidatzis method, n = 2 per condition). Stabilisation, Dex − Din ≥ 0.5 and FDR < 0.05. Destabilisation, Dex − Din ≤ −0.5 and FDR < 0.05. Small effect, FDR < 0.05 and |Dex − Din| < 0.5. Concordant/background, FDR ≥ 0.05.

[c] Removed by eisaR internal expression filtering.

**Table S18. RNA yield, 28S rRNA and spike-in (Sp) recovery in total and nascent RNA fractions.**

*A. Per-sample values*

| Exp. | Condition | RNA yield (µg) | 28S, total (copies/µg) | 28S, nascent (copies/µg) | Sp, total (copies/µg) | Sp, nascent (copies/µg) | $R_T$ | $R_{NS}$ | CE |
|---|---|---|---|---|---|---|---|---|---|
| 1 | AG | 110 | $1.41\times10^{10}$ | $4.54\times10^{8}$ | $2.34\times10^{8}$ | $1.41\times10^{8}$ | 0.74 | 0.44 | 0.60 |
| 1 | SS | 105 | $1.32\times10^{10}$ | $6.44\times10^{7}$ | $2.01\times10^{8}$ | $1.39\times10^{8}$ | 0.60 | 0.42 | 0.69 |
| 2 | AG | 109 | $1.89\times10^{10}$ | $6.63\times10^{8}$ | $3.15\times10^{8}$ | $1.64\times10^{8}$ | 0.98 | 0.51 | 0.52 |
| 2 | SS | 104 | $1.66\times10^{10}$ | $7.23\times10^{7}$ | $2.61\times10^{8}$ | $1.60\times10^{8}$ | 0.78 | 0.48 | 0.61 |
| 3 | AG | 113 | $2.19\times10^{10}$ | $5.17\times10^{8}$ | $3.63\times10^{8}$ | $1.77\times10^{8}$ | 1.17 | 0.57 | 0.49 |
| 3 | SS | 109 | $1.94\times10^{10}$ | $6.27\times10^{7}$ | $3.52\times10^{8}$ | $1.61\times10^{8}$ | 1.10 | 0.50 | 0.46 |

RNA yield, total RNA recovered per culture from equal numbers of cells. Of each yield, 9.1% was used for the total-RNA fraction and the remainder for nascent-RNA capture. Copies per µg of RNA entering each fraction, from plasmid standard curves. $R_T$ and $R_{NS}$, spike-in recovery in the total and nascent fractions (measured Sp copies per µg relative to the expected copies per µg, $3.5\times10^{10}$ copies divided by the RNA yield of the sample). CE, capture efficiency ($R_{NS}/R_T$). Each value is the mean of 6–7 qPCR wells across three dilutions.

*B. SS/AG comparison (paired by experiment)*

| Measure | SS/AG, exp. 1 / 2 / 3 | SS/AG (mean) | 95% CI | p |
|---|---|---|---|---|
| 28S rRNA, total (per µg) | 0.94 / 0.88 / 0.89 | 0.90 | 0.83–0.98 | 0.034 |
| Sp recovery, total ($R_T$) | 0.82 / 0.79 / 0.94 | 0.85 | 0.68–1.05 | 0.083 |
| Sp recovery, nascent ($R_{NS}$) | 0.94 / 0.93 / 0.87 | 0.91 | 0.83–1.01 | 0.059 |
| Capture efficiency (CE) | 1.15 / 1.17 / 0.93 | 1.08 | 0.79–1.48 | 0.402 |
| 28S rRNA, nascent (recovery-corrected) | 0.15 / 0.12 / 0.14 | 0.13 | 0.10–0.19 | 0.001 |
| 28S rRNA, nascent/total | 0.13 / 0.11 / 0.15 | 0.13 | 0.08–0.19 | 0.002 |
| Total RNA yield (per biological replicate)[a] | 0.95 / 0.95 / 0.96 | 0.96 | 0.94–0.97 | 0.007 |
| 28S rRNA, total (per biological replicate)[a] | 0.89 / 0.84 / 0.85 | 0.86 | 0.80–0.93 | 0.016 |

SS/AG (mean), geometric mean of the three paired ratios. 95% CI and p from a two-sided paired t-test on $\log_2$ values (n = 3).

[a] Equal numbers of AG and SS cells were processed, so per-biological replicate values are proportional to amounts per cell. 28S per culture = 28S copies per µg × RNA yield.

**Table S19. Assay performance: standard curves and amplification efficiencies.**

*A. Plasmid standard curves for 28S rRNA and Sp (one per plate)*

| Assay | Exp. | Levels (wells)[a] | Range, copies/reaction | Slope (95% CI) | Efficiency, % (95% CI)[b] | $R^2$ | Intercept | Status |
|---|---|---|---|---|---|---|---|---|
| 28S | 1 | 6 (16) | $3.20\times10^5$–$2.00\times10^9$ | −3.06 (−3.14 to −2.97) | 112.5 (108.4–116.9) | 0.998 | 38.27 | Refitted with pooled slope −3.25 (102.9%), intercept 39.68[c] |
| 28S | 2 | 6 (16) | $3.20\times10^5$–$2.00\times10^9$ | −3.25 (−3.36 to −3.14) | 103.1 (98.6–108.0) | 0.997 | 39.92 | Accepted |
| 28S | 3 | 6 (17) | $3.20\times10^5$–$2.00\times10^9$ | −3.26 (−3.34 to −3.17) | 102.7 (99.0–106.6) | 0.998 | 39.80 | Accepted |
| Sp | 1 | 6 (16) | $3.20\times10^5$–$2.00\times10^9$ | −3.35 (−3.47 to −3.23) | 98.9 (94.2–104.1) | 0.996 | 39.98 | Accepted |
| Sp | 2 | 6 (16) | $3.20\times10^5$–$2.00\times10^9$ | −3.25 (−3.38 to −3.13) | 102.9 (97.6–108.7) | 0.995 | 39.70 | Accepted |
| Sp | 3 | 6 (16) | $3.20\times10^5$–$2.00\times10^9$ | −3.20 (−3.34 to −3.07) | 105.2 (99.4–111.8) | 0.995 | 39.23 | Accepted |

[a] Levels and wells retained after filtering. No end level deviated by > 1 cycle from the fit through the remaining levels. The well with the largest leave-one-out residual was removed once when that residual exceeded 0.5 cycle, which removed one well on five of the six curves. Six levels, $3.2 \times 10^5$–$2.0 \times 10^9$ copies per reaction, with two wells at the highest level and three at the others.

[b] E = 10^(−1/slope) − 1. 95% CI from the t distribution of the regression slope.

[c] Slope outside −3.6 to −3.1. The curve was refitted with the pooled slope of the other two plates and its intercept was re-estimated (38.27 to 39.68).

*B. Gene assays: amplification efficiency from total-RNA cDNA dilution series*

| Gene | Samples (points)[a] | Slope (95% CI)[b] | Efficiency, % (95% CI)[b] | Pilot efficiency, %[c] | Outcome[d] |
|---|---|---|---|---|---|
| FN1 | 6 (18) | −3.69 (−3.98 to −3.41) | 86.5 (78.4–96.4) | 87.7 | Accepted |
| MMP2 | 6 (18) | −3.64 (−3.80 to −3.47) | 88.4 (83.2–94.2) | 88.2 | Accepted |
| IGFBP7 | 6 (18) | −3.45 (−3.67 to −3.24) | 94.8 (87.3–103.6) | 96.2 | Accepted |
| TNFRSF11B | 6 (18) | −3.65 (−3.78 to −3.51) | 87.9 (83.8–92.5) | 87.3 | Accepted |
| CD9 | 6 (18) | −3.36 (−3.52 to −3.19) | 98.5 (92.3–105.6) | 95.7 | Accepted |
| KLHL24 | 6 (18) | −3.63 (−3.87 to −3.40) | 88.5 (81.3–97.0) | 85.4 | Accepted |
| HMGCR | 6 (18) | −3.73 (−3.90 to −3.56) | 85.4 (80.4–91.0) | 87.7 | Accepted |
| SQLE | 6 (18) | −3.35 (−3.50 to −3.20) | 99.0 (93.2–105.5) | 98.0 | Accepted |
| INSIG1 | 6 (18) | −3.64 (−4.03 to −3.25) | 88.2 (77.0–103.1) | 97.5 | Accepted |
| BHLHE40 | 6 (18) | −3.63 (−3.85 to −3.42) | 88.4 (81.9–96.1) | 92.1 | Accepted |
| ETV1 | 6 (18) | −3.19 (−3.45 to −2.93) | 105.9 (95.1–119.5) | 103.1 | Accepted |
| TCOF1 | 6 (18) | −3.72 (−3.88 to −3.57) | 85.6 (81.0–90.6) | 85.3 | Accepted |
| MYBBP1A | 6 (17) | −3.56 (−3.89 to −3.23) | 91.0 (80.8–104.1) | 91.8 | Accepted |

| Gene | Samples (points)[a] | Slope (95% CI)[b] | Efficiency, % (95% CI)[b] | Pilot efficiency, %[c] | Outcome[d] |
|---|---|---|---|---|---|
| EIF5A | 6 (18) | −3.48 (−3.74 to −3.22) | 93.8 (85.1–104.4) | 93.8 | Accepted |
| MRPL18 | 6 (18) | −3.50 (−3.76 to −3.23) | 93.2 (84.4–104.0) | 95.1 | Accepted |
| MRPS10 | 6 (18) | −3.71 (−3.86 to −3.55) | 86.1 (81.5–91.3) | 90.4 | Accepted |
| SNRPD1 | 6 (18) | −3.59 (−3.83 to −3.34) | 90.0 (82.5–99.1) | 89.4 | Accepted |
| UBXN2A | 6 (18) | −3.62 (−3.81 to −3.43) | 89.0 (83.0–95.8) | 92.0 | Accepted |
| EFEMP2 | 6 (18) | −3.80 (−3.94 to −3.65) | 83.4 (79.3–88.0) | 86.4 | Excluded (< 85%) |

[a] Six total-RNA dilution series (AG and SS, three experiments) at 1:5, 1:25 and 1:125 (1.4 $\log_{10}$ units), one well per dilution. Points passing the Cq ≤ 35, no-template and no-RT criteria were used.

[b] Regression of Cq on $-\log_{10}$ DF with a common slope and sample-specific intercepts. $E = 10^{-1/\text{slope}} - 1$. 95% CI from the t distribution of the slope. The same efficiency was applied to total and nascent RNA.

[c] From the separate primer validation experiment.

[d] Acceptance range 85–115%. Median efficiency 88.5% for all 19 gene assays and 88.7% for the 18 accepted assays.

**Table S20. Results at the relaxed threshold (padj ≤ 0.05, |LFC| ≥ 0.58).**

*A. ATAC-seq differential accessibility*

| Dataset | Tested | Significant (%) | UP (%) | DOWN (%) | Median LFC | IQR |
|---|---|---|---|---|---|---|
| Genome-wide peaks | 81,239 | 78,815 (97.0) | 740 (0.9) | 78,075 (99.1) | -2.36 | -2.74 to -1.90 |
| Promoter peaks | 22,079 | 21,755 (98.5) | 67 (0.3) | 21,688 (99.7) | -2.44 | -2.70 to -2.06 |
| Enhancer peaks | 22,608 | 21,959 (97.1) | 40 (0.2) | 21,919 (99.8) | -2.36 | -2.78 to -1.88 |
| Other peaks | 36,552 | 35,101 (96.0) | 633 (1.8) | 34,468 (98.2) | -2.30 | -2.74 to -1.82 |

*B. Direction of significant changes*

| Dataset | UP (n) | UP (%) | DOWN (n) | DOWN (%) | UN (n) | UN (%) |
|---|---|---|---|---|---|---|
| RNA-seq (genes) | 2,427 | 14.7 | 2,623 | 15.9 | 11,434 | 69.4 |
| Promoter ATAC-seq (peaks) | 67 | 0.3 | 21,688 | 98.2 | 324 | 1.5 |
| Enhancer ATAC-seq (peaks with a gene link) | 40 | 0.2 | 21,919 | 97.0 | 649 | 2.9 |

*C. Promoter-RNA contingency (n = 11,769 genes)*

| ATAC direction | RNA-seq UP | RNA-seq DOWN | RNA-seq UN | Total |
|---|---|---|---|---|
| UP | 3 (0.03%) | 0 (0.00%) | 2 (0.02%) | 5 |
| DOWN | 1,718 (14.60%) | 2,087 (17.73%) | 7,931 (67.39%) | 11,736 |
| UN | 15 (0.13%) | 1 (0.01%) | 12 (0.10%) | 28 |
| Total | 1,736 | 2,088 | 7,945 | 11,769 |

23 genes with mixed peak calls excluded. Concordant: 2,090 genes (17.8%).

*D. Enhancer-RNA contingency (n = 6,602 genes)*

| ATAC direction | RNA-seq UP | RNA-seq DOWN | RNA-seq UN | Total |
|---|---|---|---|---|
| UP | 2 (0.03%) | 1 (0.02%) | 1 (0.02%) | 4 |
| DOWN | 1,109 (16.80%) | 1,073 (16.25%) | 4,361 (66.06%) | 6,543 |
| UN | 9 (0.14%) | 11 (0.17%) | 35 (0.53%) | 55 |
| Total | 1,120 | 1,085 | 4,397 | 6,602 |

30 genes with mixed peak calls excluded. Concordant: 1,075 genes (16.3%).

*E. Odds ratios*

| Region | Genes | OR, ATAC down / RNA down (95% CI) | p | OR, ATAC up / RNA up (95% CI) | p | RNA up: ATAC unchanged vs reduced (%) | OR, RNA up, unchanged vs reduced (95% CI) | p |
|---|---|---|---|---|---|---|---|---|

| | | | | | | | | |
|---|---|---|---|---|---|---|---|---|
| Promoter | 11,769 | 6.92 (1.16-282) | 0.021 | 8.68 (0.99-104) | 0.025 | 53.6 vs 14.6 | 6.73 (2.98-15.39) | $1.8 \times 10^{-6}$ |
| Enhancer | 6,602 | 0.77 (0.40-1.60) | 0.38 | 4.90 (0.36-67.8) | 0.14 | 16.4 vs 16.9 | 0.96 (0.41-1.99) | 1.00 |

*F. Gene classes*

| **Class** | **\|LFC\| ≥ 1, n (%)** | **\|LFC\| ≥ 0.58, n (%)** |
|---|---|---|
| Concordant up | 2 (0.0) | 2 (0.0) |
| Concordant down | 726 (5.8) | 2,210 (17.5) |
| RNA up, ATAC down | 1,078 (8.5) | 1,858 (14.7) |
| RNA down, ATAC up | 0 (0.0) | 0 (0.0) |
| ATAC only | 10,728 (85.0) | 8,470 (67.1) |
| RNA only | 12 (0.1) | 9 (0.1) |
| Mixed ATAC calls | 46 (0.4) | 59 (0.5) |
| No change detected | 27 (0.2) | 11 (0.1) |
| Total | 12,619 | 12,619 |

Genes with RNA and a promoter and/or enhancer value; ATAC direction combines promoter and enhancer calls (mixed when they disagree). Primary-threshold classes are reported in the main text (Results, Classification of regulatory response patterns).

***Section 1. ATAC-seq library analysis.***

The starvation ATAC-seq libraries showed markedly reduced FRiP and TSS enrichment scores relative to actively growing controls (**Table S1A; Figure 1D, E**). Because reduced enrichment metrics can arise either from genuine loss of focal chromatin accessibility or from technical distortion of the ATAC-seq library, we performed a series of complementary analyses to determine whether the starvation-associated reduction in ATAC-seq signal reflected library-quality artefacts, altered fragment architecture, spatial redistribution of signal away from peaks, or a genuine biological reduction in Tn5 accessibility.

*1.1 Standard library metrics and reproducibility:* Basic alignment metrics did not indicate global library failure in the starvation samples. Mapping rate, mapping quality, proper pairing, the number of fragments retained after filtering (22.8 million in SS and 22.1 million in AG libraries on average), the mitochondrial read fraction before filtering (9.7% in SS and 10.6% in AG) and the duplication rate estimated by Picard before deduplication (SS 24.7 ± 1.0%, AG 27.9 ± 2.2%) were comparable between actively growing (AG) and starved (SS) libraries (**Table S1A**). The non-redundant fraction (NRF) and PCR bottlenecking coefficient (PBC1) of the final alignments are 1 by construction, because duplicate and mitochondrial reads had been removed, and they were therefore not used to assess complexity. Complexity was assessed before filtering instead. The Picard-based NRF was 0.72 ± 0.02 in AG and 0.75 ± 0.01 in SS libraries ($p = 0.11$), and the estimated number of distinct library molecules was 52.8 ± 6.6 million in AG and 61.9 ± 3.4 million in SS libraries ($p = 0.13$). Both NRF values are below the preferred ENCODE value of 0.9, which suggests that moderate duplication is shared by all libraries (35.6 and 35.8 million aligned read pairs in AG and SS, including mitochondrial reads) and is not specific to starvation. PBC1 and PBC2 were not calculated before filtering because they require read counts per genomic position that are not part of the alignment report. The reduced FRiP and TSS enrichment scores in SS were therefore not accompanied by poor mapping, a higher duplication rate, a smaller estimated library size, or failure of read-pair recovery.

Peak calling and reproducibility analysis followed the ENCODE ATAC-seq pipeline (MACS2 peak calling and IDR; main Methods). The AG condition yielded 137,830 optimal peaks and the SS condition 52,842 (38% of AG). In both conditions the pooled-pseudoreplicate set was larger than the best true-replicate set (rescue ratios 1.10 and 1.18), so it was taken as the optimal set, and both conditions passed the ENCODE reproducibility criteria (rescue and self-consistency ratios $\leq 2$; **Table S2**). Self-consistency was lower in SS than in AG (ratio 1.68 versus 1.04) because the two SS replicates with the lowest FRiP (Rep1 and Rep3) gave fewer reproducible peaks. The reduced number of reproducible peaks in SS is consistent with lower peak-centred signal in starved chromatin, but does not by itself indicate failure of the sequencing libraries: mapping quality, duplication, estimated library size and mapped fragment architecture all remained acceptable.

*1.2 Fragment Analyzer traces versus mapped insert-size distributions:* Pre-sequencing Fragment Analyzer traces were broader and less discretely nucleosomal in SS than in AG libraries (Supplementary **Figure S2A**). However, Fragment Analyzer traces measure complete library molecules, including approximately 130 bp of sequencing adapter sequence, and are therefore not directly comparable in absolute size to mapped genomic insert-size distributions. For this reason, we assessed biological ATAC-seq fragment architecture using mapped paired-end insert sizes derived from post-trimmed, post-aligned, post-deduplicated and mitochondrially cleaned BAM files.

Mapped insert-size distributions retained interpretable ATAC-seq architecture in both AG and SS conditions (**Figure 1A**; Supplementary **Figure S2B-D; Table S1B**). The dominant short-fragment mode was 52 bp in both conditions, but a mononucleosomal shoulder (at approximately 150-200 bp) was clearly detectable in the mapped insert-size distributions (Supplementary **Figure S2B**) only in AG libraries. These observations indicate that the apparent discordance between conditions is not a sequencing or alignment artefact, but reflects a genuine, partial loss of nucleosome-scale periodicity specifically in the mapped insert-size architecture of SS libraries.

*1.3 Quantitative fragment-class analysis and exclusion of over-tagmentation:* Fragment classes were quantified using standard genomic insert-size windows: nucleosome-free region fragments (NFR, ≤100 bp), mononucleosome-sized fragments (147-247 bp), dinucleosome-sized fragments (315-473 bp), and trinucleosome-sized fragments (500-700 bp), following ATAC-seq conventions [1, 2]. Quantification was performed from Picard CollectInsertSizeMetrics histograms exported through MultiQC (**Table S1B**).

The SS libraries did not show the pattern expected from indiscriminate over-tagmentation. NFR-sized fragments were less abundant in SS than AG libraries (40.7 ± 2.0% versus 47.0 ± 1.4%, $p = 0.015$), whereas mono- and dinucleosomal size-class fractions were preserved and not significantly reduced (mononucleosome: 21.3 ± 1.0% versus 19.7 ± 1.1%, $p = 0.122$; dinucleosome: 8.9 ± 0.7% versus 7.8 ± 0.7%, $p = 0.104$; **Table S1B**). Consequently, the Mono/NFR and Di/NFR ratios were significantly higher in SS than AG libraries (Mono/NFR: 0.526 ± 0.051 versus 0.420 ± 0.036, $p = 0.048$; Di/NFR: 0.221 ± 0.025 versus 0.166 ± 0.019, p = 0.043). Median insert size was also significantly larger in SS than AG libraries (122.7 ± 5.0 bp versus 108.0 ± 4.4 bp, p = 0.020).

This pattern is opposite to that expected under over-tagmentation. Excessive, non-specific Tn5 integration would be expected to increase the proportion of short sub-nucleosomal fragments, decrease the Mono/NFR and Di/NFR ratios, shorten the median insert size, and degrade the ~10 bp Tn5 rotational periodicity. Instead, SS libraries showed a lower NFR fraction, higher nucleosomal-to-NFR ratios, larger median insert sizes, and retained (though attenuated) ~10 bp rotational periodicity. These observations argue against over-tagmentation as the explanation for the reduced FRiP and TSS enrichment scores in SS cells.

*1.4 Tn5-associated rotational periodicity:* To assess whether Tn5 retained sequence-periodic engagement with nucleosomally protected DNA, we analysed the short-fragment insert-size distribution for the characteristic approximately 10 bp rotational periodicity. This periodicity was detectable in both AG and SS libraries (Supplementary **Figure S2C, D; Table S1B**). The dominant period did not differ significantly between conditions (SS: 9.5 bp; AG: 10.9 bp; p = 0.212), although the magnitude of the periodic signal was reduced in SS, consistent with the overall reduction in ATAC-seq signal. Specifically, the FFT periodicity score was lower in SS than AG libraries (1.12 ± 0.19 versus 3.51 ± 0.42, p = 0.004), as was the FFT peak magnitude (0.0146 ± 0.0025 versus 0.0650 ± 0.0041, p < 0.001).

The retention of Tn5 rotational periodicity in SS is important because indiscriminate over-tagmentation would be expected to degrade this structural signature. Thus, although the periodicity signal is weaker in SS, its preservation supports the conclusion that Tn5 integration remained structurally informative in starved chromatin.

*1.5 Assessment of under-tagmentation and general library failure:* Under-tagmentation is also unlikely to explain the SS phenotype. Insufficient tagmentation would be expected to produce poor recovery of short accessible fragments, weak or uninterpretable fragment architecture, compromised library complexity, or inconsistent sample behaviour. Instead, SS libraries retained high mapping quality, proper pairing, duplication and estimated library size comparable to those of AG libraries, and reproducible condition-specific structure (**Table S1A**). In addition, the SS samples separated robustly from AG samples by PCA and showed consistent within-condition behaviour (**Figure 1F, G**). The observed phenotype therefore reflects a reproducible condition-specific shift rather than stochastic loss of signal caused by insufficient transposition.

*1.6 Peak-centred signal distribution and random-background analysis:* To test whether reduced FRiP and TSS enrichment reflected genuine loss of peak-centred accessibility rather than redistribution of signal away from peaks, we quantified ATAC-seq signal at peak summits, local flanking regions and matched random-background regions (Supplementary **Figure S10**; **Table S12**). Summit windows were defined as ±250 bp around consensus peak summits, or peak midpoints where summit coordinates were unavailable. Local flanks were defined as 1,000-2,000 bp on each side of the summit; peaks whose flanks overlapped other consensus peaks or ENCODE blacklist intervals were excluded, leaving 72,497 of the 88,589 consensus peaks. Random-background regions were sampled genome-wide, excluding consensus peaks, blacklisted regions and assembly gaps. Fragment counts were scaled to counts per million mapped fragments (CPM) and expressed per base pair. AG and SS were compared using two-tailed Welch's t-tests on replicate-level means (n = 3 per condition).

Summit signal was reduced 5.5-fold in SS relative to AG (0.0020 ± 0.0005 versus 0.0112 ± 0.0015 CPM per bp; SS/AG = 0.18; p = 0.0057; **Table S12A**), and the proportion of summit-proximal signal concentrated within the summit window fell from 89.4% in AG to 49.1% in SS (p = 0.002; **Table S12B**). The reduction was of similar magnitude at promoter (log2 SS/AG -2.60), enhancer (-2.42) and other peaks (-2.29), and extended to housekeeping-gene promoters (-2.69) and distal CTCF-motif sites (-2.10).

Local flank signal increased 1.61-fold (from 0.00032 ± 0.00005 to 0.00052 ± 0.00001 CPM per bp; p = 0.014) and random-background signal 2.14-fold (from 0.00019 ± 0.00004 to 0.00041 ± 0.00002 CPM per bp; p = 0.0042; **Table S12A**), both remaining small in absolute terms relative to summit signal. Because libraries from the two conditions contained similar numbers of mapped fragments (22.1 and 22.8 million on average), fragments no longer captured in peaks are necessarily distributed over the rest of the genome: with the fraction of fragments in each condition's own IDR peaks falling from 56.4% to 7.2%, the fraction of fragments outside peaks rises from 43.6% to 92.8%, a 2.13-fold increase that matches the observed rise in background signal. The background increase is therefore the counterpart of the loss of peak-centred signal rather than evidence of a separate increase in non-specific Tn5 insertion. Flank signal rose least at promoters (1.27-fold, p = 0.084) and housekeeping-gene promoters (1.23-fold, p = 0.14) and most at distal CTCF-motif sites (1.98-fold), consistent with the loss of broader promoter-associated accessibility in SS offsetting the background increase at promoters (Supplementary Figure S7).

Ratio-based metrics showed the same pattern: the summit/flank ratio decreased from 35.4 in AG to 3.9 in SS (p = 0.026), and the summit/random-background ratio from 60.6 to 5.0 (p = 0.032; **Table S12C**). This analysis shows that the dominant change in SS is a loss of focal, summit-centred accessibility. On its own, however, it cannot distinguish a genuine global reduction in accessibility from a uniformly lower signal-to-noise ratio in SS libraries; the arguments that bear on this distinction are summarised in Section 1.9 and in the main Discussion.

*1.7 Normalisation and constitutively accessible sites:* Normalisation to fragments in consensus peaks equalises the total peak-associated signal between conditions and therefore removes any change common to all peaks by construction. Because most peaks lost signal in SS, the primary differential analysis was normalised to library size, which expresses the signal at each peak per mapped fragment (main Methods, **Table 2**). Under this normalisation, the median LFC was -2.34 across all tested peaks, -2.56 at housekeeping-gene promoters (n = 4,232) and -2.17 at distal CTCF-motif sites (n = 5,721), classes of sites usually regarded as constitutively accessible (**Figure 2C**). Aggregate CPM metaplots centred on peak summits showed the same reduction for all peaks (log2 SS/AG at the summit -2.57), housekeeping-gene promoters (-2.72) and distal CTCF-motif sites (-2.46) (Supplementary **Figure S7**). Normalisation to fragments in consensus peaks gave per-peak LFCs that correlated with the library-size values at r = 0.999 and were shifted by 2.42 log2 units (**Table S3**), which is the log2 ratio of the fraction of fragments in consensus peaks in AG and SS (56.5% and 10.6%). The two normalisations therefore describe the same per-peak changes on different scales. Library-size normalisation reports the change per mapped fragment, whereas consensus-peak normalisation reports each peak's share of the remaining peak-associated signal, so that a peak losing less signal than the peak pool can appear increased on that scale. The per-fragment scale is not calibrated per cell, because no exogenous reference was included (main Discussion).

*1.8 Peak provenance and exclusion of global decompaction as the dominant explanation:* Peak-provenance analysis provides an additional test of the global decompaction model. If starvation caused broad chromatin decompaction, one would expect increased accessibility at many loci, elevated or preserved FRiP and TSS enrichment, increased NFR-sized fragments, and a substantial number of starvation-specific peaks. Instead, only 1.9% of genome-wide consensus peaks were SS-only, compared with 63.7% AG-only and 34.4% shared peaks, and 94.9% of the peaks detected in SS were also detected in AG (**Table 1; Figure 1H**). SS-only peaks were equally rare at promoters (0.8%) and enhancers (0.7%). Promoter peaks were, however, more often retained as detectable peaks in SS (41.3% AG-only, 0.8% SS-only and 57.9% shared) than enhancer peaks (67.5% AG-only, 0.7% SS-only and 31.8% shared).

Together with the lower NFR fraction, sharply reduced FRiP and TSS enrichment, and widespread decrease in peak-centred signal, this peak-provenance structure is inconsistent with global decompaction as the dominant explanation for the SS library phenotype. Local accessibility gains may still occur at specific loci, but they cannot account for the genome-wide pattern.

*1.9 Overall interpretation:* Taken together, the ATAC-seq library analyses support four conclusions. First, SS libraries retained high-quality mapping and pairing, duplication and estimated library size comparable to AG libraries, and reproducible condition-specific structure, arguing against general library failure. Second, mapped insert-size distributions retained characteristic ATAC-seq architecture and Tn5-associated rotational periodicity, arguing against loss of interpretable tagmentation structure. Third, quantitative fragment-class analysis showed reduced NFR fraction, increased nucleosomal-to-NFR ratios and increased median insert size in SS, which is opposite to the pattern expected from over-tagmentation. Fourth, peak-centred analyses showed a strong loss of summit accessibility, extending to constitutively accessible sites, with increases in flank and random-background signal of the size expected when a similar number of fragments is no longer concentrated in peaks.

Therefore, the reduced FRiP and TSS enrichment scores in SS cells are best interpreted as consequences of genuine genome-wide reduction in Tn5 accessibility at canonical accessible regions. The analysis does not by itself prove physical chromatin compaction, because ATAC-seq measures transposase accessibility rather than chromatin density directly, and without an exogenous reference it cannot fix the exact magnitude of the reduction. However, over-tagmentation, under-tagmentation, general library failure and global decompaction are not consistent with the observed library architecture. The most parsimonious interpretation is a starvation-associated, compaction-compatible reduction in chromatin accessibility, with potential contributions from other biological mechanisms that reduce Tn5 access to chromatin.

***Section 2. Statistical considerations in n=2 RNA-seq validation.***

RNA-seq was performed on two biological replicates per condition. This design limits the power to detect genes with small or highly variable changes in expression. However, in our experiments, for most genes variability between biological replicates was low (median CV~ 15%), which partly offsets this limitation. Pathway-level analyses using gene set enrichment are inherently robust to lower sample sizes as they aggregate signal across multiple genes.

*2.1 Dispersion analysis:* DESeq2 dispersion estimates had a median of 0.023 and a mean of 0.22, and 16.5% of genes had a dispersion above 0.5. These values describe variability between biological replicates and do not isolate technical variance. With two replicates per condition, DESeq2 relies on empirical Bayes shrinkage of the gene-wise estimates towards the fitted mean-dispersion trend (**Figure S3A**).
*2.2 Statistical power:* To estimate the sensitivity of the analysis, we simulated negative binomial counts for 30,000 genes with a mean normalised count of 100, dispersions drawn to match the observed distribution, and true changes in 25% of genes, and analysed them with DESeq2 as in the main analysis. With two replicates per condition, the full selection rule (padj $\leq 0.05$ and $|LFC| \geq 1$) detected 3%, 40%, 81% and 87% of genes with true changes of $|LFC| = 0.5$, 1, 1.5 and 2, respectively (**Figure S3B**). Power fell with dispersion: for a true two-fold change it was 48% for genes with dispersion $\leq 0.05$, 36% for dispersion 0.05 to 0.5 and 16% for dispersion above 0.5. Because the rule requires an estimated |LFC| of at least 1, a gene with a true two-fold change passes it only about half of the time even when the test itself is well powered. The 2,664 DEGs are therefore enriched for large changes in genes with low variability, and smaller or more variable changes are likely to be under-represented. These estimates agree with benchmark data showing that experiments with few replicates detect most genes with large changes but miss many genes with small changes [3].
*2.3 Comparison with three replicates:* Repeating the simulation with three replicates per condition, using the same genes and true changes, increased the power of the full selection rule only slightly (3%, 44%, 84% and 89% at $|LFC| = 0.5$, 1, 1.5 and 2), because the fold-change threshold, rather than replication, limits detection of genes with changes close to two-fold (**Figure S3B**). Without the fold-change threshold, the third replicate would have increased the detection of a true two-fold change from 63% to 78% and of a true change of $|LFC| = 0.5$ from 3% to 18%. The main cost of using two replicates is therefore a higher proportion of false calls among the selected genes, which in the simulation was about 9% to 16% with two replicates and 5% to 10% with three. For this reason, pathway-level conclusions were based on ranked enrichment across all genes, and the direction of change was confirmed by RT-qPCR for all genes tested (Supplementary **Table S17**).
*2.4 Sensitivity to thresholds:* Relaxing the selection thresholds to $|LFC| \geq 0.5$ and padj $< 0.1$ increased the number of DEGs from 2,664 to 6,572 and reduced the proportion of increased genes from 64.5% to 50.5% (**Figure S3C**). Across combinations of thresholds (padj 0.01 to 0.15, |LFC| 0.5 to 2.0), the proportion of increased genes rose with the fold-change threshold, from about 48% at $|LFC| \geq 0.5$ to 81% at $|LFC| \geq 2.0$, at every padj threshold (**Figure S3D**). The excess of increased genes is therefore confined to larger changes, whereas increases and decreases of smaller magnitude are similarly frequent. Because the test has the same power for increases and decreases of equal size at equal expression and dispersion, the asymmetry is unlikely to be produced by the selection rule itself, although differences in expression level or variability between increased and decreased genes could contribute. The directional asymmetry reported in the main text therefore applies to changes of at least two-fold.

***Section 3. Cellular poly(A)+ RNA content in starved and actively growing cells***

DESeq2 differential expression analysis uses median-of-ratios size factors, which recover changes in the relative composition of the transcriptome rather than absolute transcript numbers per cell. If starvation substantially reduced the total cellular mRNA pool, libraries prepared from equal RNA inputs would rescale this reduction, causing genes with unchanged absolute per-cell abundance to appear upregulated. We therefore examined whether starvation reduced cellular poly(A)+ RNA content, potentially contributing to apparent relative upregulation.

Poly(A)+ RNA was purified from counted actively growing (AG) and starved (SS) U2OS cells using the GenElute Direct mRNA Miniprep Kit (DMN10, Sigma-Aldrich) as per manufacturer instruction. Three independent cultures per condition were processed in parallel, using $1.9 \times 10^7$ AG or $7.4 \times 10^6$ SS cells per preparation. Yield was expressed as pg per cell (Supplementary **Table S11A**).

In a separate experiment, three additional cultures per condition were analysed by purifying poly(A)+ RNA from 200 µg of total RNA. Yield was expressed as a percentage of the input, providing a complementary measurement independent of cell counting (Supplementary **Table S11B**).

Eluate concentrations were measured using the Quant-iT RNA Assay Kit, Broad Range (Invitrogen), calibrated against the supplied standards. Samples were measured undiluted and at 1:5, 1:25 and 1:125 dilutions.

Potential contamination of the purified poly(A)+ fraction by rRNA was assessed by RT-qPCR using primers and probes against 18S and 28S rRNA [4]. Neither 18S nor 28S rRNA was detected in the eluates within 40 cycles, whereas matched total RNA controls amplified as expected [4].

Conditions were compared using unpaired, two-sided Student's t-tests. SS/AG ratios and their 95% confidence intervals were calculated on the $\log_2$ scale. Equivalence was assessed using two one-sided tests (TOST), with ratio bounds of 0.75–1.25 (Supplementary **Table S11C**).

Poly(A)+ RNA content was $0.737 \pm 0.053$ pg per cell in AG and $0.744 \pm 0.044$ pg per cell in SS (mean ± SD; SS/AG = 1.01, 95% CI 0.87–1.17; $p = 0.86$). Poly(A)+ RNA represented $8.06 \pm 0.58\%$ of total RNA in AG and $8.10 \pm 0.29\%$ in SS (SS/AG = 1.01, 95% CI 0.89–1.14; $p = 0.91$). Both measurements met the equivalence criterion (TOST $p = 0.008$ and $0.005$, respectively).

These results argue against substantial poly(A)+ RNA depletion during starvation. Under a simple global-rescaling model, the lower per-cell confidence limit of 0.87 would correspond to a shift of approximately 0.20 $\log_2$ units, substantially below the $|LFC| \geq 1$ threshold used to identify DEGs. The measurements therefore reduce concern that the observed RNA-seq upregulation reflects a contraction of the cellular mRNA pool, while leaving the RNA-seq results as relative abundance estimates.

**Section 4. ATAC-RNA steady-state integration pipeline.**

Differential chromatin accessibility and differential gene expression statistics were generated upstream as described in the main Methods (ATAC-seq data analysis; RNA-seq data analysis). The integration pipeline operates on the resulting DESeq2 tables and relates changes in chromatin accessibility to changes in mRNA abundance at the gene level, following Kiani et al. [5]. It addresses the following objectives:

• quantify the genome-wide relationship between accessibility changes and mRNA changes, with confidence intervals;
• compare the contributions of promoter-proximal and distal (enhancer) elements;
• test whether mRNA changes depend on the size of the accessibility change, independently of accessibility before starvation;
• describe directional agreement between accessibility and mRNA changes;
• compare accessibility changes of the genes driving enriched pathway programmes with those of all other genes.

The workflow is implemented as three R scripts that build gene-level tables, perform the statistical integration, and analyse pathway programmes. Pipeline architecture is outlined in Supplementary **Figure S6** and described below.

**Input datasets**

• RNA-seq differential expression table (DESeq2, SS vs AG; gene, baseMean, LFC, padj) and the DESeq2 object, from which apeglm-shrunken LFCs and normalised counts are obtained.
• ATAC-seq differential accessibility table for all consensus peaks (DESeq2 with library-size size factors; unshrunken and apeglm-shrunken LFCs, padj, normalised means per condition, near-background flag).
• Peak annotation: promoter peaks with their assigned gene (transcript-level TSS windows, -2,000 to +500 bp; gene of the closest transcript TSS) and enhancer peaks with their GeneHancer target genes and association scores.
• For the programme analysis, the leading-edge genes of enriched pathway categories and the programme definitions (Supplementary **Table S7**).

**Script 1: Gene-level tables.**

1. RNA-seq genes with baseMean $\geq 10$ are retained. Gene symbols in both datasets are updated to current official symbols (org.Hs.eg.db).
2. Peaks are linked to genes: each promoter peak to its assigned gene; each enhancer peak to its three highest-ranked GeneHancer target genes.
3. Each peak is called as gained, lost or unchanged at the primary threshold (padj $\leq 0.05$ and $|LFC| \geq 1$) and at the relaxed threshold ($|LFC| \geq 0.58$).
4. Peak values are summarised per gene: the promoter value is the median LFC of the gene's promoter peaks; the enhancer value is the GeneHancer-score-weighted mean LFC of its linked enhancer peaks. Shrunken and unshrunken values are both kept. A gene's promoter (or enhancer) is called reduced if at least one of its peaks is lost and none gained, increased in the converse case, unchanged if no peak changes, and mixed otherwise.
5. Baseline accessibility is computed as log2(median normalised fragments + 1) of the gene's promoter peaks in AG cells.
6. Output: one table per gene with RNA values, promoter and enhancer values, calls and baseline accessibility; peak-gene pair tables.

**Script 2: Statistical integration.**

1. Correlation: Pearson and Spearman correlations between RNA and promoter (or enhancer) shrunken LFCs across all matched genes, without filtering on significance; 95% confidence

intervals from 10,000 bootstrap resamples of genes; promoter and enhancer correlations compared by paired bootstrap in genes with both values. Sensitivity analyses: unshrunken LFCs, and genes whose promoter peaks are above background in SS.

2. Directional agreement: gene-level 3 × 3 contingency tables (RNA up, down, unchanged × accessibility increased, reduced, unchanged) at both thresholds; Fisher's exact test with odds ratios and 95% confidence intervals. Odds ratios with an empty cell are reported as not estimable.

3. Baseline accessibility: AG promoter accessibility compared between gene groups (accessibility and RNA categories) by the Wilcoxon rank-sum test.

4. Quintile analysis: genes divided into quintiles of promoter (and, separately, enhancer) LFC; trends in RNA LFC (Spearman correlation) and in the proportions of genes with increased or decreased RNA (Cochran-Armitage test); odds ratio for increased RNA in the fifth vs the first quintile; linear and logistic regression with quintile and baseline accessibility as covariates.

5. Regression: RNA LFC ~ promoter + enhancer LFC (genes with both values); the same model with baseline promoter accessibility; and RNA ~ promoter + baseline accessibility across all genes with a promoter value. Standardised coefficients and $R^2$ are reported.

6. Within-condition relationship: correlation of promoter accessibility with RNA abundance (log2 normalised mean + 1) in AG and in SS cells.

7. Gene classes: genes classified by the combined direction of their RNA and accessibility changes (concordant, discordant, accessibility only, RNA only, mixed, no change).

8. Secondary analyses: correlations restricted to genes with significant changes, and a list of high-confidence concordant genes (padj $\leq 0.01$ and $|LFC| \geq 2$ in RNA-seq and in at least one promoter or enhancer peak, same direction; Supplementary **Table S6**). Because selection on significance inflates the apparent coupling, these are reported for illustration only.

**Script 3: Pathway programmes.**

1. ATAC columns of Table S7 are rebuilt from the gene-level tables (promoter and enhancer LFC, smallest padj, normalised means, number of peaks, gene-level calls).

2. Heatmaps of RNA, promoter and enhancer LFCs for the leading-edge genes of each category (**Figure 8A, B**).

3. For each category and each adaptive programme, promoter LFCs are compared with those of all other genes by the Wilcoxon rank-sum test and by linear regression adjusted for baseline promoter accessibility, with Benjamini-Hochberg correction (**Figure 8C**).

4. Gene-level accessibility values are extracted for the genes measured by RT-qPCR (Figure 9A).

**Outputs**

**Tables 5-8** and Supplementary **S6, S7, S8 and S20**; **Figures 6, 7, 8 and 9A** and Supplementary **Figures S12 and S13**.

**Integration workflow**

The computational workflow can be summarised as the following sequence (Supplementary **Figure S6**).

Step 1. Load the RNA-seq and ATAC-seq DESeq2 results; retain RNA genes with baseMean ≥ 10; update gene symbols.
Step 2. Link promoter peaks to genes (closest transcript TSS) and enhancer peaks to their top three GeneHancer targets; call peaks as gained, lost or unchanged.
Step 3. Summarise per gene: promoter median and enhancer GeneHancer-weighted mean LFC; gene-level calls; baseline AG promoter accessibility.
Step 4. Quantify association: correlations with bootstrap confidence intervals; contingency tables with odds ratios; regression with and without baseline accessibility.
Step 5. Test dependence on the size of the change: quintiles of promoter (and enhancer) change with trend tests and baseline adjustment.
Step 6. Programmes: compare promoter changes of programme genes with all other genes, adjusted for baseline accessibility; build Supplementary **Table S7** and **Figures 8 and 9A**.
Step 7. Secondary: restricted correlations and the high-confidence concordant gene list.

***Section 5. Production of labelled spike-in RNA***

***Spike-in plasmid construction***

The spike-in plasmid (pSpike) was a kind gift from Dr Fuquan Liu (QUB) and used as a template for synthesis of external normalisation reference for quantitative RT-PCR analysis of nascent and total RNA. The construct comprises a T7 RNA polymerase promoter followed by three elements: a 297 bp fragment of the *E.coli* K-12 gene yjgL (locus tag b4253; RefSeq NC_000913.3), a 199 bp fragment of the *A. thaliana gene* AT5G25754 (RefSeq NM_147917.4) and a synthetic poly(A) tract of 40 nucleotides, cloned into pGEM-T Easy (Promega). A 13 nt sequence (nucleotides 539 to 551) lies between the Arabidopsis fragment and the poly(A) tract. After linearisation with SpeI, T7 transcription gives a run-off transcript of approximately 556 nt. The construct was confirmed by DNA sequencing (sequence provided below).

***In vitro transcription and 5-EU labelling***

The pSpike plasmid was linearised downstream of the poly(A) tail by Spe I digestion to generate a run-off template. *In vitro* transcription was performed using T7 RNA polymerase (Promega). Each 100 µL reaction contained 5 µg linearised pSpike plasmid, 40 U T7 RNA polymerase, 100 U RNasin ribonuclease inhibitor, 10 mM DTT and 1× transcription buffer (40 mM Tris–HCl, pH 7.9, 10 mM NaCl, 6 mM $MgCl_2$ and 2 mM spermidine). ATP, CTP and GTP were each supplied at 0.5 mM. The combined concentration of UTP and 5-ethynyl-UTP (5-EUTP) was 0.5 mM, with a 5-EUTP molar ratio of 1:20 (23.8 µM 5-EUTP and 476.2 µM UTP). Reactions were assembled at room temperature and incubated at 37°C for 2 h. Template DNA was subsequently removed by adding 5 U RNase-Free DNase (Promega, M6101) and incubating at 37°C for 15 min. Reaction was terminated by the addition of TRIzol reagent (ThermoFisher). The 5-EU-labelled RNA was subsequently purified, with recovery by ethanol precipitation. The RNA pellet was dissolved in water, divided into aliquots and stored at −80 °C. Each aliquot was thawed only once, immediately before use.

***pSpike sequence***

The complete pSpike construct sequence, including pGEM-T Easy (Promega)vector backbone, T7 promoter, insert elements (yjgL + at5g25754 + poly(A)), and multiple cloning site, is provided below. Regions corresponding to the three insert elements are as follows: yjgL fragment, nucleotides 43-339; at5g25754 fragment, nucleotides 340-538; poly(A) tail, nucleotides 552-591.

GTAAAACGACGGCCAGTGAATTGTAATACGACTCACTATAGGGCGAATTGGGCC
CAACGAAGCGGGTAGAAATATGATGGCTCATATTAAGTCCTCATCTTATTCTAAA
GATACTAATGGCAATGAAAAAATGAAGTTTTACATGAATAATCCTGTAGGGGAA
CGAGCGGATTCACCCAAGGTGATTATAGAAATTTCACTTTCCACTATCACTACTA
TGGGGACTCGTCAAGGACATACAGCCATTATATTTCCACAACCTGATGGTTCGAC
TAACCGTTATGAAGGGAAGTCCTTTGAAAGAAAAGATGAGAGTTCATTACACCT
GATTACTAACAAGGTTCTGGCGTGTAAGCTTGAAGGTGTGATCAACGACATGGAT
CGTGTCAAGCTGGAATGACGACTTTTCTGTGTTATACCTAATCATCAATGCCCAA
TCTGATATGATGCTTTCTCTTATAAACCATATGTTTACTTAAAGCCAAACACTTTT
TTGAGTAGCTGTCTTCATGTGGTCAAATGATAACCATAAATCATCTTTCTTATGGA
TTAAAAAAAAAAAAAAAAAAAAAAAAAAAAAAAAAAAAAAAATTTACTAGTG
AATTCGCGGCCGCCTGCAGGTCGACCATATGGGAGAGCTCCCAACGCGTTGGAT
GCATAGCTTGAGTATTCTATAGTGTCACCTAAATAGCTTGGCGTAATCATGGTCA
TAGCTGTTTCCTGTGTGAAATTGTTATCCGCTCACAATTCCACACAACATACGAG
CCGGAAGCATAAAGTGTAAAGCCTGGGGTGCCTAATGAGTGAGCTAACTCACAT
TAATTGCGTTGCGCTCACTGCCCGCTTTCCAGTCGGGAAACCTGTCGTGCCAGCT
GCATTAATGAATCGGCCAACGCGCGGGGAGAGGCGGTTTGCGTATTGGGCGCTC
TTCCGCTTCCTCGCTCACTGACTCGCTGCGCTCGGTCGTTCGGCTGCGGCGAGCG
GTATCAGCTCACTCAAAGGCGGTAATACGGTTATCCACAGAATCAGGGGATAAC
GCAGGAAAGAACATGTGAGCAAAAGGCCAGCAAAAGGCCAGGAACCGTAAAAA
GGCCGCGTTGCTGGCGTTTTTCCATAGGCTCCGCCCCCCTGACGAGCATCACAAA
AATCGACGCTCAAGTCAGAGGTGGCGAAACCCGACAGGACTATAAAGATACCAG
GCGTTTCCCCCTGGAAGCTCCCTCGTGCGCTCTCCTGTTCCGACCCTGCCGCTTAC
CGGATACCTGTCCGCCTTTCTCCCTTCGGGAAGCGTGGCGCTTTCTCATAGCTCAC
GCTGTAGGTATCTCAGTTCGGTGTAGGTCGTTCGCTCCAAGCTGGGCTGTGTGCA
CGAACCCCCCGTTCAGCCCGACCGCTGCGCCTTATCCGGTAACTATCGTCTTGAG
TCCAACCCGGTAAGACACGACTTATCGCCACTGGCAGCAGCCACTGGTAACAGG
ATTAGCAGAGCGAGGTATGTAGGCGGTGCTACAGAGTTCTTGAAGTGGTGGCCT
AACTACGGCTACACTAGAAGAACAGTATTTGGTATCTGCGCTCTGCTGAAGCCAG
TTACCTTCGGAAAAAGAGTTGGTAGCTCTTGATCCGGCAAACAAACCACCGCTG
GTAGCGGTGGTTTTTTTGTTTGCAAGCAGCAGATTACGCGCAGAAAAAAAGGATC
TCAAGAAGATCCTTTGATCTTTTCTACGGGGTCTGACGCTCAGTGGAACGAAAAC
TCACGTTAAGGGATTTTGGTCATGAGATTATCAAAAAGGATCTTCACCTAGATCC
TTTTAAATTAAAAATGAAGTTTTAAATCAATCTAAAGTATATATGAGTAAACTTG
GTCTGACAGTTACCAATGCTTAATCAGTGAGGCACCTATCTCAGCGATCTGTCTA
TTTCGTTCATCCATAGTTGCCTGACTCCCCGTCGTGTAGATAACTACGATACGGG
AGGGCTTACCATCTGGCCCCAGTGCTGCAATGATACCGCGAGACCCACGCTCACC
GGCTCCAGATTTATCAGCAATAAACCAGCCAGCCGGAAGGGCCGAGCGCAGAAG
TGGTCCTGCAACTTTATCCGCCTCCATCCAGTCTATTAATTGTTGCCGGGAAGCTA
GAGTAAGTAGTTCGCCAGTTAATAGTTTGCGCAACGTTGTTGCCATTGCTACAGG
CATCGTGGTGTCACGCTCGTCGTTTGGTATGGCTTCATTCAGCTCCGGTTCCCAAC
GATCAAGGCGAGTTACATGATCCCCCATGTTGTGCAAAAAAGCGGTTAGCTCCTT
CGGTCCTCCGATCGTTGTCAGAAGTAAGTTGGCCGCAGTGTTATCACTCATGGTT
ATGGCAGCACTGCATAATTCTCTTACTGTCATGCCATCCGTAAGATGCTTTTCTGT
GACTGGTGAGTACTCAACCAAGTCATTCTGAGAATAGTGTATGCGGCGACCGAG
TTGCTCTTGCCCGGCGTCAATACGGGATAATACCGCGCCACATAGCAGAACTTTA

AAAGTGCTCATCATTGGAAAACGTTCTTCGGGGCGAAAACTCTCAAGGATCTTAC
CGCTGTTGAGATCCAGTTCGATGTAACCCACTCGTGCACCCAACTGATCTTCAGC
ATCTTTTACTTTCACCAGCGTTTCTGGGTGAGCAAAAACAGGAAGGCAAAATGCC
GCAAAAAAGGGAATAAGGGCGACACGGAAATGTTGAATACTCATACTCTTCCTT
TTTCAATATTATTGAAGCATTTATCAGGGTTATTGTCTCATGAGCGGATACATATT
TGAATGTATTTAGAAAAATAAACAAATAGGGGTTCCGCGCACATTTCCCCGAAA
AGTGCCACCTGATGCGGTGTGAAATACCGCACAGATGCGTAAGGAGAAAATACC
GCATCAGGAAATTGTAAGCGTTAATATTTTGTTAAAATTCGCGTTAAATTTTTGTT
AAATCAGCTCATTTTTTAACCAATAGGCCGAAATCGGCAAAATCCCTTATAAATC
AAAAGAATAGACCGAGATAGGGTTGAGTGTTGTTCCAGTTTGGAACAAGAGTCC
ACTATTAAAGAACGTGGACTCCAACGTCAAAGGGCGAAAAACCGTCTATCAGGG
CGATGGCCCACTACGTGAACCATCACCCTAATCAAGTTTTTTGGGGTCGAGGTGC
CGTAAAGCACTAAATCGGAACCCTAAAGGGAGCCCCCGATTTAGAGCTTGACGG
GGAAAGCCGGCGAACGTGGCGAGAAAGGAAGGGAAGAAAGCGAAAGGAGCGG
GCGCTAGGGCGCTGGCAAGTGTAGCGGTCACGCTGCGCGTAACCACCACACCCG
CCGCGCTTAATGCGCCGCTACAGGGCGCGTCCATTCGCCATTCAGGCTGCGCAAC
TGTTGGGAAGGGCGATCGGTGCGGGCCTCTTCGCTATTACGCCAGCTGGCGAAA
GGGGGATGTGCTGCAAGGCGATTAAGTTGGGTAACGCCAGGGTTTTCCCAGTCA
CGACGTT
