## Supplementary material for "Genome-wide loss of chromatin accessibility coexists with selectively elevated RNA abundance in nutrient-starved U2OS cells": Abbreviation list

**List of Abbreviations**

AG – Actively Growing
ATAC-seq – Assay for Transposase-Accessible Chromatin using sequencing
AU – Arbitrary Units
BAM/BAI – Binary Alignment Map / BAM Index
BMP – Bone Morphogenetic Protein
BP – Biological Process
CC – Cellular Component
CE – Capture Efficiency
ChIP-seq – Chromatin Immunoprecipitation followed by sequencing
CI – Confidence Interval
CP – Canonical Pathways
CPM – Counts Per Million
Cq – Quantification Cycle (reported as Cp, Crossing Point)
CTCF – CCCTC-Binding Factor
DAPs – Differentially Accessible Peaks
DEGs – Differentially Expressed Genes
Dex / Din – Exonic / Intronic $Log_2$ Fold Change (EISA)
DF – Dilution Factor
DMEM – Dulbecco's Modified Eagle Medium
ECM – Extracellular Matrix
EISA – Exon–Intron Split Analysis
ENCODE – Encyclopedia of DNA Elements
5-EU – 5-Ethynyluridine
FACS – Fluorescence-Activated Cell Sorting
FDR – False Discovery Rate
FFT – Fast Fourier Transform
fGSEA – Fast Gene Set Enrichment Analysis
FRiP – Fraction of Reads in Peaks
GEO – Gene Expression Omnibus
GO – Gene Ontology
GRO-seq – Global Run-On sequencing
GSEA – Gene Set Enrichment Analysis
GTPase – Guanosine Triphosphatase
HK – Housekeeping (Genes)
IDR – Irreproducible Discovery Rate
IQR – Interquartile Range
LFC – $Log_2$ Fold Change
MAP – Maximum A Posteriori
MAPQ – Mapping Quality

MF – Molecular Function
MIQE – Minimum Information for Publication of Quantitative Real-Time PCR Experiments
MSigDB – Molecular Signatures Database
mTOR – Mechanistic Target of Rapamycin
NES – Normalized Enrichment Score
NFR – Nucleosome-Free Region
NRF – Non-Redundant Fraction
NS – Not Significant
Nt / Np – Best True-Replicate / Pooled-Pseudoreplicate Peak Number (IDR)
NTC – No-Template Control
OR – Odds Ratio
PBC1 – PCR Bottlenecking Coefficient 1
PBS – Phosphate-Buffered Saline
PC – Principal Component
PCA – Principal Component Analysis
PCR – Polymerase Chain Reaction
PI – Propidium Iodide
Pol I – RNA Polymerase I
PRC1 / PRC2 – Polycomb Repressive Complex 1 / 2
PRO-seq – Precision Run-On sequencing
QC – Quality Control
qPCR – Quantitative PCR
QUB – Queen's University Belfast
RNA-seq – RNA sequencing
RNS / RT – Spike-in Recovery, Nascent / Total RNA Fraction
RT-qPCR – Reverse-Transcription Quantitative PCR
SD – Standard Deviation
SE – Standard Error
SEM – Standard Error of the Mean
SF – Size Factor
SLAM-seq – Thiol(SH)-Linked Alkylation for the Metabolic Sequencing of RNA
Sp – Spike-in RNA
SS – Starved State
TF – Transcription Factor
TFT – Transcription Factor Targets
TGF-β – Transforming Growth Factor Beta
TMM – Trimmed Mean of M-values
TOST – Two One-Sided Tests
TSS – Transcription Start Site

UN – Unchanged
UTR – Untranslated Region
VST – Variance-Stabilizing Transformation
wNES – Weighted Normalized Enrichment Score
